# Field-based dissection of stomatal anatomy and conductance reveals stable QTL under drought and heat in wheat

**DOI:** 10.64898/2026.03.30.715413

**Authors:** Chaplin Edward, Tanaka Emi, Merchant Andrew, Sznajder Beata, Trethowan Richard, Salter William

## Abstract

Stomatal traits balance carbon gain with water loss, yet their breeding potential in wheat remains underexploited. This study investigated physiological and anatomical stomatal responses alongside yield across two years of large-scale field trials under water-limitation and delayed sowing-induced heat exposure.

Across both seasons, stomatal conductance (*g*_s_) declined under stress, reflecting strong environmental constraint on gas-exchange (water-limitation:-26.9%; heat:-13.8%).

Partitioning responses by leaf surface and genotype identified the adaxial surface as the dominant contributor to *g_s_* variation and the most stress responsive. Despite increases in theoretical anatomical gas-exchange capacity (*g_smax_*), *g_s_*-efficiency declined, indicating partial decoupling between structural potential and realised conductance. Drought reduced stomatal size while increasing density whereas heat increased size, suggesting stress-specific anatomical plasticity.

Moderate-to-high heritability was observed for anatomical traits (Water-limitation: 0.13-0.57; Heat: 0.42-0.71), contrasting with lower and less stable heritability for *g_s_* (water-limitation: 0.13-0.41; heat: 0.13-0.50). Genome-wide-association-mapping identified 169 putative QTLs, predominantly for anatomical traits, including stable and co-localised pleiotropic loci. Fourteen sets of closely positioned markers were detected across seasons or studies, with stable regions on chromosomes 2B, 3B and 7B emerging as key loci. Focusing on stable loci controlling adaxial stomatal anatomy offers a realistic strategy to enhance wheat photosynthetic efficiency and climate resilience.

**Highlight:** Adaxial stomatal traits dominate gas exchange responses to heat and drought in wheat, with stable anatomical QTL identified on chromosomes 2B, 3B and 7B. Their stability across environments supports their relevance for crop improvement in water-limited and high temperature systems.

## Introduction

Wheat (*Triticum aestivum* L.) provides staple food for over one-third of the global population and underpins Australian agriculture (Erenstein *et al*., 2022; FAOSTAT, 2024). However, production is highly sensitive to water limitation and elevated temperatures, resulting in yield losses (Zampieri *et al*., 2017). Approximately 70% of global wheat production and 95% of Australian production is rainfed, with yields strongly dependent on in-season rainfall and stored soil moisture (Hochman *et al*., 2012; Dadrasi *et al*., 2023). In south-eastern Australia, projected yield declines of 9.7% by 2050 and an expansion of drought-prone regions from 15% to 60% by 2100 highlight increased production risk (Gobbett *et al*., 2017; Collins and Chenu, 2021). Concurrently, mean temperatures are projected to rise by > 1.5°C by 2040 and by 3.3-5.7°C by 2100, increasing heatwave frequency, severity, and duration (Perkins-Kirkpatrick and Lewis, 2020; IPCC, 2023; Collins *et al*., 2024). Heat and water limitation rarely occur independently; declining soil moisture restricts transpirational cooling, amplifying canopy temperature and compounding stress during critical plant developmental stages (El Habti *et al*., 2020; Bramley *et al*., 2022*a*).

The yield impacts of heat and drought are substantial. Wheat yields decline by an average of 4.1-7.8% per 1 °C increase in temperature (Zhao *et al*., 2017; Miller *et al*., 2019), and up to 20% depending on the genotype and severity of stress (Ullah *et al*., 2020). Compounding this, severe water limitation reduces yields by an average of 27.5% (Zhang *et al*., 2018), with losses of up to 50-60% under severe drought (Zhao *et al*., 2020). Although drought currently contributes the largest yield penalties during flowering, recent modelling indicates that global yield losses associated with extreme drought events may decline modestly due to climate-driven phenological shifts, whereas losses from extreme heat are projected to increase substantially approaching drought level impacts by mid-to late-century (Senapati *et al*., 2026). Collectively, these predictions highlight the need for wheat cultivars resilient to both heat and drought stress, particularly during sensitive reproductive stages. Accordingly, traits regulating plant water use and carbon assimilation - particularly those linked to stomatal function - represent priority targets for improvement.

Stomata regulate leaf gas exchange by mediating CO_2_ uptake and water use (Lawson and Leakey, 2024). High stomatal conductance (*g_s_*) supports increased photosynthesis and canopy cooling, but also increases water loss, creating a physiological trade-off central to crop water use efficiency and yield stability (Roche, 2015). Under stress, *g*ₛ is typically reduced and together with associated biochemical limitations can constrain photosynthesis and plant productivity (Chaves *et al*., 2009). Under field conditions, realised *g*_s_ (herein referred to as operational stomatal conductance or *g*_sop_) arises from the interaction between short-term, dynamically responsive stomatal aperture opening and closure, and longer-term, more fixed cellular features, including stomatal complex size and stomatal density (SD) (Lawson and Leakey, 2024). SD largely determines the maximum potential anatomical stomatal conductance (*g_smax_*) (Franks and Beerling, 2009; Ochoa *et al*., 2024). Comparing *g_smax_* with *g_sop_* provides insight into gas exchange efficiency under stress (*g_se_*; *g_se_ = g_sop_/g_smax_*) (Franks *et al*., 2009; Busch *et al*., 2024). Under water limitation, *g_s_* typically declines, reflecting a water-conserving strategy (Rahnama *et al*., 2024; Wang *et al*., 2024*a*; Redhu *et al*., 2025) whereas heat responses are more variable and strongly context-and genotype-dependent (Ramya *et al*., 2016; Mahdavi *et al*., 2021; Zhang *et al*., 2023). Under both stress types, the adaxial surface generally exhibits stronger responses (Wall *et al*., 2022; Pinto *et al*., 2025). Stomatal anatomy responds to environmental stress, with heat and drought often associated with smaller, denser stomata - particularly on the adaxial surface (Kapadiya *et al*., 2017; Lakde *et al*., 2024; Samantara *et al*., 2025) - although contrasting environments and genotypes sometimes favour larger, less dense stomata under water limitation, indicating that optimal stomatal ideotypes are strongly context dependent (Li *et al*., 2017; Dunn *et al*., 2019; Guizani *et al*., 2023).

Wheat exhibits substantial genotypic diversity in stomatal traits (Lawson and Matthews, 2020; Faralli *et al*., 2024). Extensive genotypic variation in *g_s_*, heritability as high as 73%, and previously identified QTLs highlight potential for selection (Wang *et al*., 2015; Pooja and Munjal, 2019; Ramya *et al*., 2021). However, anatomical traits tend to show higher and more stable heritability with multiple QTLs reported, some with pleiotropic effects on yield. Chromosomal regions of interest for breeding include 2B for SA, and 4B, 5B and 7A for SA and SD (Wang *et al*., 2015, 2016; Shahinnia *et al*., 2016; Liu *et al*., 2025). Yet, despite their clear relevance to rainfed adaptation, the functional expression and stability of these loci across contrasting stress environments remain poorly resolved, and it remains unclear whether similar relationships govern stomatal behaviour under water limitation.

In rainfed cropping systems characterised by episodic and unpredictable water supply, integrating anatomical capacity (*g_smax_*) with realised physiological function (*g_sop_*) is essential to understand both short-term responsiveness and season-long water-use strategies (Yan, 2021). Coupled analyses enables assessment of gas exchange efficiency and facilitates identification of shared QTLs (Estrada *et al*., 2025). Advances in high-throughput field phenotyping enable integrated assessment of stomatal anatomy and physiology at breeding-relevant scales, supporting evaluation of stomatal behaviour under realistic rainfed conditions (Gibbs and Burgess, 2024; Chaplin *et al*., 2025; Kimura *et al*., 2025).

We quantified the effects of water limitation on stomatal conductance (*g*_s_), stomatal anatomy, integrated traits (e.g., *g*_se_), and grain yield across 200 wheat genotypes in 2023. In 2024, a subset of 75 genotypes was evaluated under contrasting times of sowing to impose differential heat stress, enabling cross-season comparison of stomatal responses to water and heat. We hypothesised that both stresses would reduce operational *g*_s_, with associated increases in stomatal density and decreases in stomatal size, and that these responses would vary substantially among genotypes. Extending previous heat-focused work, this study explicitly compares stomatal anatomical and physiological responses to drought and heat in rainfed field environments, providing insight into the stability and plasticity of key stomatal traits relevant to climate-resilient breeding.

## Methods

### Plant Material, Germplasm G Experimental Conditions

Wheat (*Triticum aestivum* L.) was grown in the field at the University of Sydney I.A. Watson Grains Research Centre in Narrabri, NSW Australia (30.2743°S, 149.8093°E). Two independent field trial experiments, as part of a Heat and Drought Wheat Improvement Consortium Project (HeDWIC), were carried out over two years, one in 2023 to investigate contrasting water availability (herein referred to as ‘water limitation’ environment) and one in 2024 to study heat stress. In 2023 (Season 1; S1), 200 lines (Table S1) were established with two treatments, ‘rainfed’ and ‘irrigated’, using a randomised complete block design with three field replicates of each genotype established per treatment. In 2024 (Season 2; S2), 75 lines (Table S2) were established with two planting dates, ‘TOS 1’ and ‘TOS 2’, using a randomised complete block design with four field replicates of each genotype established per TOS.

The diverse germplasm comprised CIMMYT (International Maize and Wheat Improvement Center) and ICARDA (International Centre for Agricultural Research in the Dry Areas) heat tolerant materials imported through the CIMMYT Australia ICARDA Germplasm Evaluation (CAIGE) program. Lines include materials from SATYN (Stress Adapted Trait Yield Nursery), EDPIE (Elite Diversity International Experiment), ESWYT (Elite Selection Wheat Yield Trial), SAWYT (Semi-Arid Wheat Yield Trial), HTWYT (High Temperature Wheat Yield Trials) and the University of Sydney including recombinants from 3 cycles of genomic selection for heat tolerance (including progeny derived from crosses among CIMMYT and ICARDA materials).

Plants were sown in 12 m^2^ plots (2 x 6 m) with five planting rows at a planting density of 100 plants/m^2.^ Plots were subsequently trimmed to 8 m^2^ (2 x 4 m) before harvest. The distance between rows in each plot (row spacing) was approximately 23.5 cm and the distance between each plot was 63 cm. The predominant soil type at the field site is a black vertosol cracking clay with high water retention. For both trials, minimum tillage was used to maintain soil integrity and moisture. Field traffic was controlled as much as possible with allocated roads and pathways and GPS guidance on all machinery. Fertiliser application was kept consistent across both treatments and both trials (Urea [46% N] 100 kg/ha and Cotton Sustain [5% N, 10% P, 21% K, 1% Z] 80 kg/ha pre-planting). The experimental sites were fallowed over the summer months and rotated with a legume crop during alternate years to minimise disease outbreak and maintain soil integrity.

In S1, all plants were sown on 24.05.2023 and were harvested on 26.10.2023. Average soil moisture at planting is provided in Table S3. Irrigation was used to establish two water availability treatments to drive diversity in stomatal traits. Rainfed plants received 102.4 mm of rainfall and irrigated plants received 197.4 mm of total moisture (rainfall + irrigation) during the growing season. The mean daily minimum and maximum temperatures across the growing season were 5.8 °C and 22.5 °C, respectively. During anthesis measurement windows, mean daily minimum and maximum temperatures were 5.0 °C and 24.9 °C, respectively, and relative humidity (RH) ranged from 25.0-81.0%, with corresponding mean vapour pressure deficit (VPD) of 0.80 kPa.

In S2, TOS 1 was planted on 20.05.2024 and harvested on 05.11.2024 while TOS 2 was planted on 22.07.2024 and harvested on 27.11.2024. Table S3 details average soil moisture at planting. TOS 1 plants received 278.4 mm of rainfall and TOS 2 plants received 160.4 mm of total moisture from rainfall and irrigation. The mean daily minimum and maximum temperature across the growing season were 6.3 °C and 21.1 °C for TOS 1, respectively, and 8.7 °C and 25.0 °C for TOS 2, respectively. During measurements at anthesis, mean daily minimum and maximum temperatures were 3.3 °C and 22.3 °C at TOS 1, and 10.1 °C and 28.7 °C at TOS 2. RH ranged from 30.3-89.7% at TOS 1 and from 30.7-88.0% at TOS 2, with corresponding mean VPD of 0.59 kPa and 0.91 kPa, respectively.

### Field Measurements

In both years, measurements of stomatal physiology and anatomy were taken from fully expanded flag leaves at anthesis (Zadok stage 48-68) and chosen using a systematic randomised sampling technique with representative plants selected from the middle three planting rows of each plot, at least 50 cm from the end of each plot. Data was collected on consecutive days, and all measurements were taken between 09:30 and 14:00 to minimise diurnal effects on *g_s_* and rates of photosynthesis. In S1, one leaf was sampled from each replicate plot per treatment (n=3 per genotype per treatment) and in S2, two leaves were sampled from each replicate plot per TOS (n=8 per genotype per TOS), reflecting the implementation of a high-throughput field-phenotyping approach to increase sampling capacity (Chaplin *et al*., 2025), along with a reduced number of lines in S2.

### Stomatal Conductance

Stomatal conductance (*g_s_*) was measured in both seasons (S1 and S2) using a LI-COR LI-600 porometer/fluorometer (LI-COR Inc., Nebraska, USA), following a standardised protocol previously validated for field-grown wheat (Chaplin *et al*., 2025, Preprint). Measurements were collected from the adaxial and abaxial surfaces of the flag leaf on the main tiller.

The settings were as follows: flow rate: 150 µmol s^-1^, phase length: 300 ms, ramp amount: 25%, leaf absorptance: 0.8, fraction abs PSII: 0.5, actinic modulation rate: 500 Hz, and integrated modulation intensity: 6.67 µmol m^-2^ s^-1^, stability: medium. Measurements were taken from the central region of the leaf blade under full sunlight conditions.

### Stomatal Anatomy Image Capture

In S1 and S2, in-field microscopy was used to image stomatal anatomy on both leaf surfaces of the same flag leaf used for physiological measurements. Images were acquired using a handheld USB microscope (Dino-Lite 5MP; AnMo Electronics Corporation, Taiwan), with a 200× magnification model used in S1 (AM7515MT2A) and a 400× model in S2 (AM7515MT4A). Across seasons, imaging parameters were standardised (LED brightness = 5; axial LED = 0; resolution = 2592 × 1944 pixels).

A custom 3D-printed leaf clip with a reduced aperture was used to aid with in-field focussing and interference from external light was minimised by shielding the microscope shaft with black tape. The microscope was connected via USB to a Windows computer (Dell Latitude 7230 Rugged Extreme Tablet), and images were captured from the same leaf region previously assessed for *g_s_*.

Fine focus was achieved by adjusting clamp pressure once the leaf was secured. Images were acquired using FieldDino, a custom Python-based image capture application (PyQt5; Dino-Lite SDK), which integrates live video image feed, easily adjustable settings, and plot-linked automatic file naming. The software and installation instructions are publicly available (https://github.com/williamtsalter/FieldDinoMicroscopy) and full details are described by (Chaplin *et al*., 2025).

### Yield and Yield Components

At physiological maturity, plots were machine harvested and grain yield was expressed on a per-hectare basis. Subsamples of cleaned grain were used to quantify yield components and quality traits. Thousand kernel weight (TKW) was determined using an optical seed counter (Contador, Pfeuffer GmbH, Germany) and expressed on a dry-weight basis. Screenings (%) were assessed using a standard 2.8 mm slotted sieve. Grain protein concentration, test weight, and moisture content were measured by near-infrared spectroscopy (FOSS analytical system) according to manufacturer protocols.

## Data Analyses

### Stomatal Anatomy & Deep Learning Model

Stomatal anatomical traits were quantified using a previously developed deep-learning image analysis pipeline, described in full by Chaplin et al., (2025) and validated in wheat at field-scales by Chaplin et al., (2025, Preprint). Briefly, a subset of images were manually annotated using instance segmentation in Roboflow and partitioned into training (60%), validation (30%), and test (10%) datasets. Images were tiled (1280 × 1280 px) and augmented using rotations, flips, and contrast and saturation adjustments. Multiple YOLOv8 model architectures were trained and evaluated, with the YOLOv8-M model selected based on performance metrics (precision, recall, mean average precision) (Table S4). Automated stomatal detection and measurement were performed using custom Python scripts integrating Ultralytics and OpenCV. An ellipse was fitted to each detected stomatal complex, from which guard cell length (major axis), guard cell width (minor axis), and stomatal area were derived (Figures S1 and S2).

Maximum theoretical stomatal conductance (*g_smax_*) was calculated following Franks and Beerling (2009). The commonly assumed pore-length-to-guard-cell-length ratio was empirically evaluated, and a refined coefficient (0.543) was incorporated into *g_smax_* calculations (Franks and Farquhar, 2007; Franks and Beerling, 2009). Full computational details are provided in the associated GitHub repository (Chaplin *et al*., 2025), with methodological validation in wheat field trials provided by Chaplin et al., (2025, Preprint).

### Phenotypic Statistical & Visual Analyses

All phenotypic analyses were conducted in R (R Core Team, 2021) using packages dplyr (Wickham *et al*., 2023), ggplot2 (Wickham, 2016), gridExtra (Baptiste, 2017) and tidyr (Wickham *et al*., 2024). Linear mixed-effects models (LMM) were fitted for each trait using the ‘lmer’ function from the lme4 package to account for experimental design structure. In S1 and S2, models included Treatment (Water Limitation; S1 or Time of Sowing; S2), Genotype, and Surface as fixed effects, with Rep specified as a random effect in S1, as well as Leaf as a random effect in S2, as follows:

S1 Model:

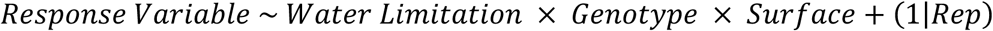

S2 Model:

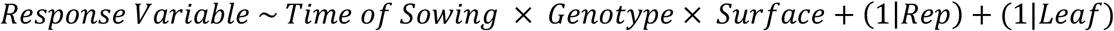

Model significance was assessed using ANOVA to test the significance of fixed effects and interactions, and post hoc comparisons were performed using the emmeans package, with Tukey’s adjustment for multiple comparisons between separate treatment groups. Packages lme4 (Bates *et al*., 2015), lmerTest (Kuznetsova *et al*., 2017), ggplot2 (Wickham, 2016), emmeans (Lenth *et al*., 2020) and MASS (Venables and Ripley, 2002) were used for statistical analyses.

Model assumptions were evaluated visually prior to fitting. Where residuals violated normality or homoscedasticity assumptions, Box–Cox transformations were applied (Box and Cox, 1964) using the MASS package, with λ selected by maximising log-likelihood across λ ∈ [−2, 2]. The most appropriate transformation was then applied (e.g., log if 𝜆 ≈ 0, square root if 𝜆 ≈ 0.5, cube root if 𝜆 ≈ 0.33) (Wilkinson and Rogers, 1973).Interpretation of TOS or treatment effects should be made cautiously, as sowing time and treatment were spatially confounded by block.

### Genome wide association mapping

For each trial, the phenotypic model was fitted to each appropriately transformed trait measured at each surface. More specifically, the phenotypic model had the form

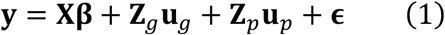

where 𝐲 is the vector of appropriately transformed observations, 𝐗 is the design matrix for fixed effects with corresponding fixed effects 𝛃 (in this case the overall mean), 𝐙_g_ and 𝐙_𝑝_are the design matrices for the random effects for genotype and experimental design factors, 𝐮_g_ and 𝐮_𝑝_, respectively, and 𝛜 is the vector of residual errors. For both seasons, the experimental design factors included the random effects for block, row, column and sampling date. For S2 trials, the measurements were made on two leaf replicates per plot, thus included a random effect for plot to account for the within-plot correlation. The random effects for genotype were assumed to be distributed as 𝐮_g_ ∼ 𝒩S𝟎, 𝜎^2^𝐈X, where 𝜎^2^ is the genotypic variance and 𝐈 is the identity matrix commensurate with the length of corresponding vector of random effects. The 𝑘-th random effects for experimental design factors were assumed to be distributed as 𝐮_𝑝_ ∼ 𝒩S𝟎, 𝜎𝑝_𝑘_^2^ 𝐈X, where 𝜎𝑝^2^ is the variance associated with the experimental design factors. The residual errors were assumed to be distributed as 𝛜 ∼ 𝒩(𝟎, 𝜎𝜖^2^𝐈), where 𝜎𝜖^2^ is the residual variance.

The phenotypic models (Equation 1) were fitted using the asreml package in R (Butler *et al*., 2023). Broad sense heritability was estimated using this model using the method described by Cullis et al. (2006) via the heritable R package (Kar *et al*., 2026). Subsequent QTL analysis was performed at a trial only if the estimated heritability was greater than 0.05.

The lines were genotyped using the Illumina Infinium Wheat Barley 40K SNP array (Keeble-Gagnère *et al*., 2021). After quality control, 35,426 SNPs were retained for the QTL analysis. In the working model, the 𝐮_g_in the phenotypic model (Equation 1) was disaggregated as 𝐮_𝑎_ + 𝐮_𝑛_, where 𝐮_𝑎_and 𝐮_𝑛_ are the additive and non-additive genetic effects, respectively. The additive and non-additive genetic effects were assumed to be distributed as 𝐮_𝑎_ ∼ 𝒩(𝟎, 𝜎𝑎^2^𝐊𝑎) and 𝐮𝑛 ∼ 𝒩(𝟎, 𝜎𝑛^2^𝐈), where 𝜎𝑎^2^ and 𝜎𝑑^2^ are the additive and non-additive genetic variances, respectively, and 𝐊_𝑎_is the genomic relationship matrix, estimated from the SNP data internally in the wgaim R package (Taylor and Verbyla, 2011; Verbyla *et al*., 2012). The QTL analysis was performed using a one-stage whole genome approach by iteratively fitting selected markers as an additive set of fixed effects in the working model in a forward selection algorithm using a family-wise error rate threshold of 0.05 and an exclusion window of 10 mega base pair (Mbp) using an approach similar to Taylor et al., (2023). The QTL analysis code is available in the data repository provided.

## Results

### Water Limitation

#### Stomatal Conductance

*g_s_* was significantly influenced by water availability, genotype, and leaf surface, as well as interactions between water availability and surface (p<0.001) (Table 1). *g_s_* was 26.9% lower under rainfed conditions compared with irrigated (p<0.001; Table S5). Mean *g_s_* under irrigated conditions were 0.082 mmol m^-2^ s^-1^ and 0.195 mmol m^-2^ s^-1^ for the abaxial and adaxial surfaces, respectively, while under rainfed conditions, mean *g_s_* were 0.062 mmol m^-2^ s^-1^ and 0.140 mmol m^-2^ s^-1^ for the two surfaces, respectively.

**Table 1:**
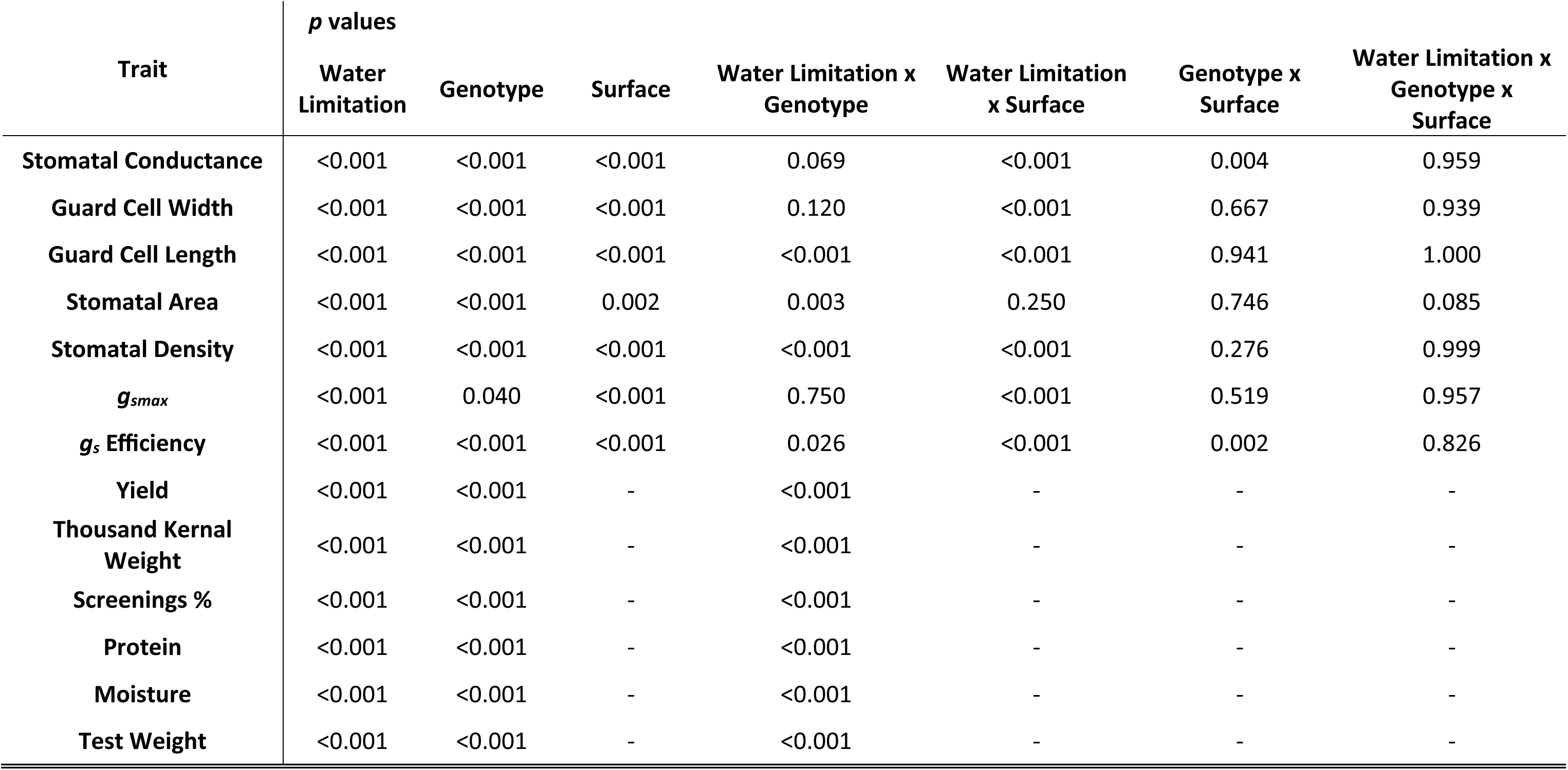
Linear mixed model ANOVA p values for Water Limitation, Genotype and Surface on traits of wheat from rainfed vs irrigated trial.

There was a significantly higher *g_s_* on the adaxial surface compared to the abaxial surface, regardless of water limitation (p<0.001). *g_s_* was highest on the adaxial surface under irrigated conditions, followed by irrigated abaxial, rainfed adaxial, and lowest on rainfed abaxial (Figure 1A). The water availability and surface interaction (p<0.001) highlighted that while water limitation reduced *g_s_* across both surfaces, this of greater significance for the adaxial surface.

**Figure 1:**
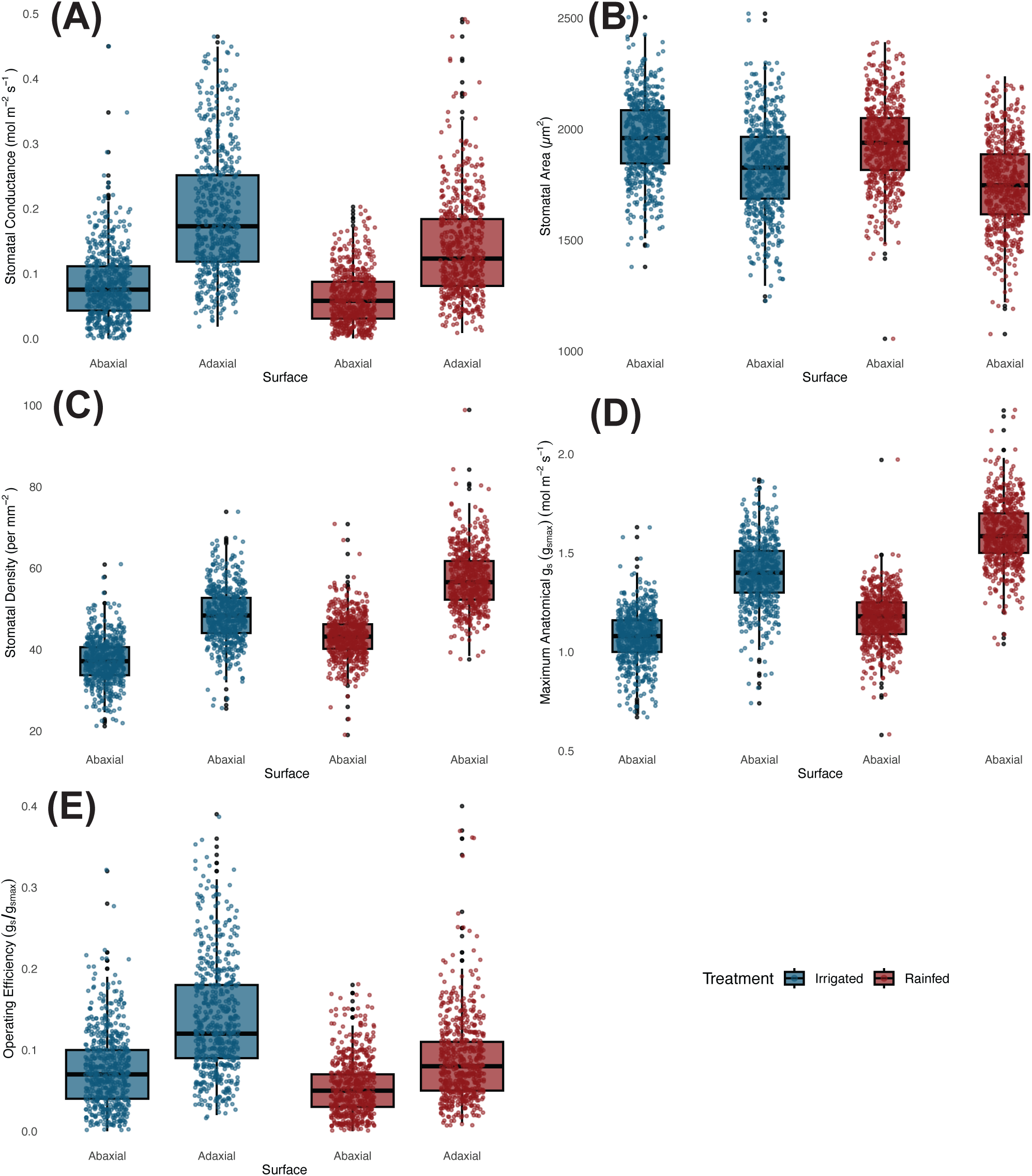
Operational stomatal conductance (*g*_sop_) and anatomical traits across 200 wheat genotypes under two watering treatments for rainfed vs irrigated trial. (a) *g*_sop_; (b) stomatal area; (c) stomatal density; (d) maximal anatomical stomatal conductance; and (e) stomatal conductance operating efficiency (*g_se_*). Boxes represent the interquartile range (25th–75th percentiles), with horizontal lines indicating the median. Whiskers denote the minimum and maximum values, and points represent individual observations.

There was also significant genotypic variability in *g_s_* across the panel (p<0.001; Figure 2). There was no water availability × genotype interaction (p=0.44), implying that the effect of water limitation was relatively consistent across genotypes. Additionally, no interactions were observed between genotype x surface (p=0.094) or in the three-way interaction between water availability x genotype x surface (p=0.95).

**Figure 2:**
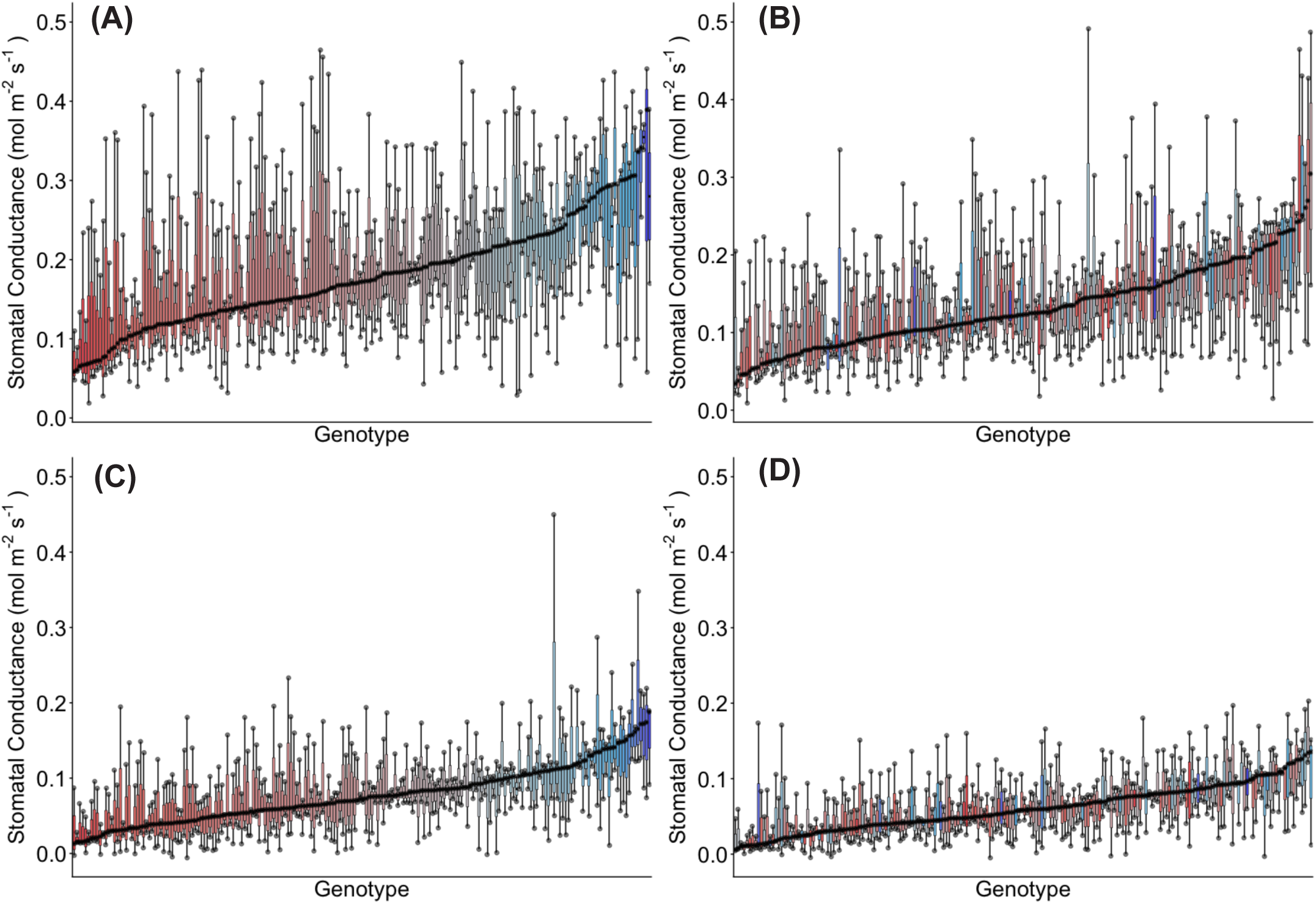
Genotypic distribution of stomatal conductance (*g_s_*) across 200 wheat genotypes under two watering treatments for rainfed vs irrigated trial. (A) Adaxial surface Irrigated; (B) adaxial surface Rainfed; (C) abaxial surface Irrigated; and (D) abaxial surface Rainfed. Genotypes are ranked by median *g_s_*. Colour assigned to each genotype based on irrigated *g_s_*. Thick horizontal lines within boxes indicate the median and boxes indicate the upper (75%) and lower (25%) quartiles. Whiskers indicate the ranges of the minimum and maximum values. Points indicate individual measurements.

### Stomatal Anatomy

Guard cell length (GCL) was significantly influenced by water limitation (p<0.001), genotype (p<0.001) and surface (p<0.001) (Table 1). For both leaf surfaces, GCL was significantly shorter under rainfed conditions than under irrigated conditions, averaging a 4.8% reduction under water limitation (p<0.001; Table S5). However, significant water availability × genotype (p<0.001) and water availability x surface (p<0.001) interactions suggest that the effect of water limitation varied dependent on the genotype and surface. There were no other interactions with respect to GCL. While there were no significant differences in GCL between the surfaces under irrigated conditions, however, the adaxial surface GCL was significantly higher under rainfed conditions (p<0.001).

Guard cell width (GCW) was also affected by water limitation (p<0.001), and there was a significant water availability x surface interaction (p<0.001), suggesting that the effect of water limitation on GCW was dependent on surface. Closer inspection of this interaction showed that abaxial GCW increased by 3.7% under rainfed conditions, while adaxial GCW was 1.7% higher under irrigation (p<0.001) (Table S5). Under both watering treatments, GCW was significantly larger on the abaxial surface (p<0.001) (Table 1). There was also significant genotypic variability in GCW across the panel (p<0.001). There were no water availability x genotype (p=0.120), genotype x surface (p=0.667) or three-way interactions (p=0.939).

For SA, a measure of stomatal size based on guard cell length and width, the effect of water limitation was significant (p<0.001) with SA declining on both surfaces by an average of 2.5% (Table S5). Significant effects of genotype (p<0.001) and surface (p<0.01) were also detected with abaxial SA consistently greater than adaxial SA (p<0.001) (Table 1; Figure 1B). The water availability x genotype interaction was significant (p<0.01), indicating that the effect of water limitation on SA varied depending on genotype. The water availability × surface, genotype × surface, and three-way Interactions were all non significant.

For SD, there were strong main effects of water limitation (p<0.001), surface (p<0.001) and genotype (p<0.001) (Table 1; Figure 1C). Significant water availability x genotype (p<0.001) and water availability x surface interactions (p<0.001) suggest that the effect of water limitation was not uniform between surfaces or across genotypes. However, the genotype x surface interaction was non significant (p=0.276; Figure 3). SD was highest on the adaxial surface regardless of water limitation (p<0.001).

**Figure 3:**
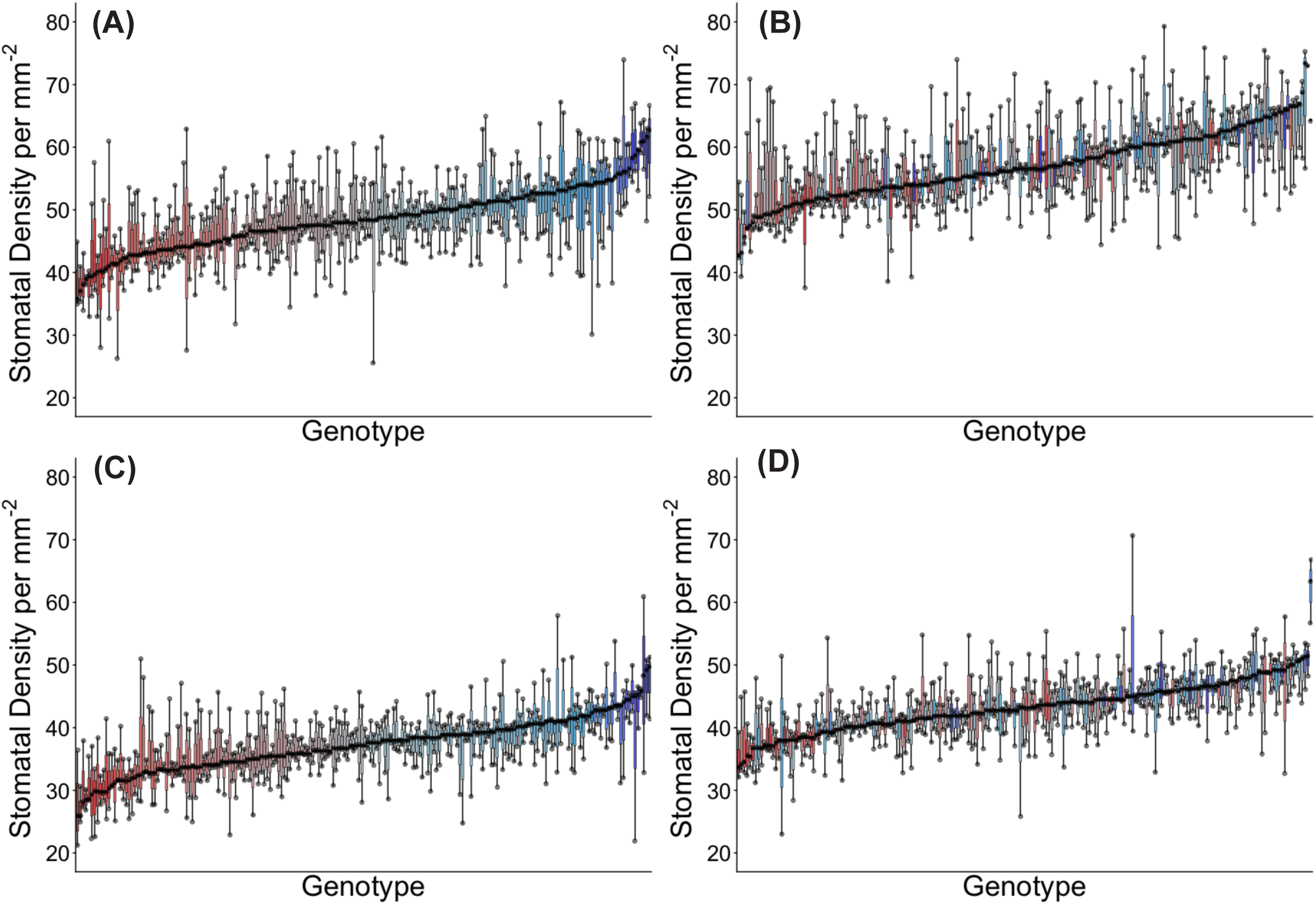
Genotypic distribution of stomatal density across 200 wheat genotypes under two watering treatments for rainfed vs irrigated trial. (A) Adaxial surface Irrigated; (B) adaxial surface Rainfed; (C) abaxial surface Irrigated; and (D) abaxial surface Rainfed. Genotypes are ranked by median stomatal density. Colour assigned to each genotype based on irrigated stomatal density. Thick horizontal lines within boxes indicate the median and boxes indicate the upper (75%) and lower (25%) quartiles. Whiskers indicate the ranges of the minimum and maximum values. Points indicate individual measurements.

SD was significantly higher under rainfed conditions on both leaf surfaces, by 17.4% on average (p<0.001; Table S5). Mean abaxial SD was 37.29 per mm^2^ and 43.21 per mm^2^ for the irrigated and rainfed treatments, respectively, while mean adaxial SD was 48.46 per mm^2^ and 57.41 per mm^2^ for the two watering treatments, respectively. SD was weakly positively correlated with *g_s_* (R^2^=0.097) and moderately negatively correlated with SA (R^2^=0.3311) (Figure S3). There was also a positive correlation between adaxial and abaxial SD.

For *g_smax_*, surface had the strongest main effect (p<0.001) while there were also significant effects of water limitation (p<0.001) and genotype (p<0.05) (Table 1; Figure 1D; Figure S4). For both surfaces, rainfed *g_smax_* was higher than irrigated *g_smax_*, by an average of 11.8% (p<0.001). Exploring the effect of surface further, adaxial *g_smax_* was significantly higher than abaxial *g_smax_*, regardless of water limitation (p<0.001). There were no significant interactions between water availability x genotype (p=0.750) or genotype x surface (p=0.519), indicating that these factors independently influenced *g_smax_*. However, a significant water availability x surface interaction was found (p<0.001), suggesting that the effect of water limitation on *g_smax_* varied depending on the surface. No three-way interaction was observed (p=0.9573). *g_smax_* showed a weak positive correlation with observed values of *g_s_* (R^2^=0.1304).

### Integrated Stomatal Traits

When *g_s_* and *g_smax_*, were considered together to determine *g_se_* (unitless), significant effects of water limitation (p<0.001), genotype (p<0.001) and surface (p<0.001) were identified (Table 1). Additionally, water availability x surface (p<0.001) and genotype x surface interactions (p<0.001) suggest that surface influences the effect of water limitation and genotype on *g_se_*. *g_se_* was significantly higher under irrigated conditions than rainfed conditions for both surfaces (p<0.001), declining by an average of 34.7% under water limitation (Figure 1E; Table S5). Under both watering treatments, adaxial *g_se_* was significantly higher than abaxial *g_se_* (p<0.001). The highest *g_se_* was for the adaxial surface under irrigated conditions (p<0.001).

### Yield Parameters

Water limitation reduced yield by an average of 21.6% under rainfed conditions (p<0.001) (Table S5). There was also significant genotypic variation (p<0.001) and the water availability x genotype interaction was significant (p<0.001) (Table 1). No significant relationships were found between yield and stomatal traits.

Thousand kernel weight (TKW) was significantly lower under rainfed conditions (p<0.001), with significant genotypic variation (p<0.001) and a significant water availability x genotype interaction (p<0.001) (Table 1). Screenings percentage was significantly higher under rainfed conditions (p<0.001) and there was also significant variation in screenings percentage across genotypes (p<0.001) (Table 1). The water availability x genotype interaction was significant (p<0.001). Protein, moisture content and test weight were all significantly affected by water limitation (p<0.001) and all varied significantly across genotypes (p<0.001) (Table 1).

### Elevated Temperatures

#### Stomatal Conductance

*g_s_* was significantly influenced by sowing time, genotype, and leaf surface, along with multiple interactions (Table 2). *g_s_* declined by an average of 13.8% under heat (delayed sowing) (p<0.001; Table S6). The significant TOS × surface interaction (p<0.01) highlights that the effect of sowing time on *g_s_* differed between the adaxial and abaxial leaf surfaces. While abaxial *g_s_* did not differ significantly between TOS 1 and TOS 2 (p=0.629), later sowing (TOS 2) significantly reduced *g_s_* on the adaxial surface (p<0.001) (Figure 4A). Mean adaxial *g_s_* were 0.311 mmol m^-2^ s^-1^ and 0.270 mmol m^-2^ s^-1^ for TOS 1 and TOS 2, respectively, while mean abaxial *g_s_* were 0.143 mmol m^-2^ s^-1^ and 0.121 mmol m^-2^ s^-1^ for TOS 1 and TOS 2, respectively.

**Figure 4:**
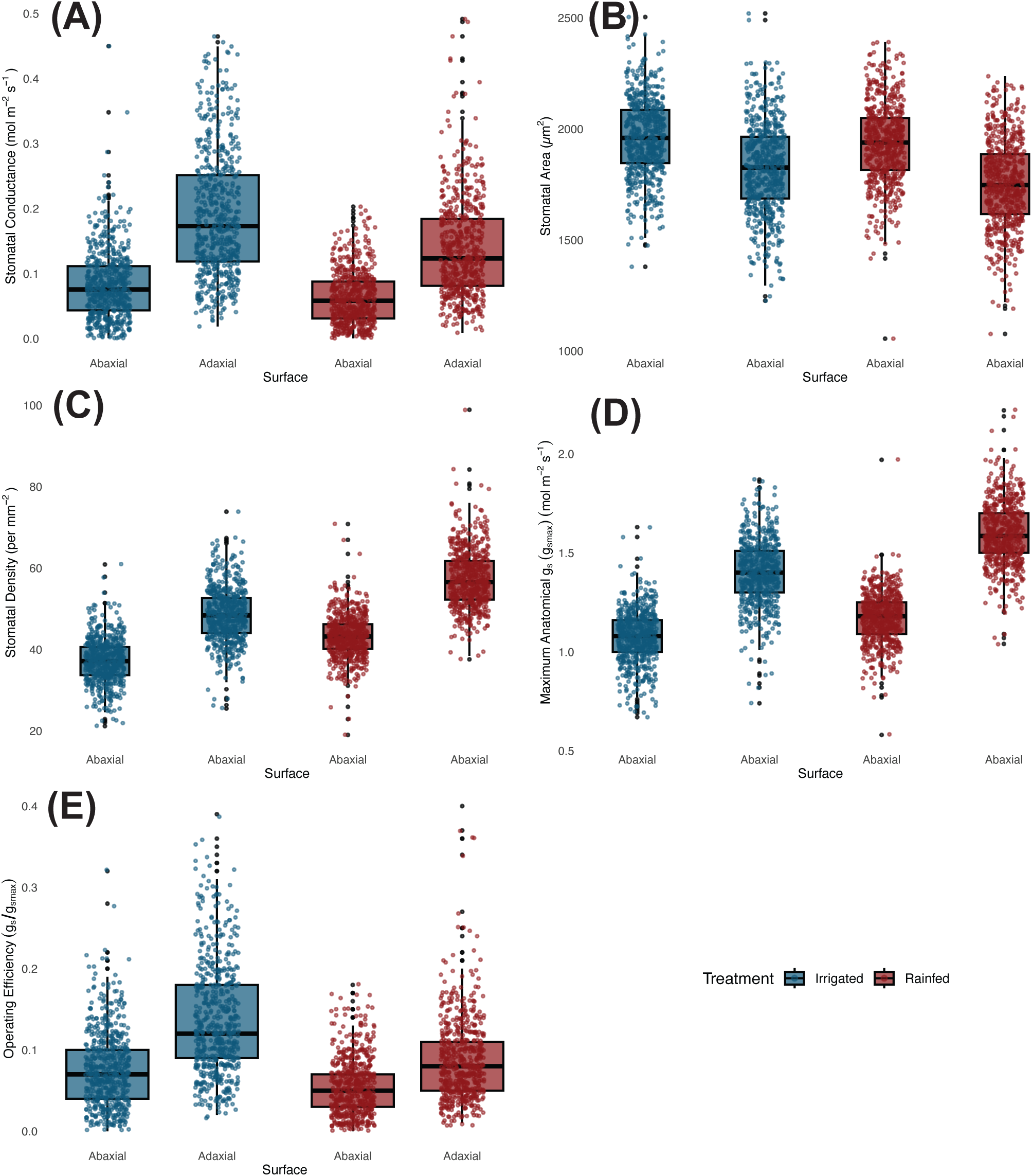
Opera-onal stomatal conductance (g_sop_) and anatomical traits across 200 wheat genotypes under two watering treatments for TOS trial. (a) gsop; (b) stomatal area; (c) stomatal density; (d) maximal anatomical stomatal conductance; and (e) stomatal conductance opera6ng efficiency (gse). Boxes represent the interquar6le range (25th–75th percen6les), with horizontal lines indica6ng the median. Whiskers denote the minimum and maximum values, and points represent individual observa6ons.

**Table 2:** Linear mixed model ANOVA p values for TOS, Genotype and Surface on traits of wheat from TOS trial.

| Trait | *p* values |  |  |  |  |  |  |
| --- | --- | --- | --- | --- | --- | --- | --- |
|  | TOS | Genotype | Surface | TOS x Genotype | TOS x Surface | Genotype x Surface | TOS x Genotype x Surface |
| **Stomatal Conductance** | <0.001 | <0.001 | <0.001 | <0.001 | 0.019 | 0.247 | 0.360 |
| **Guard Cell Width** | <0.001 | <0.001 | <0.001 | 0.277 | 0.458 | 0.988 | 1.000 |
| **Guard Cell Length** | <0.001 | <0.001 | 0.003 | 0.726 | 0.260 | 1.000 | 1.000 |
| **Stomatal Area** | <0.001 | <0.001 | <0.001 | 0.227 | 0.305 | 1.000 | 1.000 |
| **Stomatal Density** | <0.001 | <0.001 | <0.001 | 0.111 | 0.009 | <0.001 | 0.805 |
| ***g<sub>smax</sub>*** | <0.001 | <0.001 | <0.001 | 0.946 | 0.087 | 0.337 | 0.997 |
| ***g<sub>s</sub>* Efficiency** | <0.001 | <0.001 | <0.001 | <0.001 | 0.421 | 0.301 | 0.262 |
| **Yield** | <0.001 | <0.001 | - | <0.001 | - | - | - |
| **Thousand Kernal Weight** | <0.001 | <0.001 | - | <0.001 | - | - | - |
| **Screenings %** | <0.001 | <0.001 | - | <0.001 | - | - | - |
| **Protein** | <0.001 | <0.001 | - | <0.001 | - | - | - |
| **Moisture** | <0.001 | <0.001 | - | <0.001 | - | - | - |
| **Test Weight** | <0.001 | <0.001 | - | <0.001 | - | - | - |

Surface was also a significant main effect (p<0.001) with adaxial *g_s_* significantly higher than abaxial *g*ₛ across both times of sowing (p<0.001). Additionally, the largest differences in *g_s_* occurred between surfaces rather than between TOS, reinforcing the strong effect of leaf surface on *g_s_*. Significant genotypic variation in *g_s_* was also observed across the panel (p<0.001; Figure 5). A significant TOS × genotype interaction (p<0.001) suggests that the effect of sowing time on *g_s_* varied among genotypes, with some genotypes showing more pronounced *g_s_* responses under delayed sowing than others.

**Figure 5:**
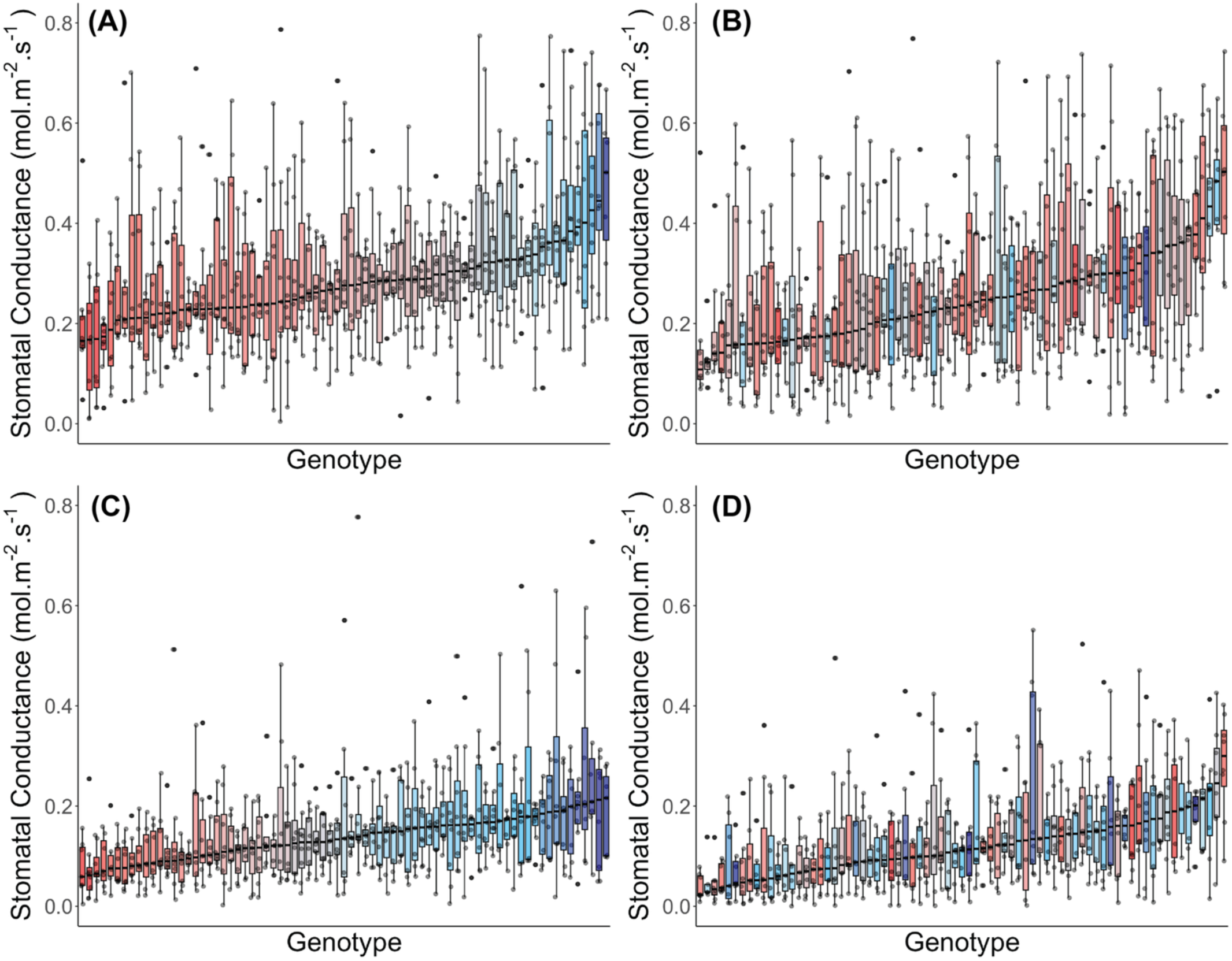
Genotypic distribution of stomatal conductance (*g_s_*) across 75 wheat genotypes at two sowing times for TOS trial. (A) Adaxial surface TOS 1; (B) adaxial surface TOS 2; (C) abaxial surface TOS 1; and (D) abaxial surface TOS 2. Genotypes are ranked by median *g_s_*. Colour assigned to each genotype based on TOS 1 *g_s_*. Thick horizontal lines within boxes indicate the median and boxes indicate the upper (75%) and lower (25%) quartiles. Whiskers indicate the ranges of the minimum and maximum values. Points indicate individual measurements.

#### Stomatal Anatomy

GCW was significantly affected by TOS (p<0.001), leaf surface (p<0.001) and genotype (p<0.001) (Table 2). GCW was on average 14.3% higher at TOS 2 compared to TOS 1 (p<0.001; Table S6). Additionally, abaxial GCW was greater than adaxial GCW, regardless of TOS (p<0.001). No significant interactions between genotype, TOS or leaf surface were identified. The lowest GCW was the adaxial surface at TOS 1 (p<0.001). The highest GCW was the abaxial surface at TOS 2 (p<0.001).

GCL was significantly higher at TOS 2 compared with TOS 1, by an average of 8.7% (p<0.001; Table 2; Table S6). Leaf surface was also a significant main effect with adaxial GCL significantly higher than abaxial GCL (p<0.05). No significant interactions between leaf surface, TOS or genotype were identified.

For SA, significant main effects of genotype (p<0.001), TOS (p<0.001) and surface (p<0.001) were identified (Table 2). Abaxial SA was significantly higher than adaxial SA across both times of sowing (p<0.001). Additionally, for both leaf surfaces, SA at TOS 2 was significantly higher than at TOS 1 (p<0.001), by an average of 17.6% (Figure 4B; Table S6).

SD was significantly affected by genotype (p<0.001), time of sowing (p<0.001) and leaf surface (p<0.001) (Table 2; Figure 6). Regardless of sowing time, adaxial SD was significantly higher than abaxial SD (p<0.001). Additionally, SD was significantly higher at TOS 2 than at TOS 1 for both surfaces (p<0.001), corresponding to an average increase of 8.8% at TOS 2 (Figure 4C) (Table S6). TOS x surface (p<0.01) and genotype x surface (p<0.001) interactions were significant, indicating that the effects of TOS and genotype on SD were dependent upon the leaf surface. No other significant interactions were found.

**Figure 6:**
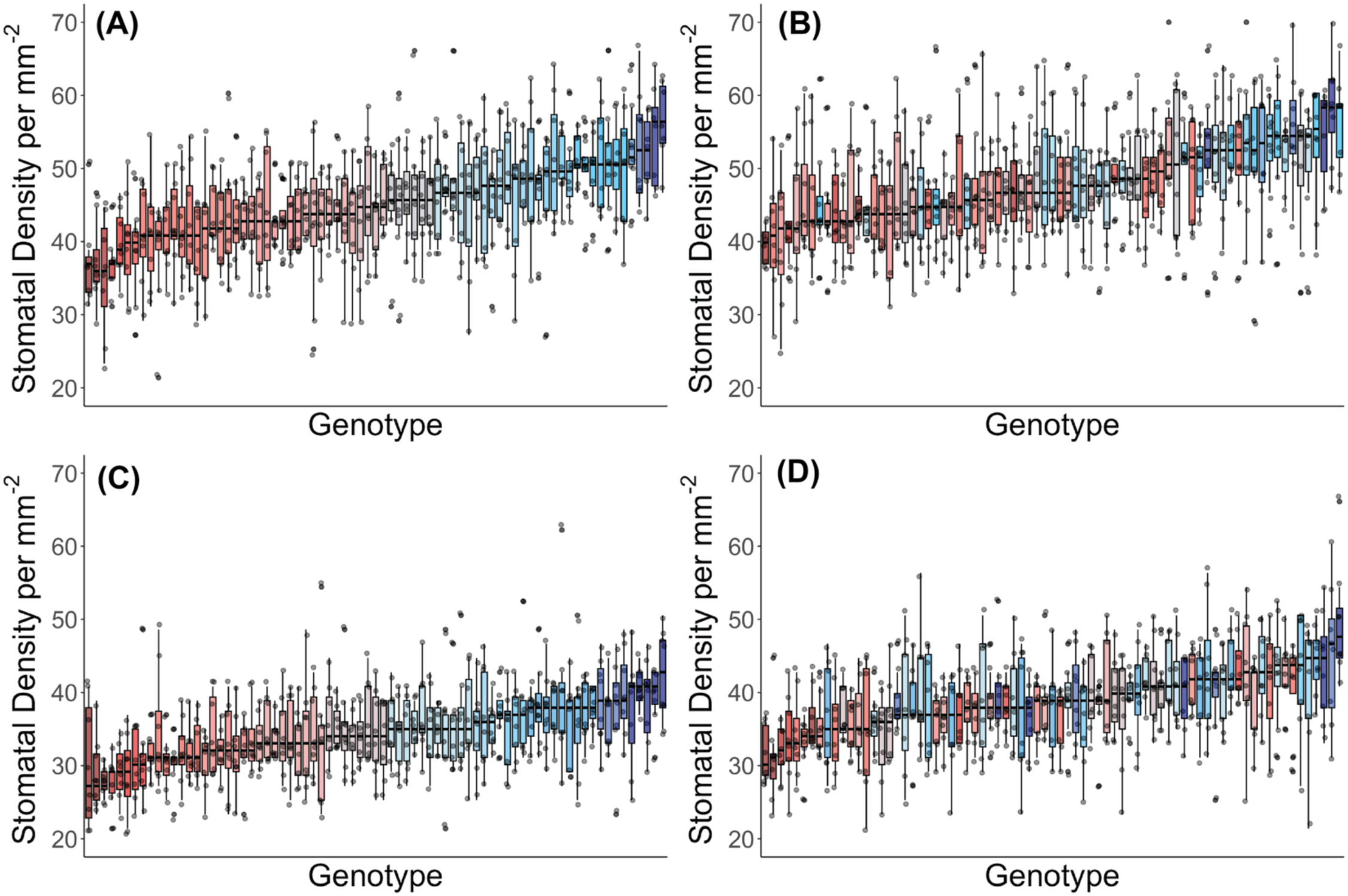
Genotypic distribution of stomatal density across 75 wheat genotypes at two sowing times for TOS trial. (A) Adaxial surface TOS 1; (B) adaxial surface TOS 2; (C) abaxial surface TOS 1; and (D) abaxial surface TOS 2. Genotypes are ranked by median stomatal density. Colour assigned to each genotype based on TOS 1 stomatal density. Thick horizontal lines within boxes indicate the median and boxes indicate the upper (75%) and lower (25%) quartiles. Whiskers indicate the ranges of the minimum and maximum values. Points indicate individual measurements.

*g_smax_* differed significantly across genotypes (p<0.001), TOS (p<0.001) and leaf surfaces (p<0.001) (Table 2; Figure 4D; Figure S5). No significant interactions were found. At both TOS, adaxial *g_smax_* was significantly higher than abaxial *g_smax_* (p<0.001) while for each leaf surface, *g_smax_* was significantly higher at TOS 2 than at TOS 1 (p<0.001), by an average of 17.5%.

#### Integrated Stomatal Traits

*g_se_* was significantly affected by TOS (p<0.001), genotype (p<0.001) and surface (p<0.001) (Table 2). Regardless of leaf surface, *g_se_* was significantly lower at TOS 2 compared with TOS 1 (p<0.001), declining by an average of 33.5% (Table S6). Additionally, abaxial *g_se_* was significantly lower than adaxial *g_se_* at both sowing times (p<0.001) (Figure 4E). There was also a significant TOS x genotype (p<0.001) interaction indicating that the effect of sowing time on *g_se_* was modulated by genotype.

#### Yield Parameters

Yield was 51.1% lower under delayed sowing at TOS 2 compared with TOS 1 (p<0.001) (Table S6) with significant genotypic variation (p<0.001) and a significant TOS x genotype interaction (p<0.001) (Table 2). No significant relationships were found between yield and stomatal traits under heat.

TKW was significantly lower at TOS 2 than at TOS 1 (p<0.001) and there was also significant genotypic variation in TKW (p<0.001) (Table 2). The TOS x genotype interaction was significant (p<0.001). Screenings percentage was 237.5% higher at TOS 2 under delayed sowing conditions than at TOS 1 under timely sown conditions (p<0.001) (Table S6), along with significant genotypic variation (p<0.001) and significant TOS x genotype interaction (p<0.001) (Table 2). Protein, moisture and test weight were all significantly affected by both genotype (p<0.001) and TOS (p<0.001) (Table 2).

#### Genome-Wide Association Mapping

All trials demonstrated moderate genetic variation for stomatal traits across seasons, sowing times and water treatments. For water availability trials, heritability for *g_s_* ranged from 0.129-0.412 and was generally higher on the abaxial surface. Heritability estimates for stomatal anatomical traits (GCL, GCW, SA, SD & *g_smax_*) were generally higher, ranging from 0.128-0.574 in S1 (Table 3). For TOS trials, heritability estimates for *g_s_* ranged from 0.131-0.496 and were again generally higher for abaxial and at TOS 2 relative to TOS 1. Anatomical traits again displayed considerably higher heritability estimates than *g_s_* across both surfaces and TOS, with estimates ranging from 0.416-0.711. *g_se_* also displayed moderate heritability, particularly at TOS 2, ranging from 0.216-0.520 (Table 3).

**Table 3:** Broad-sense heritability estimates for each trait, arranged by season, surface and treatment.

| Trait | Season 1 |  |  |  | Season 2 |  |  |  |
| --- | --- | --- | --- | --- | --- | --- | --- | --- |
|  | Abaxial |  | Adaxial |  | Abaxial |  | Adaxial |  |
|  | Rainfed | Irrigated | Rainfed | Irrigated | TOS1 | TOS2 | TOS1 | TOS2 |
|  | 2023 | 2023 | 2023 | 2023 | 2024 | 2024 | 2024 | 2024 |
| Stomatal Conductance | 0.314 | 0.412 | 0.156 | 0.129 | 0.283 | 0.496 | 0.131 | 0.496 |
| Guard Cell Width | 0.319 | 0.481 | 0.128 | 0.357 | 0.457 | 0.689 | 0.642 | 0.613 |
| Guard Cell Length | 0.518 | 0.461 | 0.574 | 0.514 | 0.687 | 0.667 | 0.688 | 0.705 |
| Stomatal Area | 0.437 | 0.525 | 0.527 | 0.530 | 0.526 | 0.707 | 0.661 | 0.674 |
| Stomatal Density | 0.505 | 0.518 | 0.458 | 0.499 | 0.637 | 0.572 | 0.711 | 0.560 |
| $g_{smax}$ | 0.462 | 0.517 | 0.384 | 0.498 | 0.416 | 0.473 | 0.629 | 0.488 |
| $g_s$ Efficiency | 0.295 | 0.408 | 0.215 | 0.111 | 0.288 | 0.520 | 0.216 | 0.464 |

QTL analyses identified a total of 169 significant marker-trait associations across stomatal traits and treatments across both trials and surfaces (Table 4). The majority of significant associations (155 markers) were observed for stomatal anatomical traits (GCW, GCL, SA, SD and *g_smax_*), particularly in water availability trials, while fewer (14 markers) were observed for stomatal physiological traits (*g_s_* and *g_se_*) (Table 4). The full list of 169 putative QTLs is available in Table S7. The significant associations were distributed across multiple chromosomes, with a higher concentration on chromosomes 3B, 6B, 1A, 2B, 5B, 7B, 1B and 6A, in order of the total markers (Table S8; Figure 7). Seven markers exhibited LOD scores greater than 10, located on chromosomes 4A (SD; LOD = 14.55), 6B (GCL; LOD = 12.48), 1A (SA; LOD = 12.27), 2B (GCL; 12.13), two at 5A (SD; LOD = 11.3 & 11.2) and 3B (SD; 11.19) (Figure S6). These high-confidence loci were predominantly associated with SD and size-related traits.

**Figure 7:**
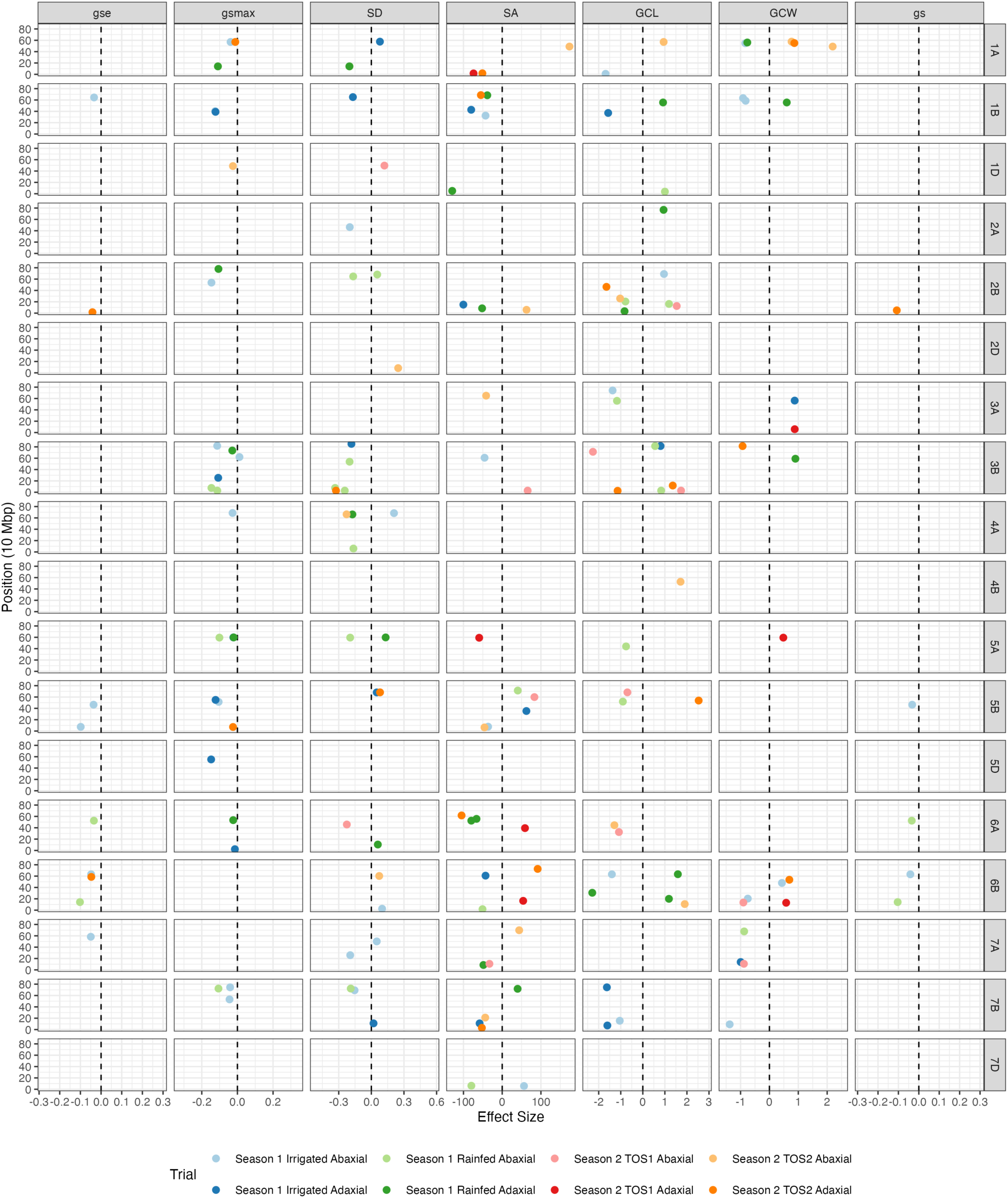
**Putative QTLs by trait and chromosome**. The black lines represent the markers (same across all traits), while the colored lines indicate putatiative QTLs identified in at least one trial, with each color corresponding to a specific season, treatment and surface. The vertical dashed lines are the y-intercepts. To avoid overplotting the positions that are co-located are adjusted using a sunflower pattern using the vayr package (Coppock, 2025).

**Table 4:** The number of putative QTL candidates by season, surface and treatment for each trait. The “.” represents 0.

| Trait | Season 1 |  |  |  | Season 2 |  |  |  |
| --- | --- | --- | --- | --- | --- | --- | --- | --- |
|  | Abaxial |  | Adaxial |  | Abaxial |  | Adaxial |  |
|  | Irrigated | Rainfed | Irrigated | Rainfed | TOS1 | TOS2 | TOS1 | TOS2 |
| Stomatal Conductance | 2 | 2 | . | . | . | . | . | 1 |
| Guard Cell Width | 6 | 1 | 2 | 3 | 2 | 2 | 3 | 3 |
| Guard Cell Length | 5 | 8 | 4 | 6 | 5 | 5 | . | 4 |
| Stomatal Area | 4 | 3 | 5 | 7 | 3 | 6 | 4 | 5 |
| Stomatal Density | 6 | 9 | 5 | 4 | 2 | 3 | . | 2 |
| $g_{smax}$ | 9 | 5 | 6 | 5 | . | 1 | . | 2 |
| $g_s$ Efficiency | 5 | 2 | . | . | . | . | . | 2 |

Multiple instances of pleiotropy were detected at co-located identical markers (Table S9). Notably, three markers on chromosome 1A associated with SD and *g*_smax_ were identified across multiple surfaces or trials, while a marker on chromosome 3B showed five associations spanning two seasons and three traits (GCL, SA and SD) (Table S9). Additional pleiotropic markers were detected on chromosomes 5A (SD and *g*_smax_ under rainfed conditions on both surfaces), 5B (*g*_s_ and *g*_se_ under irrigated conditions; GCL and SD), 6B (*g*_s_ and *g*_se_ under rainfed conditions; GCL and *g*_s_ under irrigated conditions), and 7B (SD and *g*_smax_ under rainfed conditions). The full set of pleiotropic QTLs is available in Table S9.

Several significant markers mapped to shared chromosomal regions and were therefore likely linked to the same underlying QTL. To assess clustering, we quantified significant associations within 10 Mbp windows per chromosome and identified multiple regions harbouring associations for more than one trait (Table S10). For example, chromosome 1A contained two regions with significant associations across GCW, SA, SD and *g_smax_*, while a narrow region on chromosome 3B was associated with GCL, SA, SD and *g_smax_*.

## Discussion

By integrating stomatal conductance and stomatal anatomy across two years of field trials encompassing contrasting drought and heat stress environments, this study provides new mechanistic insight into the environmental and genotypic drivers of variation in *g*_s_ and stomatal anatomy, and highlights clear opportunities to improve wheat stress tolerance under realistic field conditions.

### Stomatal Anatomical Responses to Water Limitation & Heat

Stomatal anatomical traits represent promising targets for improving heat and drought tolerance, as they are typically more heritable and under stronger genetic control than stomatal conductance itself. That being said, contrasting stress responses were observed for stomatal size, with water scarcity reducing SA whereas heat increased SA. The water limitation response aligns with established findings that smaller stomata enhance dynamic stomatal control and kinetics by enabling faster aperture responses to environmental fluctuations (Lawson and Blatt, 2014), optimising CO_2_ uptake while minimising water loss (McAusland *et al*., 2016; Faralli *et al*., 2019). In contrast, heat increased SA, consistent with some, but not all reports (Kapadiya *et al*., 2017; Pinto *et al*., 2025). Divergent responses to heat and water availability reflects distinct physiological pressures, as plants under heat stress may prioritise transpirational cooling via larger stomata with wider pores (Bramley *et al*., 2022*b*; Li *et al*., 2023), particularly during thermally sensitive reproductive stages (Ullah *et al*., 2022). Alternatively, increased size may reflect a developmental trade-off where cell expansion outpaces division, leading to fewer but larger stomata (Bertolino *et al*., 2019).

These contrasting responses highlight distinct stress-specific anatomical strategies and reinforce that the stomatal ideotype remains unknown and is likely to be environment-specific.

Consistent with previous reports, adaxial SD increased strongly under both stress conditions (Baloch *et al*., 2013; Liu *et al*., 2025). Under water limitation, reduced SA, accompanied by higher SD likely maintained gas exchange capacity (Mohi-Ud-Din *et al*., 2024; Pinto *et al*., 2025). The weak negative SA-SD correlation under rainfed conditions aligns with earlier work (Franks and Beerling, 2009; Haworth *et al*., 2023). Higher SD and smaller SA may enhance stomatal responsiveness, reducing stomatal opening/closure times from an hour to minutes (Lawson and Blatt, 2014; Raven, 2014). This developmental trade-off may be critical for dynamically balancing CO₂ uptake and water conservation, though its field-level functional consequences remain unresolved (Bertolino *et al*., 2019; Haworth *et al*., 2023).

Importantly, Figure 6 illustrates that genotypic rankings for SD remain largely conserved across TOS, a pattern also evident across water-availability treatments (Figure 3). This stability underscores that stomatal anatomical traits exhibit limited plasticity relative to physiological traits, reinforcing their potential value as reliable breeding targets.

Under both stresses, adaxial *g_smax_* exceeded abaxial *g_smax_*, aligning with previous reports (Samantara *et al*., 2025). The observation of higher *g_smax_* under stress aligns with Pinto et al., (2025), but challenges assumptions that anatomical capacity for *g_s_* declines under adverse conditions (Mohi-Ud-Din *et al*., 2024). *g_smax_* may reflect a legacy of development occurring pre-stress, or plants may upregulate anatomical capacity (via SA or SD) as a stress-avoidance strategy, particularly under heat where cooling is beneficial. The decoupling of *g*_smax_ and *g*_sop_ highlights the need to jointly consider anatomical capacity and physiological regulation, suggesting breeding should favour high gas-exchange potential coupled with dynamic *g*_s_ control.

#### QTL for Stomatal Anatomy

High heritability of stomatal anatomy across seasons supports strong genetic control reinforcing that stomatal anatomy is more genetically determined than *g_s_*. QTLs for stomatal anatomy, including pleiotropic loci linked to yield, have been reported previously (Shahinnia *et al*., 2016; Liu *et al*., 2025). We identified 155 putative anatomical QTLs, and the dominance of anatomical over physiological QTLs (155 of 169 total), further highlights their stronger genetic determinism and stability.

Comparing QTLs by chromosome and trait with published studies revealed substantial overlap (Table S11). When combined with Chaplin et al. (2025, Preprint), which examined the same germplasm across different environments, our results identified 14 clusters of two to three closely co-located QTLs for the same trait and leaf surface that were detected in multiple seasons or studies (Table S12, bolded rows; Figure 8). All markers within a set shared the same effect direction, except one set for abaxial SA in S1. Among these, eight putative QTLs for anatomical traits in our study replicated in a close position to that of the corresponding trait and surface, indicating robust, repeatedly detected loci suitable for breeding. Chromosome 2B emerged as a particularly promising region for marker-assisted-selection, consistent with earlier findings that mapped QTL to this locus (Wang *et al*., 2015, 2016; Ahmed *et al*., 2021; Liu *et al*., 2025). However, co-location with *Ppd-B1*, a photoperiod response gene, suggests phenology may indirectly influence stomatal morphology, requiring fine mapping to separate direct genetic versus developmental effects (Beales *et al*., 2007; Bentley *et al*., 2011).

**Figure 8:**
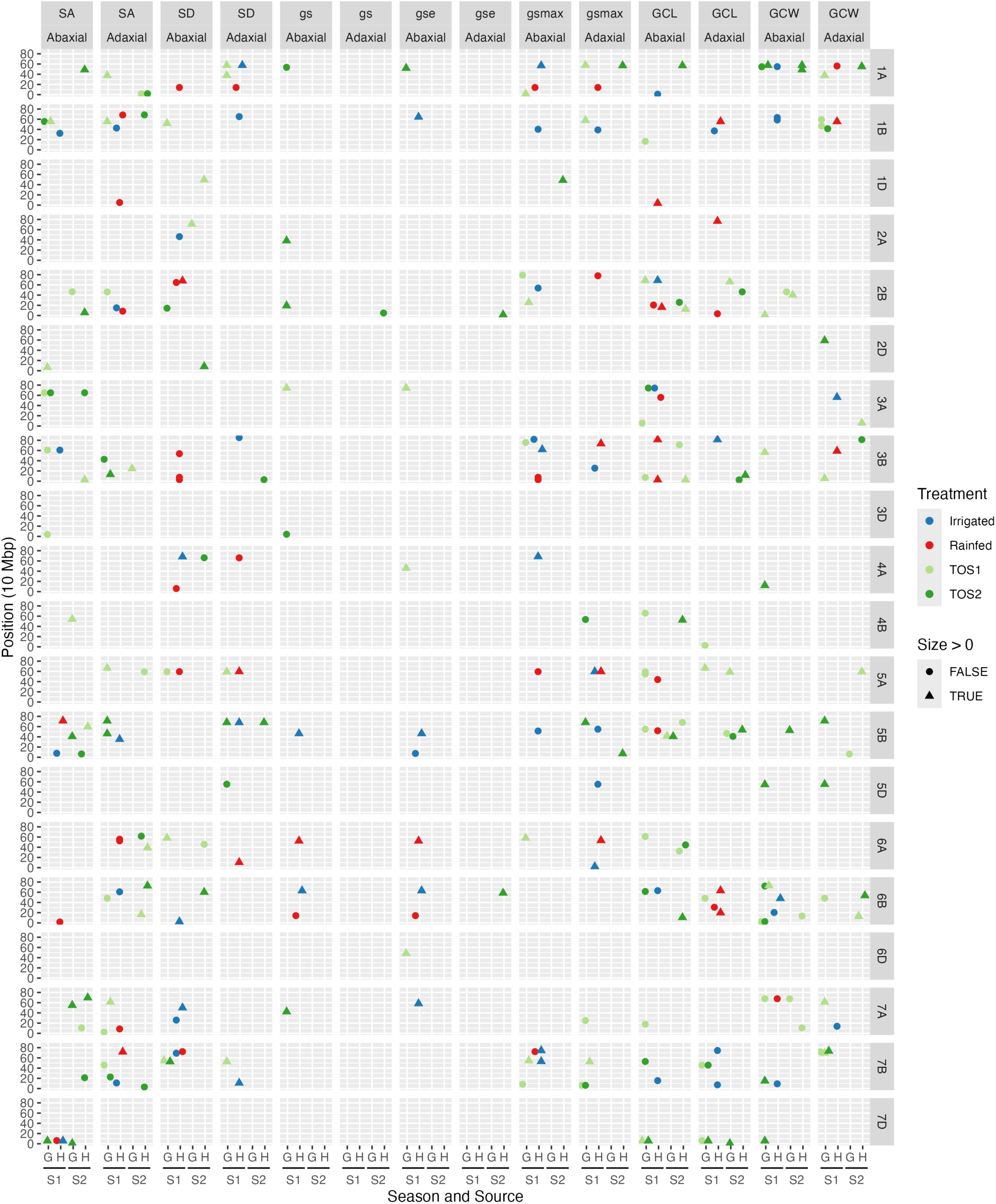
Putative QTLs across sources. The scatterplot of putative QTL candidates by position by season and source across traits, surface and chromosome. The colour indicates the treatment of the corresponding trial and the shape shows whether the effect size is greater than zero or not. The season is labelled as 1 or 2 and source is labelled as “G” if the the result is from Chaplin et al (2025), as part of a GRDC trial, otherwise “H” if the result is from this paper, as part of CIMMYT trials.

We also identified potentially stable QTL across seasons. Consistent with prior studies, chromosome 3B (∼28 Mbp) was a particularly compelling locus; a small region in this interval was associated with GCL across seasons and with SA, SD and *g_smax_*, yielding five associations (Shahinnia *et al*., 2016; Ahmed *et al*., 2021). Its high LOD score and clustering of size-and density-related traits, strongly suggest coordinated control at this locus. Chromosome 5A contained two closely positioned markers linked to adaxial *g_smax_* under both water-limited and irrigated conditions, indicating environmentally-independent control (Table S12). This stability may reflect effects of *Vrn A1* located on chromosome 5A which regulates vernalisation requirement and flowering time, potentially indirectly affecting stomatal traits (Kiss *et al*., 2014). Chromosome 7B was also prominent, with associations spanning GCL, SD, and *g*_smax_, consistent with previously reported clusters of co-located QTLs at this locus (Wang *et al*., 2015; Ahmed *et al*., 2021) (Table S10), warranting scope for further studies to validate this locus.

The identification of stable, co-localised loci across seasons supports their relevance for stomatal anatomy, while co-occurrence of multiple traits within narrow genomic regions suggests shared developmental control. Given the broad genomic distribution of associated loci, the development of large, well-characterised training populations will be critical to enable effective genomic selection for favourable stomatal traits (Meuwissen *et al*., 2001).

#### Stomatal Physiological Responses to Water Limitation & Heat

Stomatal physiological responses to water limitation and heat converge on several consistent patterns. The adaxial surface exhibited higher *g_s_* and greater stress responsiveness, highlighting it as a priority target for physiological selection. Strong genotypic variation in *g_s_* across both stress types environments aligns with previous studies (Rahnama *et al*., 2024; Wang *et al*., 2024*a*; Pantha *et al*., 2025), with stable genotypic rankings under differing moisture conditions suggesting that selection for inherently high *g*_s_ may be effective under both irrigated and moderately water-limited scenarios. In contrast, variation in thermal plasticity under heat highlights the value of selecting genotypes that maintain or appropriately adjust *g*_s_ under elevated temperatures – an increasingly important trait under unpredictable heat regimes. The lack of a genotype x water limitation interaction indicates stable genotypic rankings across moisture conditions, suggesting that selecting inherently high-*g_s_* lines may be effective under both irrigated and moderately water-limited conditions. In contrast, the significant TOS × genotype interaction demonstrates that genotypes differ in both baseline *g_s_* and their thermal plasticity, highlighting the value of selecting lines that maintain or appropriately adjust *g_s_* under heat. Taken together, these findings indicate greater genotypic variability in thermal responses than in drought responses, likely reflecting differences in phenology, development, and stress-specific physiological constraints. Such stress-responsive plasticity may be increasingly valuable under variable climates where the timing and intensity of heat waves are unpredictable.

Water limitation and heat stress regulate *g_s_* through fundamentally different physiological pathways, with hydraulic limitation dominating under water stress and VPD-and genotype-mediated responses under heat. While *g*_s_ declined under both stresses, the observed reductions are best interpreted within their respective stress contexts rather than as direct comparisons, reflecting differences in regulatory mechanisms and stress co-occurrence. The patterns observed are consistent with strong hydraulic control under water limitation and more indirect, environment-dependent regulation under heat (Rahnama *et al*., 2024; Wang *et al*., 2024*b*,*a*).

Under both stresses, the stress x surface interaction was significant, highlighting surface-specific sensitivity and non-uniform stomatal responses. The adaxial surface showed greater proportional declines. Its heightened sensitivity may reflect tighter control under evaporative demand or differential exposure to stress, though the mechanistic basis remains unclear (Wall *et al*., 2022, 2023). In contrast, abaxial stomata behaved more constitutively. This asymmetric response underscores the importance of surface specific measurements and highlights adaxial regulation as central to maintaining photosynthetic function under stress. Breeding programs may therefore benefit from prioritising adaxial specific stomatal traits rather than assuming uniform leaf level responses.

#### QTL for Stomatal Physiology

Substantial genotypic variation in *g_s_* across environments indicated genetic control of *g_s_*, though heritability was consistently lower than for anatomical traits. This is expected given *g*_s_ is highly dynamic in responsiveness to micro environmental variation. Heritability estimates reported for *g*_s_ in wheat (0.26-0.73) (Ramya *et al*., 2021) broadly overlap with the moderate estimates we observed (0.13-0.50). Despite this instability, previous studies have identified QTL for *g_s_* in wheat, some with pleiotropic loci (Wang *et al*., 2015). We detected 14 putative QTLs for stomatal physiology, but none were replicated across environments.

This contrasts sharply with the cross-season stability of anatomical QTL. Environment specific detection suggests that *g_s_* expression reflects dynamic integration of anatomical capacity, hydraulic status and environmental conditions, rather than control by a small number of robust loci. Nevertheless, pleiotropic markers linking *g_s_* and *g_se_* were identified on chromosomes 5B and 6B, suggesting partially shared genetic regulation. The signal on chromosome 5B may reflect indirect control of stomatal behaviour from phenology-related loci such as *Vrn-B1* controlling vernalisation requirement and flowering time (Kiss *et al*., 2014). However, the confinement of these loci to specific environments and leaf surfaces (irrigated abaxial for 5B, rainfed abaxial for 6B) indicates environment-dependent expression, particularly with respect to water availability.

Collectively, these findings highlight that direct selection for *g_s_* is less reliable than selection for anatomical determinants underpinning *g_smax_*. Stomatal anatomy appears governed by shared, environmentally robust genomic regions, whereas physiological traits are more variable. Anatomical loci therefore represent more stable and immediately amenable targets for breeding programs aiming to enhance climate resilience.

#### Integrated Stomatal Conductance C Anatomy Traits and QTL

The 34.7% and 33.5% reductions in *g_se_* under water limitation and higher temperatures, respectively, indicate that plants operated well below anatomical capacity, likely due to stomatal closure driven by internal water status and VPD. This inefficiency suggests that although plants were structurally capable of high gas exchange, physiological constraints limited realisation of *g_smax_*, consistent with Chaplin et al., (2025, Preprint). Thus, selection for high *g_smax_* alone is insufficient; breeders must consider how efficiently anatomical potential is realised under stress. The weak positive *g_smax_*-*g_sop_* correlation supports this and aligns with previous work showing that anatomical adjustments aim to optimise, not constrain, gas exchange (Franks *et al*., 2009; Fanourakis *et al*., 2015). This aligns with the view that smaller, denser stomata can increase *g*_smax_ and accelerate stomatal kinetics, thereby improving the efficiency of stomatal control (McAusland *et al*., 2016; Durand *et al*., 2019).

Moderate-high heritability for *g_se_* under both stresses indicates potential for selection, particularly under heat and on the abaxial surface. Nine QTLs were identified for *g_se_*, though confined to the abaxial surface under water limitation and the adaxial surface at TOS 2.

Optimising *g*_se_ remains an important breeding strategy, but our results suggest this would be more readily achieved through selection for constitutive anatomical or physiological traits rather than *g*_se_ itself. Focusing on stable traits that confer high gas-exchange capacity under stress would likely provide a more practical and robust pathway to improving climate resilience in wheat.

#### Implications for Grain Yield

Stomatal responses to stress coincided with declines in grain yield and quality (reduced TKW and increased screenings), underscoring their agronomic relevance while recognising that yield integrates multiple interacting processes. However, the optimal combination of stomatal anatomical traits for maximising yield remains unresolved, with studies reporting contrasting outcomes (Shahinnia *et al*., 2016; McAusland *et al*., 2021; Mohi-Ud-Din *et al*., 2024). This highlights the importance of elevating stomatal traits - often overlooked in breeding - while recognising that crop productivity reflects the integration of multiple processes. Effective improvement of climate resilience will require stomatal traits to be considered alongside other key limitations to productivity, including biochemical capacity for carbon fixation, light-use efficiency, respiration, source–sink dynamics, and developmental/phenological processes.

## Conclusions

By resolving surface-specific and genotype-dependent responses of stomatal conductance under field-relevant drought and heat stress, this study refines current understanding of stomatal regulation in wheat. The adaxial surface emerged as the dominant and most stress-responsive contributor to gas exchange, highlighting adaxial *g*_s_ as a biologically meaningful yet underutilised selection target. Although *g*_s_ declined under both stresses, reductions under rainfed conditions highlight the strong constraint imposed by water limitation in these environments, while genotype-and context-dependent responses indicate differential plasticity across stress types. Concurrent shifts in stomatal anatomy and *g*_smax_ demonstrate stress-specific anatomical adjustment, whereas the consistent reduction in *g*_se_ reveals a decoupling between structural capacity and realised physiological performance.

Genetically, stomatal physiology was largely environment-dependent, while anatomical traits showed stronger heritability and clearer genomic control. We identified 169 putative QTL, predominantly for anatomical traits, including stable and pleiotropic loci detected across seasons, treatments, and studies. Fourteen clusters of closely positioned QTLs, eight shared with Chaplin et al. (2025, Preprint), reinforce the robustness of several genomic regions. Together, these findings prioritise chromosomes 2B, 3B, and 7B as high-confidence targets for marker-assisted selection and demonstrate that focusing on anatomically stable, adaxial-relevant stomatal traits offers a tractable pathway to improve photosynthetic water-use efficiency and sustain yield gains under heat and water limitation, as future productivity increasingly depends on gains in photosynthetic capacity and stability (Roche, 2015).

## Supplementary Materials

**Figure S1: Representative images of stomatal anatomy collected *in situ* using 200x magnification handheld digital microscope for rainfed vs irrigated trial.** (A) adaxial and (B) abaxial raw images collected using microscope. (C) and (D) show image with automatically labelled stomata with ellipses using the deep learning model we trained.

**Figure S2: Representative images of stomatal anatomy collected *in situ* using 400x magnification handheld digital microscope for TOS trial.** (A) adaxial and (B) abaxial raw images collected using microscope. (C) and (D) show image with automatically labelled stomata with ellipses using the deep learning model we trained.

**Figure S3: Relationship between stomatal density and stomatal area for rainfed vs irrigated trial**. Shaded markers represent irrigated plants and unshaded markers represent rainfed plants. Circular markers represent abaxial leaf surface and triangular represent adaxial leaf surface.

**Figure S4: Genotypic distribution of maximum anatomical stomatal conductance, *g_smax_*, across 200 wheat genotypes under two watering treatments for rainfed vs irrigated trial.** (A) Adaxial surface irrigated; (B) adaxial surface rainfed; (C) abaxial surface irrigated; and (D) abaxial surface rainfed. Genotypes are ranked by median *g_smax_*. Colour assigned to each genotype based on irrigated *g_smax_*. Thick horizontal lines within boxes indicate the median and boxes indicate the upper (75%) and lower (25%) quartiles. Whiskers indicate the ranges of the minimum and maximum values. Points indicate individual measurements.

**Figure S5: Genotypic distribution of maximum anatomical stomatal conductance, *g_smax_*, across 75 wheat genotypes at two sowing times for TOS trial.** (A) Adaxial surface TOS 1; (B) adaxial surface TOS 2; (C) abaxial surface TOS 1; and (D) abaxial surface TOS 2. Genotypes are ranked by median *g_smax_*. Colour assigned to each genotype based on TOS 1 *g_smax_*. Thick horizontal lines within boxes indicate the median and boxes indicate the upper (75%) and lower (25%) quartiles. Whiskers indicate the ranges of the minimum and maximum values. Points indicate individual measurements.

**Figure S6: LOD score of putative QTLs by trait and chromosome**. The scatterplot shows the LOD scores of putatiative QTLs identified by chromosome and position (in mega base pairs), with each color corresponding to a specific season, treatment and surface.

**Table S1: Overview of germplasm screened in rainfed vs irrigated HeDWIC field trials.**

**Table S2: Overview of germplasm screened in TOS HeDWIC field trials. All lines grown in TOS trials were grown in rainfed vs irrigated trials.**

**Table S3: Average soil moisture content at planting for rainfed vs irrigated trial and TOS trial.**

**Table S4: Performance metrics for YOLOv8-M model.** Metrics used were: Precision, Recall, mAP50 (mean average precision at Intersection over Union = 0.5) and mAP50-95 (mean average precision at Intersection over Union from 0.5 to 0.95).

**Table S5: Rainfed vs irrigated trial data showing average values for key traits under Irrigated and Rainfed treatments, and % change from Irrigated to Rainfed.** Values are given for each surface independently and for both surfaces averaged.

**Table S6: TOS trial data showing average values for key traits at TOS 1 and at TOS 2, and % change from TOS 1 to TOS 2.** Values are given for each surface independently and for both surfaces averaged.

**Table S7: List of the putative QTL candidates for each trait, organized by trial (indexed by season and treatment) and surface.** The chromosome, position (in base pairs), effect size and LOD score for each corresponding QTL are also provided.

**Table S8: The number of distinct putative QTL candidates across all trials by trait and chromosome.** The “.” represents 0 and if the number is marked with a * then one of the QTL candidates was also detected in one other trial (*) or two other trials (**).

**Table S9: The list of pleiotropic QTL candidate markers that were identified in more than one trial or trait.**

**Table S10: The number of putative QTLs within a 10 Mbp region by chromosome and trait.** A region was clustered using complete-linkage in hierarchical clustering such that no marker in a region would have distance of more than 10Mbp. The 18 bolded rows indicate a region that may contain pleiotropic QTLs for stomatal traits.

**Table S11: The number of QTLs reported in literature by trait and chromosome.** The last column shows the number of putative QTLs we found for the same trait and chromosome.

**Table S12: A list of potential pleiotropic or stable QTL candidates that were found within a 10 Mbp region (as determined by hierarchical clustering based on complete linkage of marker position) in more than one trial or trait.** The source column indicates whether the QTL candidate was found in the Chaplin et al. (2025, preprint) (G) or this study (H). The size column shows the effect size. The row colour alternates between white and grey to easily see markers in the same range. The rows are bolded if there are more than one QTL candidate for the same trait and surface within the same range.

## Acknowledgements

We thank University of Sydney colleagues Fiona Foster and Annette Tredrea and internship student Léon Mathé (UniLaSalle, France) for supporting the field work in this study and Prof Richard Trethowan & Dr Rebecca Thistlethwaite for supplying the wheat germplasm used for method validation. We thank Dr Hannah Robinson and Dr Andrew Bowerman for the imputed genotype dataset. We acknowledge the use of the facilities, and scientific and technical assistance of the Sydney Informatics Hub and the University of Sydney Node of the Australian Plant Phenomics Network (APPN), which is supported by the Australian Government’s National Collaborative Research Infrastructure Strategy (NCRIS) and the Grains Research and Development Corporation.

We acknowledge the use of artificial intelligence (Microsoft Copilot GPT-5) in the preparation of this manuscript, solely in the final stages to enhance clarity, flow and language. We have rigorously vetted all content edited by AI tools. No content, ideas or substantive bodies of text were generated by AI, and the manuscript reflects the original ideas and work of the authors.

## Author Contributions

WS & EC: Conceptualisation; EC, ET, BS & WS: Data Curation; EC & ET: Formal Analysis; WS & RT: Funding Acquisition; EC & WS: Investigation; EC & WS: Methodology; WS: Project Administration; WS: Resources; WS: Software; AM & WS: Supervision; EC, WS & ET: Validation; EC & ET: Visualisation; EC: Writing - Original Draft; EC, AM, ET, BS, RT & WS: Writing – Review & Editing.

## Data Availability

Original datasets are available in a publicly accessible repository. The original contributions presented in the study are publicly available and the data can be found here: <u>10.5281/zenodo.19059679</u>

All files for the 3D printed leaf clip, the Python script for stomatal annotation and the files and instructions for installing and using the FieldDino App are provided in a public GitHub repository which guides users through each step – https://github.com/williamtsalter/FieldDinoMicroscopy.

## Abbreviations

CAIGE: CIMMYT Australia ICARDA Germplasm Evaluation
CIMMYT: International Maize and Wheat Improvement Center
CO_2_: Carbon Dioxide
EDPIE: Elite Diversity International Experiment
ESWYT: Elite Selection Wheat Yield Trial
GCL: Guard Cell Length
GCW: Guard Cell Width
_gs_: _Stomatal conductance_
_gse_: gsop/gsmax - Stomatal conductance operating efficiency (unitless)
_gsmax_: _Maximum anatomical stomatal conductance_
_gsop_: _Operating rate of stomatal conductance_
H_2_O: Water
HeDWIC: Heat and Drought Wheat Improvement Consortium Project
HTWYT: High Temperature Wheat Yield Trials
ICARDA: International Centre for Agricultural Research in the Dry Areas
IRGA: Infra-red gas analysers
mAP: Mean average precision
Mbp: Mega base pair
LOD: Logarithm of the odds
PSII: Photosystem II
QTL: Quantitative Trait Loci
RH: Relative Humidity
SATYN: Stress Adapted Trait Yield Nursery
SAWYT: Semi-Arid Wheat Yield Trial
S1: Season 1; 2023
S2: Season 2; 2024
SA: Stomatal Area
SD: Stomatal density (stomata m^-2^)
TKW: Thousand Kernel Weight
TOS: Time of Sowing
VPD: Vapour Pressure Deficit
WUE: Water Use Efficiency
YOLO: You Only Look Once

## References

Ahmed HGM-D, Iqbal MN, Iqbal MA, et al. 2021. Genome-Wide Association Mapping for Stomata and Yield Indices in Bread Wheat under Water Limited Conditions. Agronomy 11, 1646.

Baloch MJ, Dunwell J, Khan DrN, Jatoi W, Khakhwani A, Vessar N, Gul S. 2013. Morpho-physiological Characterization of Spring Wheat Genotypes under Drought Stress. International Journal of Agriculture and Biology 15, 945–950.

Baptiste A. 2017. gridExtra: Miscellaneous Functions for ‘Grid’ Graphics.

Bates D, Maechler M, Bolker B, Walker S. 2015. lme4: Linear Mixed-Effects Models using ‘Eigen’ and S4.

Beales J, Turner A, Griffiths S, Snape JW, Laurie DA. 2007. A pseudo-response regulator is misexpressed in the photoperiod insensitive Ppd-D1a mutant of wheat (Triticum aestivum L.). TAG. Theoretical and applied genetics. Theoretische und angewandte Genetik 115, 721– 733.

Bentley AR, Turner AS, Gosman N, Leigh FJ, Maccaferri M, Dreisigacker S, Greenland A, Laurie DA. 2011. Frequency of photoperiod-insensitive Ppd-A1a alleles in tetraploid, hexaploid and synthetic hexaploid wheat germplasm. in press.

Bertolino LT, Caine RS, Gray JE. 2019. Impact of Stomatal Density and Morphology on Water-Use Efficiency in a Changing World. Frontiers in Plant Science 10.

Box GEP, Cox DR. 1964. An Analysis of Transformations. Journal of the Royal Statistical Society: Series B (Methodological) 26, 211–243.

Bramley H, Ranawana SRWMCJK, Palta JA, Stefanova K, Siddique KHM. 2022a. Transpirational Leaf Cooling Effect Did Not Contribute Equally to Biomass Retention in Wheat Genotypes under High Temperature. Plants 11.

Bramley H, Ranawana SRWMCJK, Palta JA, Stefanova K, Siddique KHM. 2022b. Transpirational Leaf Cooling Effect Did Not Contribute Equally to Biomass Retention in Wheat Genotypes under High Temperature. Plants 11, 2174.

Busch FA, Ainsworth EA, Amtmann A, et al. 2024. A guide to photosynthetic gas exchange measurements: Fundamental principles, best practice and potential pitfalls. Plant, Cell & Environment n/a.

Butler DG, Cullis BR, Gilmour AR, Gogel BJ, Thompson R. 2023. ASReml - ASReml estimates variance components under a general linear mixed model by residual maximum likelihood (REML). VSN International Ltd. 2 Amberside House, Wood Lane, Hemel Hempstead, HP2 4TP, UK.

Chaplin E, Coleman G, Merchant A, Salter W. 2025. FieldDino: Rapid In-Field Stomatal Anatomy and Physiology Phenotyping. Plant, Cell & Environment doi: 10.1111/pce.15639.

Chaplin ED, Tanaka E, Merchant A, Sznajder B, Trethowan R, Salter WT. 2025. QTL for Heat-Induced Stomatal Anatomy Underpin Gas Exchange Variation in Field-Grown Wheat. bioRxiv.

Chaves MM, Flexas J, Pinheiro C. 2009. Photosynthesis under drought and salt stress: regulation mechanisms from whole plant to cell. Annals of Botany 103, 551–560.

Collins M, Beverley JD, Bracegirdle TJ, et al. 2024. Emerging signals of climate change from the equator to the poles: new insights into a warming world. Frontiers in Science 2.

Collins B, Chenu K. 2021. Improving productivity of Australian wheat by adapting sowing date and genotype phenology to future climate. Climate Risk Management 32, 100300.

Coppock A. 2025. vayr: Extensions for ‘ggplot2’ to Visualize as You Randomize.

Cullis BR, Smith AB, Coombes NE. 2006. On the design of early generation variety trials with correlated data. Journal of Agricultural, Biological, and Environmental Statistics 11, 381–393.

Dadrasi A, Chaichi M, Nehbandani A, Soltani E, Nemati A, Salmani F, Heydari M, Yousefi AR. 2023. Global insight into understanding wheat yield and production through Agro-Ecological Zoning. Scientific Reports 13, 15898.

Dunn J, Hunt L, Afsharinafar M, Meselmani MA, Mitchell A, Howells R, Wallington E, Fleming AJ, Gray JE. 2019. Reduced stomatal density in bread wheat leads to increased water-use efficiency. Journal of Experimental Botany 70, 4737–4748.

Durand M, Brendel O, Buré C, Le Thiec D. 2019. Altered stomatal dynamics induced by changes in irradiance and vapour-pressure deficit under drought: impacts on the whole-plant transpiration efficiency of poplar genotypes. New Phytologist 222, 1789–1802.

El Habti A, Fleury D, Jewell N, Garnett T, Tricker PJ. 2020. Tolerance of Combined Drought and Heat Stress Is Associated With Transpiration Maintenance and Water Soluble Carbohydrates in Wheat Grains. Frontiers in Plant Science 11.

Erenstein O, Jaleta M, Mottaleb KA, Sonder K, Donovan J, Braun H-J. 2022. Global Trends in Wheat Production, Consumption and Trade. In: Reynolds MP, Braun H-J, eds. Wheat Improvement: Food Security in a Changing Climate. Cham: Springer International Publishing, 47–66.

Estrada F, Gonzàlez-Meler MA, Dias de Oliveira EA, del Pozo A, Lobos GA. 2025. Morphophysiological Plant Phenotyping for the Development of Plant Breeding Under Drought and Heat Conditions: A Practical Approach. Food and Energy Security 14, e70030.

Fanourakis D, Giday H, Milla R, Pieruschka R, Kjaer KH, Bolger M, Vasilevski A, Nunes-Nesi A, Fiorani F, Ottosen C-O. 2015. Pore size regulates operating stomatal conductance, while stomatal densities drive the partitioning of conductance between leaf sides. Annals of Botany 115, 555–565.

FAOSTAT. 2024. Food and Agriculture Organization of the United Nations, Statistics Division - Wheat production for all countries - 2023. https://www.fao.org/faostat/en/#data/QCL. Accessed February 2024.

Faralli M, Cockram J, Ober E, Wall S, Galle A, Van Rie J, Raines C, Lawson T. 2019. Genotypic, Developmental and Environmental Effects on the Rapidity of gs in Wheat: Impacts on Carbon Gain and Water-Use Efficiency. Frontiers in Plant Science 10.

Faralli M, Mellers G, Wall S, et al. 2024. Exploring natural genetic diversity in a bread wheat multi-founder population: Dual imaging of photosynthesis and stomatal kinetics. Journal of Experimental Botany, 233.

Franks PJ, Beerling DJ. 2009. Maximum leaf conductance driven by CO2 effects on stomatal size and density over geologic time. Proceedings of the National Academy of Sciences 106, 10343–10347.

Franks PJ, Drake PL, Beerling DJ. 2009. Plasticity in maximum stomatal conductance constrained by negative correlation between stomatal size and density: an analysis using Eucalyptus globulus. Plant, cell & environment 32, 1737–1748.

Franks PJ, Farquhar GD. 2007. The mechanical diversity of stomata and its significance in gas-exchange control. Plant Physiology 143, 78–87.

Gibbs JA, Burgess AJ. 2024. Application of deep learning for the analysis of stomata: A review of current methods and future directions. Journal of Experimental Botany, erae207.

Gobbett DL, Hochman Z, Horan H, Garcia JN, Grassini P, Cassman KG. 2017. Yield gap analysis of rainfed wheat demonstrates local to global relevance. The Journal of Agricultural Science 155, 282–299.

Guizani A, Askri H, Amenta ML, Defez R, Babay E, Bianco C, Rapaná N, Finetti-Sialer M, Gharbi F. 2023. Drought responsiveness in six wheat genotypes: identification of stress resistance indicators. Frontiers in Plant Science 14, 1232583.

Haworth M, Marino G, Materassi A, Raschi A, Scutt CP, Centritto M. 2023. The functional significance of the stomatal size to density relationship: Interaction with atmospheric [CO2] and role in plant physiological behaviour. Science of The Total Environment 863, 160908.

Hochman Z, Gobbett D, Holzworth D, McClelland T, van Rees H, Marinoni O, Garcia JN, Horan H. 2012. Quantifying yield gaps in rainfed cropping systems: A case study of wheat in Australia. Field Crops Research 136, 85–96.

IPCC. 2023. Climate Change 2023: Synthesis Report. Contribution of Working Groups I, II and III to the Sixth Assessment Report of the Intergovernmental Panel on Climate Change. Geneva, Switzerland: IPCC: Intergovernmental Panel on Climate Change.

Kapadiya K, Singh C, Bhalara R, Kandoliya U, Dabhi K. 2017. Effect of higher temperature on leaf anatomy of heat tolerance and heat susceptible wheat genotypes (Triticum aestivum L.) by scanning electron microscopy. 6, 2270–2277.

Kar F, Deng Y, Tanaka E. 2026. heritable: Heritability Estimation from Mixed Models.

Keeble-Gagnère G, Pasam R, Forrest KL, et al. 2021. Novel Design of Imputation-Enabled SNP Arrays for Breeding and Research Applications Supporting Multi-Species Hybridization. Frontiers in Plant Science 12.

Kimura K, Fushimi E, Kumagai E, Nomura K, Matsunami T, Konno S, Maruyama A. 2025. Estimating Leaf CO2 Assimilation in C3 Plants Using a Handheld Porometer With Chlorophyll Fluorometer in Field Conditions. Plant, Cell & Environment 48, 7213–7224.

Kiss T, Balla K, Veisz O, Láng L, Bedő Z, Griffiths S, Isaac P, Karsai I. 2014. Allele frequencies in the VRN-A1, VRN-B1 and VRN-D1 vernalization response and PPD-B1 and PPD-D1 photoperiod sensitivity genes, and their effects on heading in a diverse set of wheat cultivars (Triticum aestivum L.). Molecular Breeding 34, 297–310.

Kuznetsova A, Brockhoff PB, Christensen RHB. 2017. lmerTest Package: Tests in Linear Mixed Effects Models. Journal of Statistical Software 82, 1–26.

Lakde S, Khobra R, Sahi VP, Mamrutha HM, Wadhwa Z, Rani P, Kumar Y, Ahlawat OP, Singh G. 2024. Unraveling the ability of wheat to endure drought stress by analyzing physio-biochemical, stomatal and root architectural traits. Plant Physiology Reports doi: 10.1007/s40502-024-00799-z.

Lawson T, Blatt MR. 2014. Stomatal size, speed, and responsiveness impact on photosynthesis and water use efficiency. Plant Physiology 164, 1556–1570.

Lawson T, Leakey ADB. 2024. Stomata: custodians of leaf gaseous exchange. Journal of Experimental Botany 75, 6677–6682.

Lawson T, Matthews J. 2020. Guard Cell Metabolism and Stomatal Function. Annual Review of Plant Biology 71, 273–302.

Lenth R, Banfai B, Bolker B, Buerkner P. 2020. emmeans: Estimated Marginal Means, aka Least-Squares Means.

Li Q, Gao Y, Hamani AKM, Fu Y, Liu J, Wang H, Wang X. 2023. Effects of Warming and Drought Stress on the Coupling of Photosynthesis and Transpiration in Winter Wheat (Triticum aestivum L.). Applied Sciences 13, 2759.

Li Y, Li H, Li Y, Zhang S. 2017. Improving water-use efficiency by decreasing stomatal conductance and transpiration rate to maintain higher ear photosynthetic rate in drought-resistant wheat. The Crop Journal 5, 231–239.

Liu D, Lu S, Tian R, Zhang X, Dong Q, Ren H, Chen L, Hu Y-G. 2025. Mining genomic regions associated with stomatal traits and their candidate genes in bread wheat through genome-wide association study (GWAS). Theoretical and Applied Genetics 138, 20.

Mahdavi S, Arzani A, Maibody SAMM, Mehrabi AA. 2021. Photosynthetic and yield performance of wheat (Triticum aestivum L.) under sowing in hot environment. Acta Physiologiae Plantarum 43, 106.

McAusland L, Smith KE, Williams A, Molero G, Murchie EH. 2021. Nocturnal stomatal conductance in wheat is growth-stage specific and shows genotypic variation. New Phytologist 232, 162–175.

McAusland L, Vialet-Chabrand S, Davey P, Baker NR, Brendel O, Lawson T. 2016. Effects of kinetics of light-induced stomatal responses on photosynthesis and water-use efficiency. New Phytologist 211, 1209–1220.

Meuwissen TH, Hayes BJ, Goddard ME. 2001. Prediction of total genetic value using genome-wide dense marker maps. Genetics 157, 1819–1829.

Miller O, Helman D, Svoray T, Morin E, Bonfil DJ. 2019. Explicit wheat production model adjusted for semi-arid environments. Field Crops Research 231, 93–104.

Mohi-Ud-Din M, Hossain MA, Rohman MM, Uddin MN, Haque MS, Tahery MH, Hasanuzzaman M. 2024. Multi-Trait Index-Based Selection of Drought Tolerant Wheat: Physiological and Biochemical Profiling. Plants 14, 35.

Ochoa ME, Henry C, John GP, Medeiros CD, Pan R, Scoffoni C, Buckley TN, Sack L. 2024. Pinpointing the causal influences of stomatal anatomy and behavior on minimum, operational, and maximum leaf surface conductance. Plant Physiology 196, kiae292.

Pantha S, Kilian B, Özkan H, Zeibig F, Frei M. 2025. A comparison of drought responses in wild wheat relatives and domesticated wheat grown under irrigated and rainfed field conditions. Field Crops Research 321, 109678.

Perkins-Kirkpatrick SE, Lewis SC. 2020. Increasing trends in regional heatwaves. Nature Communications 11, 3357.

Pinto RS, Garatuza-Payán J, Murchie EH, Reynolds MP, Yepez EA. 2025. Wheat genotypes selected for their high early daytime stomatal conductance under elevated nocturnal temperatures maintain high yield and biomass. AoB PLANTS, plae072.

Pooja, Munjal R. 2019. Heat tolerant wheat genotypes for late sown conditions identified on the basis of physiological traits. Journal of Agrometeorology 21, 97–100.

R Core Team. 2021. A language and environment for statistical computing. https://www.r-project.org/.

Rahnama A, Hosseinalipour B, Farrokhian Firouzi A, Tom Harrison M, Ghorbanpour M. 2024. Root architecture traits and genotypic responses of wheat at seedling stage to water-deficit stress. Cereal Research Communications 52, 1499–1510.

Ramya KT, Bellundagi A, Harikrishna, Rai N, Jain N, Singh PK, Arora A, Singh GP, Prabhu KV. 2021. Gene Action Governing the Inheritance of Stomatal Conductance in Four Wheat Crosses Under High Temperature Stress Condition. Frontiers in Plant Science 12.

Ramya KT, Jain N, Amasiddha B, Singh PK, Arora A, Singh GP, Prabhu KV. 2016. Genotypic response for stomatal conductance due to terminal heat stress under late sown condition in wheat (Triticum aestivum L.). Indian Journal of Genetics and Plant Breeding 76, 255–265.

Raven JA. 2014. Speedy small stomata? Journal of Experimental Botany 65, 1415–1424.

Redhu M, Singh V, Kumari A, et al. 2025. Evaluation of physio-biochemical traits in bread wheat RILs for terminal heat stress. Cereal Research Communications 53, 1401–1412.

Roche D. 2015. Stomatal Conductance Is Essential for Higher Yield Potential of C3 Crops. Critical Reviews in Plant Sciences 34, 429–453.

Samantara K, Ivandi E, Tulva I, et al. 2025. Higher adaxial stomatal density is associated with lower grain yield in spring wheat. New Phytologist n/a.

Senapati N, Halford NG, Hawkesford MJ, Shewry PR, Semenov MA. 2026. Extreme heat and drought at flowering could threaten global wheat yields under climate change. Climatic Change 179, 28.

Shahinnia F, Le Roy J, Laborde B, Sznajder B, Kalambettu P, Mahjourimajd S, Tilbrook J, Fleury D. 2016. Genetic association of stomatal traits and yield in wheat grown in low rainfall environments. BMC Plant Biology 16, 150.

Taylor J, Jorgensen D, Moffat CS, Chalmers KJ, Fox R, Hollaway GJ, Cook MJ, Neate SM, See PT, Shankar M. 2023. An international wheat diversity panel reveals novel sources of genetic resistance to tan spot in Australia. TAG. Theoretical and applied genetics. Theoretische und angewandte Genetik 136, 61.

Taylor J, Verbyla A. 2011. R Package wgaim: QTL Analysis in Bi-Parental Populations Using Linear Mixed Models. Journal of Statistical Software 40, 1–18.

Ullah S, Bramley H, Mahmood T, Trethowan R. 2020. A strategy of ideotype development for heat-tolerant wheat. Journal of Agronomy and Crop Science 206, 229–241.

Ullah A, Nadeem F, Nawaz A, Siddique KHM, Farooq M. 2022. Heat stress effects on the reproductive physiology and yield of wheat. Journal of Agronomy and Crop Science 208, 1– 17.

Venables W, Ripley B. 2002. Modern Applied Statistics with S. Springer, New York.

Verbyla AP, Taylor JD, Verbyla KL. 2012. RWGAIM: an efficient high-dimensional random whole genome average (QTL) interval mapping approach. Genetics Research 94, 291–306.

Wall S, Cockram J, Vialet-Chabrand S, Van Rie J, Gallé A, Lawson T. 2023. The impact of growth at elevated [CO2] on stomatal anatomy and behavior differs between wheat species and cultivars. Journal of Experimental Botany 74, 2860–2874.

Wall S, Vialet-Chabrand S, Davey P, Van Rie J, Galle A, Cockram J, Lawson T. 2022. Stomata on the abaxial and adaxial leaf surfaces contribute differently to leaf gas exchange and photosynthesis in wheat. New Phytologist 235, 1743–1756.

Wang S, Jia S, Sun D, Fan H, Chang X, Jing R. 2016. Mapping QTLs for stomatal density and size under drought stress in wheat (Triticum aestivum L.). Journal of Integrative Agriculture 15, 1955–1967.

Wang SG, Jia SS, Sun DZ, Wang HY, Dong FF, Ma HX, Jing RL, Ma G. 2015. Genetic basis of traits related to stomatal conductance in wheat cultivars in response to drought stress. Photosynthetica 53, 299–305.

Wang Q, Wu Y, Ozavize SF, Qiu C-W, Holford P, Wu F. 2024a. Genotypic Differences in Morphological, Physiological and Agronomic Traits in Wheat (Triticum aestivum L.) in Response to Drought. Plants 13, 307.

Wang L, Zhang Y, Luo D, Hu X, Feng P, Mo Y, Li H, Gong S. 2024b. Integrated Effects of Soil Moisture on Wheat Hydraulic Properties and Stomatal Regulation. Plants 13, 2263.

Wickham H. 2016. ggplot2: Elegant Graphics for Data Analysis. Springer International Publishing.

Wickham H, François R, Henry L, Müller K, Vaughan D. 2023. dplyr: A grammar of data manipulation.

Wickham H, Vaughan D, Maximillian G. 2024. Tidyr: Tidy messy data.

Wilkinson GN, Rogers CE. 1973. Symbolic Description of Factorial Models for Analysis of Variance. Journal of the Royal Statistical Society. Series C (Applied Statistics) 22, 392–399.

Yan W. 2021. A Systematic Narration of Some Key Concepts and Procedures in Plant Breeding. Frontiers in Plant Science 12.

Zampieri M, Ceglar A, Dentener F, Toreti A. 2017. Wheat yield loss attributable to heat waves, drought and water excess at the global, national and subnational scales. Environmental Research Letters 12, 064008.

Zhang X, Ma M, Wu C, Huang S, Danish S. 2023. Mitigation of heat stress in wheat (Triticum aestivum L.) via regulation of physiological attributes using sodium nitroprusside and gibberellic acid. BMC Plant Biology 23, 302.

Zhang J, Zhang S, Cheng M, Jiang H, Zhang X, Peng C, Lu X, Zhang M, Jin J. 2018. Effect of drought on agronomic traits of rice and wheat: A meta-analysis. International journal of environmental research and public health 15, 839.

Zhao C, Liu B, Piao S, Wang X, Lobell DB, Huang Y, Huang M, Yao Y, Bassu S, Ciais P. 2017. Temperature increase reduces global yields of major crops in four independent estimates. Proceedings of the National Academy of sciences 114, 9326–9331.

Zhao W, Liu L, Shen Q, Yang J, Han X, Tian F, Wu J. 2020. Effects of Water Stress on Photosynthesis, Yield, and Water Use Efficiency in Winter Wheat. Water 12, 2127.

