## Supplementary Tables and Figures for "Field-based dissection of stomatal anatomy and conductance reveals stable QTL under drought and heat in wheat"

### Supplementary Materials

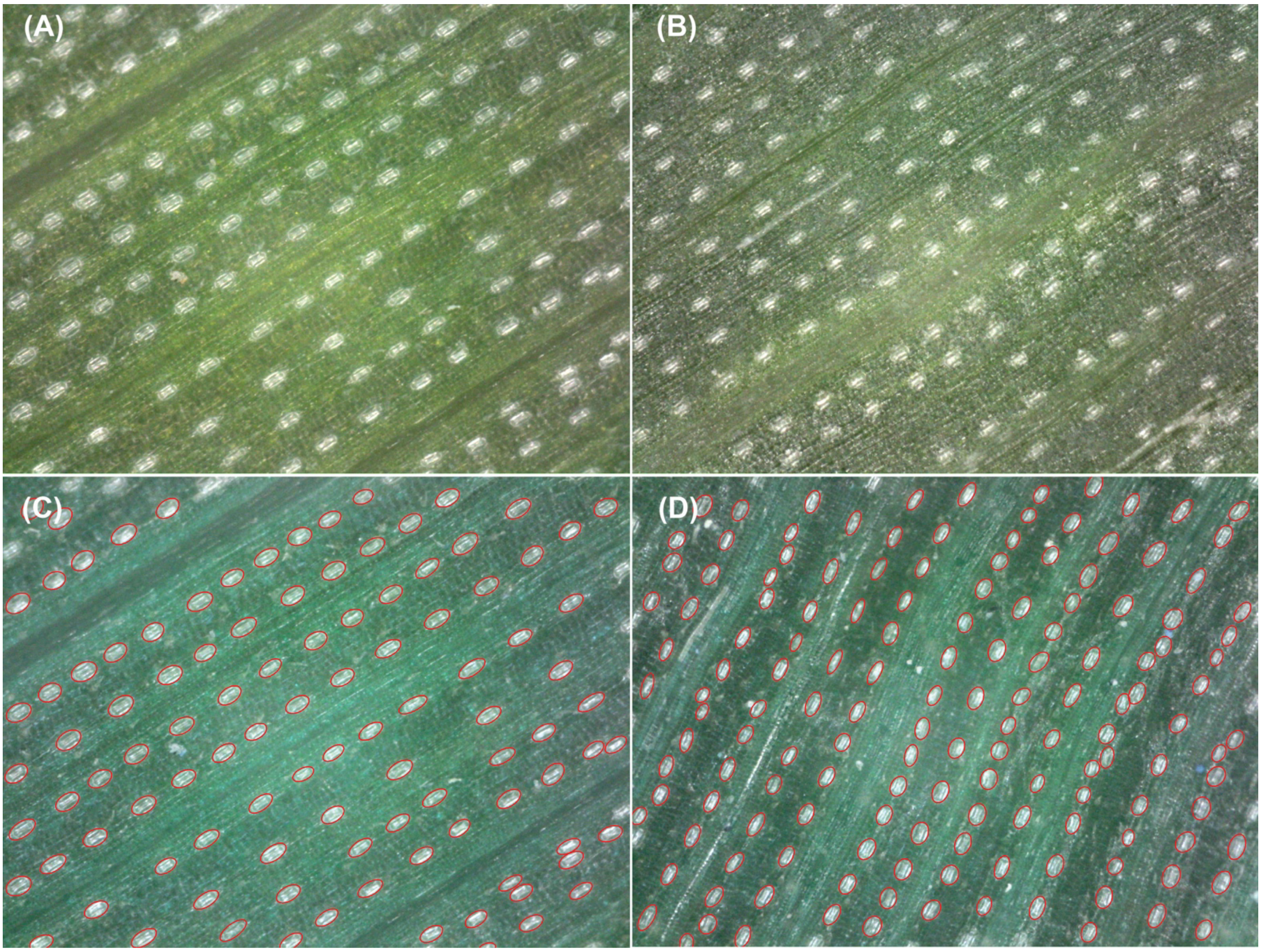


**Figure S1: Representative images of stomatal anatomy collected *in situ* using 200x magnification handheld digital microscope for rainfed vs irrigated trial.** (A) adaxial and (B) abaxial raw images collected using microscope. (C) and (D) show image with automatically labelled stomata with ellipses using the deep learning model we trained.

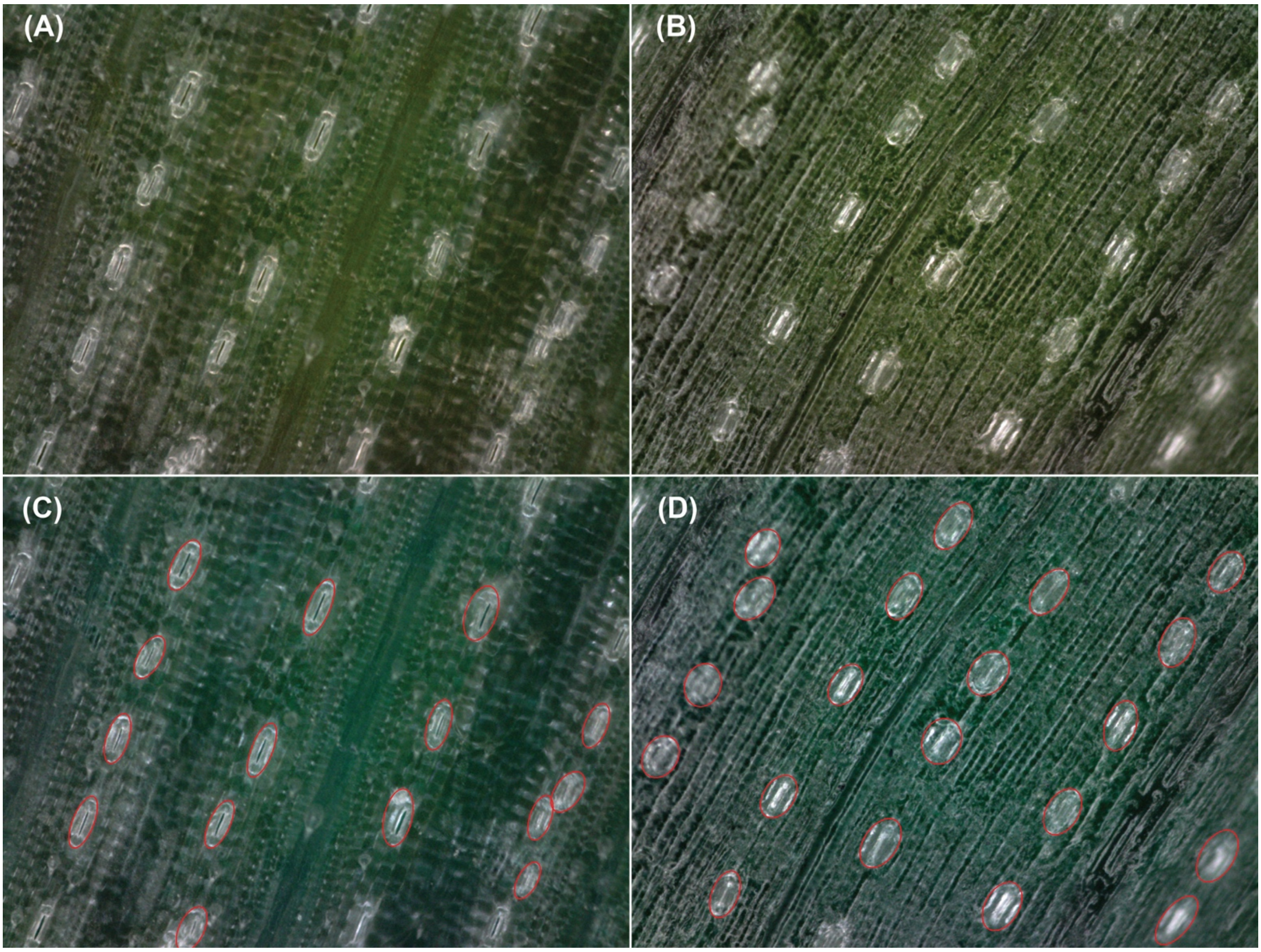


**Figure S2: Representative images of stomatal anatomy collected *in situ* using 400x magnification handheld digital microscope for TOS trial.** (A) adaxial and (B) abaxial raw images collected using microscope. (C) and (D) show image with automatically labelled stomata with ellipses using the deep learning model we trained.

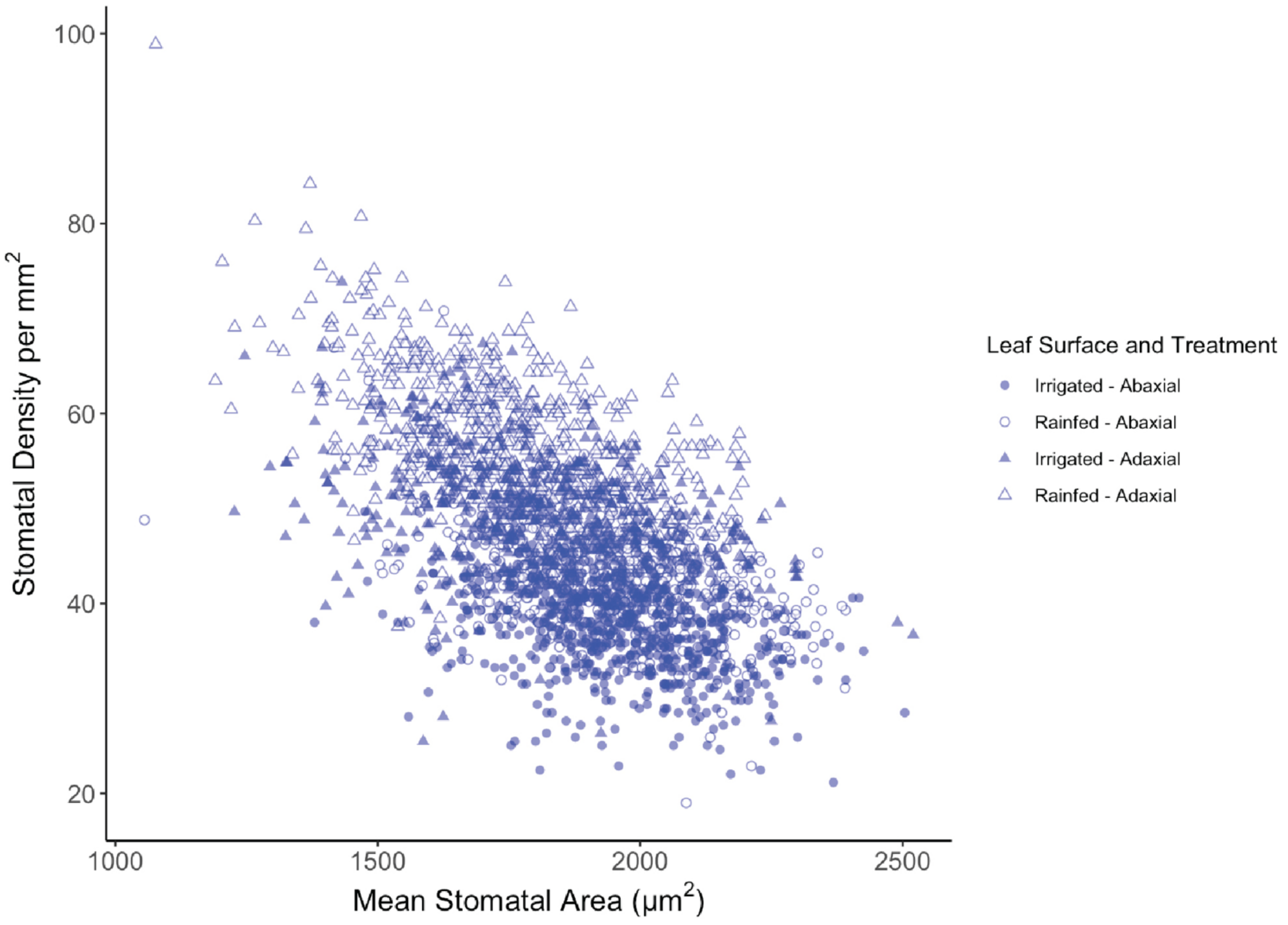


**Figure S3: Relationship between stomatal density and stomatal area for rainfed vs irrigated trial.** Shaded markers represent irrigated plants and unshaded markers represent rainfed plants. Circular markers represent abaxial leaf surface and triangular represent adaxial leaf surface.

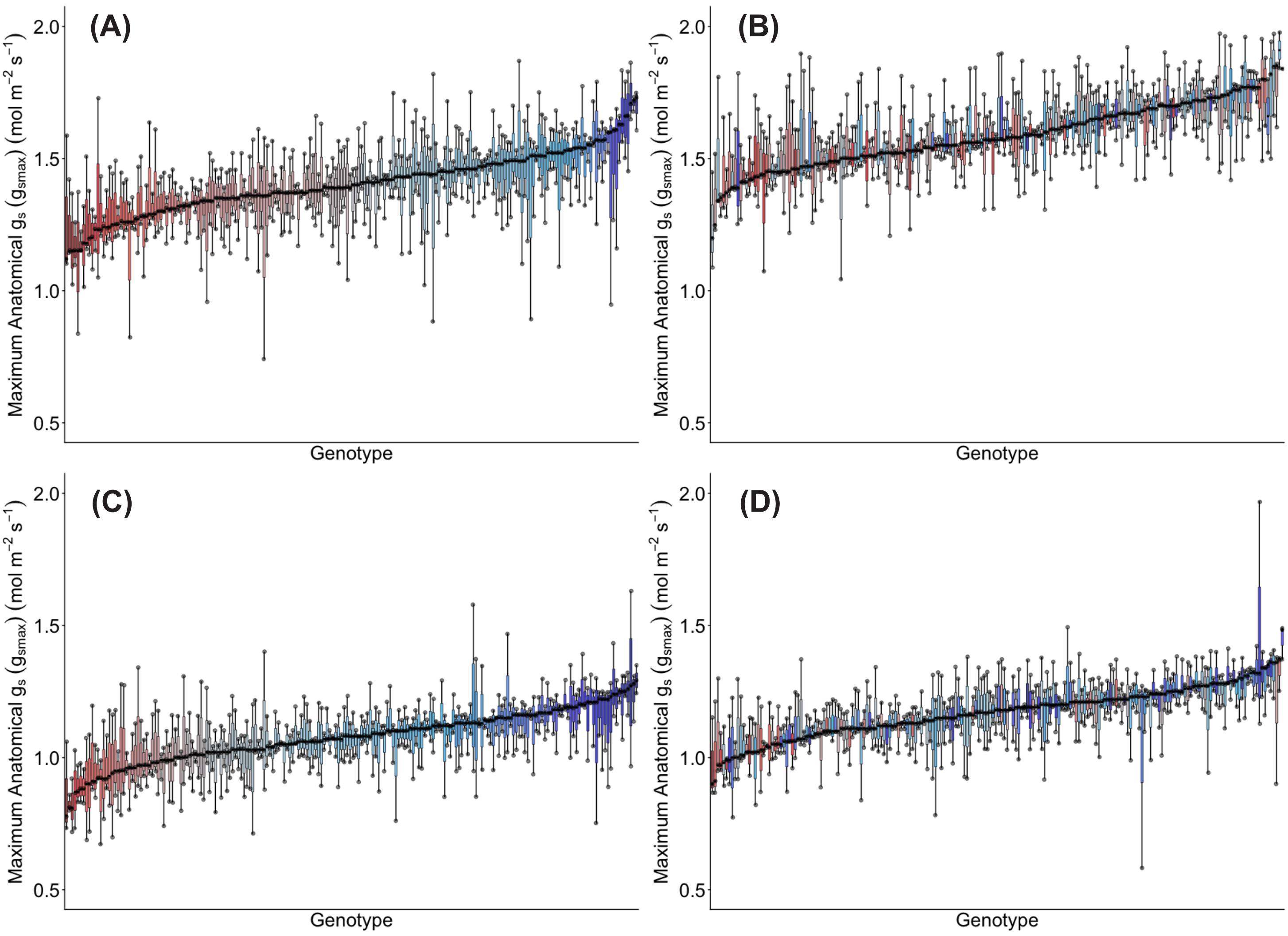


**Figure S4: Genotypic distribution of maximum anatomical stomatal conductance, $g_{smax}$, across 200 wheat genotypes under two watering treatments for rainfed vs irrigated trial.** (A) Adaxial surface Irrigated; (B) adaxial surface Rainfed; (C) abaxial surface Irrigated; and (D) abaxial surface Rainfed. Genotypes are ranked by median $g_{smax}$. Colour assigned to each genotype based on irrigated $g_{smax}$. Thick horizontal lines within boxes indicate the median and boxes indicate the upper (75%) and lower (25%) quartiles. Whiskers indicate the ranges of the minimum and maximum values. Points indicate individual measurements.

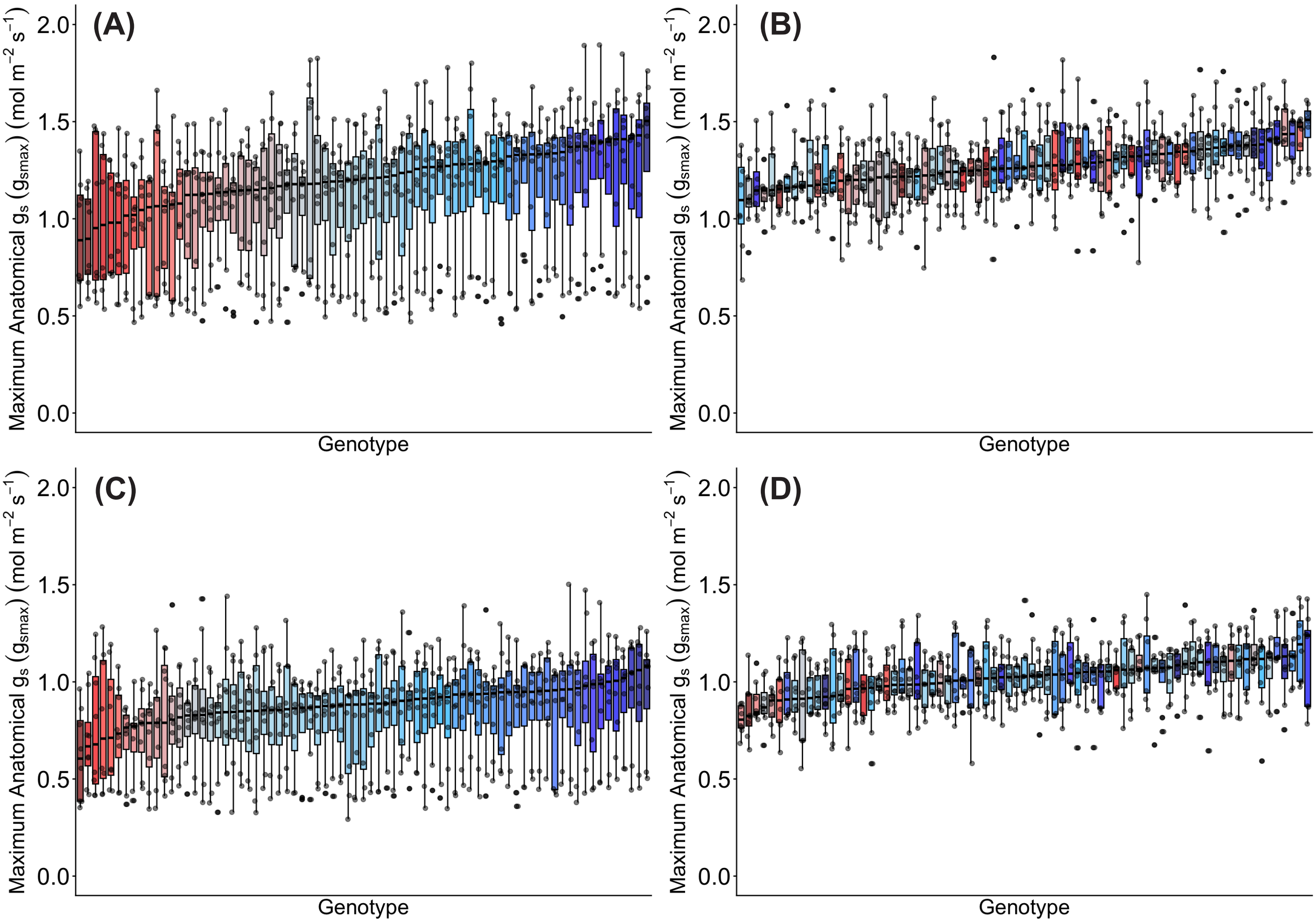


**Figure S5: Genotypic distribution of maximum anatomical stomatal conductance, *$g_{smax}$*, across 75 wheat genotypes at two sowing times for TOS trial.** (A) Adaxial surface TOS 1; (B) adaxial surface TOS 2; (C) abaxial surface TOS 1; and (D) abaxial surface TOS 2. Genotypes are ranked by median $g_{smax}$. Colour assigned to each genotype based on TOS 1 $g_{smax}$. Thick horizontal lines within boxes indicate the median and boxes indicate the upper (75%) and lower (25%) quartiles. Whiskers indicate the ranges of the minimum and maximum values. Points indicate individual measurements.

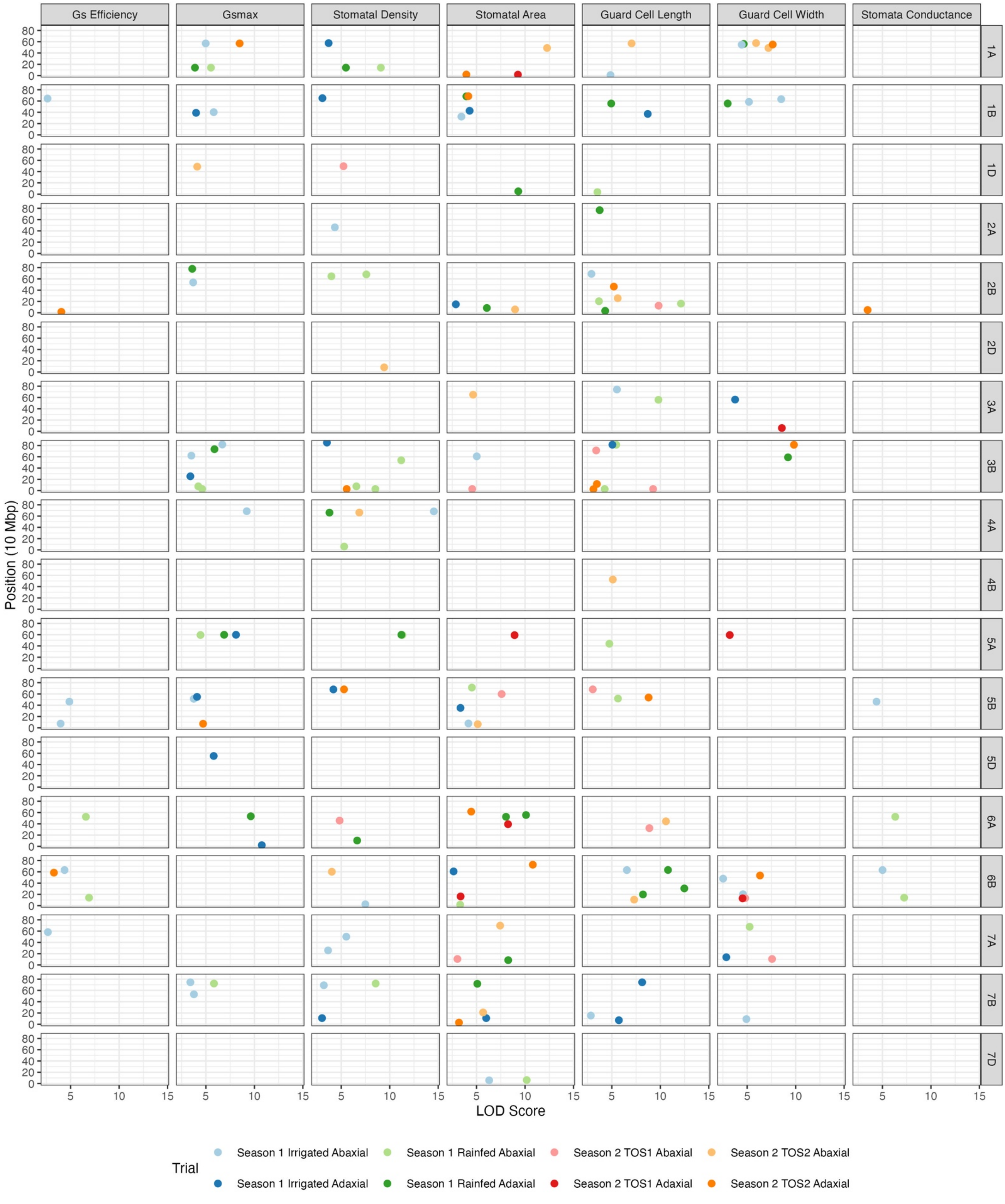


**Figure S6: LOD score of putative QTLs by trait and chromosome**. The scatterplot shows the LOD scores of putatiative QTLs identified by chromosome and position (in mega base pairs), with each color corresponding to a specific season, treatment and surface.

**Table S1: Overview of germplasm screened in rainfed vs irrigated HeDWIC field trials.**

| No. | Designation | No. | Designation |
|---|---|---|---|
| 1 | PBI09C009-BC-DH80 | 101 | PBI17N013-0C-0N-0N-010N-8N |
| 2 | PBI09C008-BC-DH17 | 102 | PBI17N013-0C-0N-0N-010N-9N |
| 3 | PBI09C043-BC-0C-20N-99N | 103 | PBI17N014-0C-0N-0N-010N-4N |
| 4 | PBI09C048-BC-0C-6N-99N | 104 | PBI17N014-0C-0N-0N-010N-5N |
| 5 | ACIAR09PBI C04-52C-DH3 | 105 | PBI17N014-0C-0N-0N-010N-6N |
| 6 | ACIAR09PBI C04-52C-DH10 | 106 | PBI17N014-0C-0N-0N-010N-8N |
| 7 | PBI07C101-DH96 | 107 | PBI17N014-0C-0N-0N-010N-16N |
| 8 | PBI07C101-DH147 | 108 | PBI17N014-0C-0N-0N-010N-19N |
| 9 | CMSA05Y00954T-040M-040ZTP0Y-040ZTM-040SY-12ZTM-01Y-0B | 109 | PBI17N015-0C-0N-0N-010N-2N |
| 10 | SUNTOP | 110 | PBI17N015-0C-0N-0N-010N-5N |
| 11 | ICW02.00099-11APTS-0AP-0AP-9AP-0AP | 111 | PBI17N015-0C-0N-0N-010N-8N |
| 12 | ICW04.20024-9AP-0AP-0AP-0AP-9AP-0AP | 112 | PBI17N015-0C-0N-0N-010N-10N |
| 13 | CMSS08B00645T-099TOPY-099M-099NJ-099NJ-6WGY-0B | 113 | PBI17N015-0C-0N-0N-010N-19N |
| 14 | CMSS08B00648T-099TOPY-099M-099NJ-2WGY-0B | 114 | PBI17N015-0C-0N-0N-010N-24N |
| 15 | CMSA08Y00613S-050Y-050ZTM-050Y-59BMX-010Y-0B | 115 | PBI17N016-0C-0N-0N-010N-27N |
| 16 | CMSA08M00007T-030(AWW5L7BO1HET)Y-040M-0NJ-4Y-0B | 116 | PBI17N018-0C-0N-0N-010N-1N |
| 17 | SUNLIN | 117 | PBI17N019-0C-0N-0N-010N-6N |
| 18 | CMSS06Y00885T-099TOPM-099Y-099ZTM-099NJ-099NJ-26WGY-0B | 118 | PBI17N020-0C-0N-0N-010N-12N |
| 19 | CMSA06M00008T-024(PINBD1BHET)Y-040ZTM-026(PINBD1BPOS)ZTY-20ZTM-0Y-0B | 119 | PBI17N021-0C-0N-0N-010N-5N |
| 20 | CMSA05M00140S-021(CRE1M19-GWM577BO1)M-028(BO1 POS& CRE1 POS)ZTY-040ZTM-040SY-11ZTM-0Y-0B | 120 | PBI16C001-0C-0N-0N-010N-5N |
| 21 | MACE | 121 | PBI16C007-0C-0N-0N-010N-2N |
| 22 | SCOUT | 122 | PBI16C008-0C-0N-0N-010N-7N |
| 23 | CMSA05Y01186T-040M-040ZTP0Y-040ZTM-040SY-32ZTM-02Y-0B | 123 | PBI16C009-0C-0N-0N-010N-14N |
| 24 | CGSS05B00258T-099TOPY-099M-099Y-099ZTM-12WGY-0B | 124 | PBI16C009-0C-0N-0N-010N-15N |
| 25 | SUNCO | 125 | PBI16C009-0C-0N-0N-010N-17N |

| | | | |
|---|---|---|---|
| 26 | BIOINTA 1005 | 126 | PBI16C010-0C-0N-0N-010N-6N |
| 27 | BORLAUG 100 | 127 | PBI16C011-0C-0N-0N-010N-1N |
| 28 | COOLAH | 128 | PBI16C011-0C-0N-0N-010N-5N |
| 29 | CUTLASS | 129 | PBI16C011-0C-0N-0N-010N-10N |
| 30 | SCEPTER | 130 | PBI16C013-0C-0N-0N-010N-4N |
| 31 | CONDO | 131 | PBI16C013-0C-0N-0N-010N-12N |
| 32 | FLANKER | 132 | PBI16C015-0C-0N-0N-010N-10N |
| 33 | VIKING | 133 | PBI16C015-0C-0N-0N-010N-13N |
| 34 | ICW05.0660-18AP-0AP-0AP-1AP-0AP | 134 | PBI16C016-0C-0N-0N-010N-11N |
| 35 | CMSS11B00167S-099M-0SY-11M-0WGY | 135 | PBI16C016-0C-0N-0N-010N-14N |
| 36 | PTSS02B00096T-0TOPY-0B-0Y-0B-33Y-0M-0SY-0Y-0Y | 136 | TUK004-0Q-0N-020N-0N-5N |
| 37 | PTSS02B00094T-0TOPY-0B-0Y-0B-3Y-0ZTB-0SY-0Y-0Y | 137 | TUK004-0Q-0N-020N-0N-8N |
| 38 | PTSS02Y00021S-099B-099Y-030ZTM-040SY-040M-21Y-0M-0SY-0Y-0Y | 138 | TUK004-0Q-0N-020N-0N-36N |
| 39 | PTSS02Y00023S-099B-099Y-099B-099Y-100B-0Y | 139 | PTSS15Y00010S-099B-099Y-099M-2Y-020Y |
| 40 | PTSS08GHB00013S-0SH-099SHB-099SHB-099SHB-32Y-0Y | 140 | PTSS15Y00023S-099B-099Y-099M-23Y-020Y |
| 41 | PTSS08GHB00016S-099SH-099Y-099SHB-099Y-155Y-0Y | 141 | PTSS15Y00024S-099B-099Y-099M-2Y-020Y |
| 42 | SDSS12B00849T-0Y-0B-0B-12Y-0B | 142 | PTSS15Y00024S-099B-099Y-099M-10Y-020Y |
| 43 | SUNCHASER | 143 | PTSS15Y00024S-099B-099Y-099M-17Y-020Y |
| 44 | VIXEN | 144 | PTSS14Y00062S-0B-099Y-099B-39Y-020Y |
| 45 | IMPALA | 145 | PTSS14Y00071S-0B-099Y-099B-33Y-020Y |
| 46 | RELIANT | 146 | PTSS14Y00328S-0B-099Y-099B-32Y-020Y |
| 47 | BECKOM | 147 | PTSS14Y00329S-0B-099Y-099B-33Y-020Y |
| 48 | HELLFIRE | 148 | PTSS11Y00152S-0SHB-099B-099Y-099B-099Y-17Y-020Y-0B |
| 49 | MUSTANG | 149 | PTSS15Y00053S-099B-099Y-099M-8Y-020Y-0B |
| 50 | PBIC15039-0C-9N-010N-2N-0N | 150 | PTSS15Y00056S-099B-099Y-099M-14Y-020Y-0B |
| 51 | PBIC15038-0C-29N-010N-1N-0N | 151 | PTSS15Y00082S-099B-099Y-099M-16Y-020Y-0B |

| | | | |
|---|---|---|---|
| 52 | PBIC15034-0C-24N-010N-1N-0N | 152 | PTSS02B00102T-0TOPY-0B-0Y-0B-11Y-0M-0SY-0B-0Y-4Y-0M |
| 53 | PBIC15034-0C-63N-010N-1N-0N | 153 | PTSS14Y00330S-0B-099Y-099B-40Y-0B |
| 54 | PBIC15034-0C-73N-010N-1N-0N | 154 | PTSS14Y00057S-0B-099Y-099B-10Y-020Y-0B |
| 55 | PBIC15030-0C-13N-010N-1N-0N | 155 | PTSS14Y00013S-0B-099Y-099B-14Y-020Y-0B |
| 56 | PBI19N002-0N-5N | 156 | PTSS15Y00032S-099B-099Y-099M-25Y-020Y |
| 57 | PBI19N003-0N-11N | 157 | PTSS15Y00050S-099B-099Y-099M-1Y-020Y |
| 58 | PBI19N007-0N-3N | 159 | SUNMASTER |
| 59 | PBI19N007-0N-6N | 158 | CATAPULT |
| 60 | PBI19N007-0N-8N | 160 | ROCKSTAR |
| 61 | PBI17N001-0N-0N-12N | 161 | SHERIFF CL PLUS |
| 62 | PBI17N002-0N-0N-38N | 162 | VALIANT CL PLUS |
| 63 | PBI17N002-0N-0N-76N | 163 | HAVOC |
| 64 | PBI17N003-0N-0N-32N | 164 | STEALTH |
| 65 | PBI17N004-0N-0N-16N | 165 | IG-ANU-HeatLine-001 |
| 66 | PBI17N004-0N-0N-18N | 166 | IG-ANU-HeatLine-002 |
| 67 | PBI17N005-0N-0N-29N | 167 | IG-ANU-HeatLine-008 |
| 68 | PBI17N005-0N-0N-32N | 168 | IG-ANU-HeatLine-009 |
| 69 | PBI17N006-0N-0N-89N | 169 | IG-ANU-HeatLine-011 |
| 70 | PBI17N006-0N-0N-96N | 170 | IG-ANU-HeatLine-012 |
| 71 | PBI17N007-0N-0N-13N | 171 | IG-ANU-HeatLine-014 |
| 72 | PBI17N008-0N-0N-10N | 172 | IG-ANU-HeatLine-015 |
| 73 | PBI17N008-0N-0N-113N | 173 | IG-ANU-HeatLine-019 |
| 74 | PBI17N009-0N-0N-4N | 174 | IG-ANU-HeatLine-023 |
| 75 | PBI17N009-0N-0N-46N | 175 | IG-ANU-HeatLine-027 |
| 76 | PBI17N011-0N-0N-20N | 176 | IG-ANU-HeatLine-028 |
| 77 | PBI17N011-0N-0N-41N | 177 | IG-ANU-HeatLine-033 |
| 78 | PBI17N012-0N-0N-106N | 178 | IG-ANU-HeatLine-034 |
| 79 | PBI17N013-0N-0N-3N | 179 | IG-ANU-HeatLine-036 |
| 80 | PBI17N013-0N-0N-62N | 180 | IG-ANU-HeatLine-038 |
| 81 | PBI17N014-0N-0N-54N | 181 | IG-ANU-HeatLine-040 |
| 82 | PBI17N015-0N-0N-58N | 182 | IG-ANU-HeatLine-041 |
| 83 | PBI17N015-0N-0N-96N | 183 | IG-ANU-HeatLine-052 |
| 84 | PBI17N001-0C-0N-0N-010N-8N | 184 | IG-ANU-HeatLine-055 |
| 85 | PBI17N002-0C-0N-0N-010N-16N | 185 | IG-ANU-HeatLine-056 |

| | | | |
|---|---|---|---|
| 86 | PBI17N004-0C-0N-0N-010N-5N | 186 | IG-ANU-HeatLine-060 |
| 87 | PBI17N005-0C-0N-0N-010N-8N | 187 | IG-ANU-HeatLine-062 |
| 88 | PBI17N005-0C-0N-0N-010N-10N | 188 | IG-ANU-HeatLine-065 |
| 89 | PBI17N005-0C-0N-0N-010N-14N | 189 | IG-ANU-HeatLine-066 |
| 90 | PBI17N006-0C-0N-0N-010N-3N | 190 | IG-ANU-HeatLine-070 |
| 91 | PBI17N007-0C-0N-0N-010N-8N | 191 | IG-ANU-HeatLine-072 |
| 92 | PBI17N007-0C-0N-0N-010N-11N | 192 | IG-ANU-HeatLine-073 |
| 93 | PBI17N007-0C-0N-0N-010N-15N | 193 | IG-ANU-HeatLine-074 |
| 94 | PBI17N009-0C-0N-0N-010N-10N | 194 | IG-ANU-HeatLine-078 |
| 95 | PBI17N010-0C-0N-0N-010N-5N | 195 | IG-ANU-HeatLine-087 |
| 96 | PBI17N010-0C-0N-0N-010N-6N | 196 | IG-ANU-HeatLine-093 |
| 97 | PBI17N010-0C-0N-0N-010N-10N | 197 | IG-ANU-HeatLine-096 |
| 98 | PBI17N010-0C-0N-0N-010N-13N | 198 | IG-ANU-HeatLine-098 |
| 99 | PBI17N012-0C-0N-0N-010N-17N | 199 | IG-ANU-HeatLine-105 |
| 100 | PBI17N013-0C-0N-0N-010N-7N | 200 | IG-ANU-HeatLine-110 |

**Table S2: Overview of germplasm screened in TOS HeDWIC field trials. All lines grown in TOS trials were grown in rainfed vs irrigated trials.**

| No. | Designation | No. | Designation |
|---|---|---|---|
| 1 | 107:ZIZ13 | 39 | PBI16C009-0C-0N-0N-010N-14N |
| 2 | 108:ZWB10 | 40 | PBI16C011-0C-0N-0N-010N-5N |
| 3 | 13:ZSA20 | 41 | PBI16C015-0C-0N-0N-010N-10N |
| 4 | 137:ZWB17 | 42 | PBI17N002-0C-0N-0N-010N-16N |
| 5 | 14:ZSA20 | 43 | PBI17N003-0N-0N-32N |
| 6 | 16:ZWY20 | 44 | PBI17N004-0C-0N-0N-010N-5N |
| 7 | 171:ZWB13 | 45 | PBI17N004-0N-0N-16N |
| 8 | 18:ZSA20 | 46 | PBI17N004-0N-0N-18N |
| 9 | 24:ZSA20 | 47 | PBI17N005-0C-0N-0N-010N-10N |
| 10 | 25:ZWY20 | 48 | PBI17N005-0C-0N-0N-010N-8N |
| 11 | 26:ZWY20 | 49 | PBI17N006-0N-0N-89N |
| 12 | 266:ZWB13 | 50 | PBI17N007-0C-0N-0N-010N-15N |
| 13 | 7:ZSA20 | 51 | PBI17N007-0C-0N-0N-010N-8N |
| 14 | CATAPULT | 52 | PBI17N009-0C-0N-0N-010N-10N |
| 15 | COOLAH | 53 | PBI17N009-0N-0N-4N |
| 16 | HAVOC | 54 | PBI17N011-0N-0N-41N |
| 17 | HELLFIRE | 55 | PBI17N013-0C-0N-0N-010N-8N |
| 18 | IG-ANU-HeatLine-002 | 56 | PBI17N013-0C-0N-0N-010N-9N |
| 19 | IG-ANU-HeatLine-008 | 57 | PBI17N014-0C-0N-0N-010N-16N |
| 20 | IG-ANU-HeatLine-033 | 58 | PBI17N014-0C-0N-0N-010N-4N |
| 21 | IG-ANU-HeatLine-034 | 59 | PBI17N014-0C-0N-0N-010N-6N |
| 22 | IG-ANU-HeatLine-040 | 60 | PBI17N014-0N-0N-54N |
| 23 | IG-ANU-HeatLine-052 | 61 | PBI17N015-0C-0N-0N-010N-5N |
| 24 | IG-ANU-HeatLine-055 | 62 | PBI17N016-0C-0N-0N-010N-27N |
| 25 | IG-ANU-HeatLine-066 | 63 | PBI17N021-0C-0N-0N-010N-5N |
| 26 | IG-ANU-HeatLine-078 | 64 | PBI19N007-0N-3N |
| 27 | IG-ANU-HeatLine-093 | 65 | PBIC15034-0C-24N-010N-1N-0N |
| 28 | IMPALA | 66 | ROCKSTAR |
| 29 | ISR1086.123 | 67 | SHERIFF_CL_PLUS |
| 30 | ISR1086.5 | 68 | STEALTH |
| 31 | ISR1086.60 | 69 | SUNCHASER |
| 32 | ISR1086.7 | 70 | SUNCO |
| 33 | MACE | 71 | SUNMASTER |
| 34 | MUSTANG | 72 | TUK004-0Q-0N-020N-0N-36N |
| 35 | PBI07C101-DH147 | 73 | VALIENT |
| 36 | PBI09C043-BC-0C-20N-99N | 74 | VIKING |
| 37 | PBI09C048-BC-0C-6N-99N | 75 | VIXEN |
| 38 | PBI16C008-0C-0N-0N-010N-7N | | |

**Table S3: Average soil moisture content at planting for rainfed vs irrigated trial and TOS trial.**

| | Soil Moisture (%) | | | |
|---|---|---|---|---|
| **Depth (cm)** | **S1 - Rainfed** | **S1 - Irrigated** | **S2 - TOS 1** | **S2 - TOS 2** |
| **10** | 23.4 | 22.2 | 24.5 | 33.6 |
| **20** | 25.8 | 26.9 | 24.7 | 36.9 |
| **40** | 26.9 | 28.9 | 25.7 | 35.1 |
| **60** | 28.2 | 29.1 | 23.5 | 32.4 |
| **80** | 28.4 | 29.1 | 23.7 | 32.7 |
| **100** | 26.9 | 26.6 | 22.1 | 31.4 |
| **120** | 23.5 | 22.8 | 23.0 | 29.1 |

**Table S4: Performance metrics for YOLOv8-M model.** Metrics used were: Precision, Recall, mAP50 (mean average precision at Intersection over Union = 0.5) and mAP50-95 (mean average precision at Intersection over Union from 0.5 to 0.95).

| Metric | Value |
|---|---|
| Precision (M) | 0.9568 |
| Recall (M) | 0.9602 |
| mAP59 (M) | 0.9441 |
| mAP50-95 (M) | 0.6775 |

**Table S5: Rainfed vs irrigated trial data showing average values for key traits under Irrigated and Rainfed treatments, and % change from Irrigated to Rainfed.** Values are given for each surface independently and for both surfaces averaged.

| Trait & Surface | | Irrigated | Rainfed | % Change Irrigated to Rainfed |
|---|---|---|---|---|
| **Abaxial** | | | | |
| $g_s$ | mol.m$^{-2}$.s$^{-1}$ | 0.082 | 0.062 | -24.02 |
| **Stomatal Density** | per mm$^{-2}$ | 37.29 | 43.21 | 15.89 |
| **Guard cell Length** | µm | 68.39 | 64.30 | -5.98 |
| **Guard Cell Width** | µm | 40.96 | 42.49 | 3.74 |
| **Stomatal Area** | µm$^{-2}$ | 1960.00 | 1936.64 | -1.19 |
| $g_{smax}$ | mol.m$^{-2}$.s$^{-1}$ | 1.07 | 1.17 | 8.90 |
| $g_{se}$ | - | 0.08 | 0.05 | -30.09 |
| **Adaxial** | | | | |
| $g_s$ | mol.m$^{-2}$.s$^{-1}$ | 0.195 | 0.140 | -28.08 |
| **Stomatal Density** | per mm$^{-2}$ | 48.46 | 57.41 | 18.47 |
| **Guard cell Length** | µm | 68.55 | 66.04 | -3.66 |
| **Guard Cell Width** | µm | 38.99 | 38.35 | -1.66 |
| **Stomatal Area** | µm$^{-2}$ | 1825.32 | 1752.90 | -3.97 |
| $g_{smax}$ | mol.m$^{-2}$.s$^{-1}$ | 1.40 | 1.60 | 14.00 |
| $g_{se}$ | - | 0.14 | 0.09 | -37.17 |
| **Adaxial & Abaxial Average** | | | | |
| $g_s$ | mol.m$^{-2}$.s$^{-1}$ | 0.138 | 0.101 | -26.88 |
| **Stomatal Density** | per mm$^{-2}$ | 42.87 | 50.31 | 17.35 |
| **Guard cell Length** | µm | 68.47 | 65.17 | -4.82 |
| **Guard Cell Width** | µm | 39.98 | 40.42 | 1.10 |
| **Stomatal Area** | µm$^{-2}$ | 1892.66 | 1844.77 | -2.53 |
| $g_{smax}$ | mol.m$^{-2}$.s$^{-1}$ | 1.24 | 1.38 | 11.79 |
| $g_{se}$ | - | 0.11 | 0.07 | -34.66 |
| **Yield** | t.ha$^{-1}$ | 5.65 | 4.43 | -21.59 |
| **Thousand Kernel Weight** | g | 36.15 | 33.87 | -6.32 |
| **Screenings** | % | 4.19 | 4.52 | 7.87 |
| **Moisture** | % | 10.93 | 11.14 | 1.89 |
| **Protein** | % | 12.99 | 12.54 | -3.46 |
| **Test Weight** | kg.hL$^{-1}$ | 79.90 | 81.78 | 2.35 |

**Table S6: TOS trial data showing average values for key traits at TOS 1 and at TOS 2, and % change from TOS 1 to TOS 2.** Values are given for each surface independently and for both surfaces averaged.

| Trait & Surface | | TOS 1 | TOS 2 | % Change TOS 1 to TOS 2 |
|---|---|---|---|---|
| **Abaxial** | | | | |
| $g_s$ | mol.m$^{-2}$.s$^{-1}$ | 0.143 | 0.121 | -15.70 |
| **Stomatal Density** | per mm$^{-2}$ | 34.57 | 38.78 | 12.21 |
| **Guard cell Length** | µm | 57.02 | 62.43 | 9.48 |
| **Guard Cell Width** | µm | 34.03 | 38.72 | 13.79 |
| **Stomatal Area** | µm$^{-2}$ | 1599.27 | 1884.75 | 17.85 |
| $g_{smax}$ | mol.m$^{-2}$.s$^{-1}$ | 0.83 | 1.02 | 22.25 |
| $g_{se}$ | - | 0.20 | 0.12 | -38.93 |
| **Adaxial** | | | | |
| $g_s$ | mol.m$^{-2}$.s$^{-1}$ | 0.311 | 0.270 | -12.97 |
| **Stomatal Density** | per mm$^{-2}$ | 45.25 | 48.08 | 6.24 |
| **Guard cell Length** | µm | 58.35 | 62.95 | 7.88 |
| **Guard Cell Width** | µm | 31.32 | 35.95 | 14.78 |
| **Stomatal Area** | µm$^{-2}$ | 1494.73 | 1752.75 | 17.26 |
| $g_{smax}$ | mol.m$^{-2}$.s$^{-1}$ | 1.12 | 1.27 | 13.95 |
| $g_{se}$ | - | 0.31 | 0.22 | -30.05 |
| **Adaxial & Abaxial Average** | | | | |
| $g_s$ | mol.m$^{-2}$.s$^{-1}$ | 0.227 | 0.196 | -13.83 |
| **Stomatal Density** | per mm$^{-2}$ | 39.91 | 43.43 | 8.84 |
| **Guard cell Length** | µm | 57.69 | 62.69 | 8.67 |
| **Guard Cell Width** | µm | 32.68 | 37.34 | 14.26 |
| **Stomatal Area** | µm$^{-2}$ | 1547.00 | 1818.70 | 17.56 |
| $g_{smax}$ | mol.m$^{-2}$.s$^{-1}$ | 0.98 | 1.15 | 17.51 |
| $g_{se}$ | - | 0.26 | 0.17 | -33.53 |
| **Yield** | t.ha$^{-1}$ | 5.77 | 2.82 | -51.13 |
| **Thousand Kernel Weight** | g | 41.86 | 32.17 | -23.15 |
| **Screenings** | % | 1.83 | 6.19 | 237.52 |
| **Moisture** | % | 10.23 | 10.19 | -0.41 |
| **Protein** | % | 12.08 | 13.74 | 13.74 |
| **Test Weight** | kg.hL$^{-1}$ | 86.10 | 79.12 | -8.11 |

**Table S7: List of the putative QTL candidates for each trait, organized by trial (indexed by season and treatment) and surface.** The chromosome, position (in base pairs), effect size and LOD score for each corresponding QTL are also provided.

| Trait | Surface | Treatment | Season | Chromosome | Position (bp) | Size | LOD |
|---|---|---|---|---|---|---|---|
| Guard Cell Length | Abaxial | Rainfed | Season 1 | 1D | 36,448,754 | 1.08 | 3.51 |
| Guard Cell Length | Abaxial | Rainfed | Season 1 | 2B | 161,587,608 | 1.26 | 12.13 |
| Guard Cell Length | Abaxial | Rainfed | Season 1 | 2B | 204,288,596 | -0.71 | 3.67 |
| Guard Cell Length | Abaxial | Rainfed | Season 1 | 3A | 558,339,875 | -1.10 | 9.81 |
| Guard Cell Length | Abaxial | Rainfed | Season 1 | 3B | 28,249,993 | 0.91 | 4.26 |
| Guard Cell Length | Abaxial | Rainfed | Season 1 | 3B | 813,286,435 | 0.63 | 5.45 |
| Guard Cell Length | Abaxial | Rainfed | Season 1 | 5A | 439,046,386 | -0.69 | 4.74 |
| Guard Cell Length | Abaxial | Rainfed | Season 1 | 5B | 518,333,082 | -0.83 | 5.63 |
| Guard Cell Length | Adaxial | Rainfed | Season 1 | 1B | 555,625,935 | 0.99 | 4.94 |
| Guard Cell Length | Adaxial | Rainfed | Season 1 | 2A | 767,262,218 | 1.01 | 3.74 |
| Guard Cell Length | Adaxial | Rainfed | Season 1 | 2B | 33,525,704 | -0.75 | 4.31 |
| Guard Cell Length | Adaxial | Rainfed | Season 1 | 6B | 199,072,853 | 1.26 | 8.21 |
| Guard Cell Length | Adaxial | Rainfed | Season 1 | 6B | 305,715,199 | -2.22 | 12.48 |
| Guard Cell Length | Adaxial | Rainfed | Season 1 | 6B | 632,822,413 | 1.66 | 10.79 |
| Guard Cell Length | Abaxial | Irrigated | Season 1 | 1A | 8,527,135 | -1.61 | 4.86 |
| Guard Cell Length | Abaxial | Irrigated | Season 1 | 2B | 688,832,053 | 1.03 | 2.89 |

| | | | | | | | |
|---|---|---|---|---|---|---|---|
| Guard Cell Length | Abaxial | Irrigated | Season 1 | 3A | 741,357,936 | -1.30 | 5.52 |
| Guard Cell Length | Abaxial | Irrigated | Season 1 | 6B | 631,016,005 | -1.33 | 6.55 |
| Guard Cell Length | Abaxial | Irrigated | Season 1 | 7B | 154,317,772 | -0.97 | 2.83 |
| Guard Cell Length | Adaxial | Irrigated | Season 1 | 1B | 371,130,997 | -1.50 | 8.70 |
| Guard Cell Length | Adaxial | Irrigated | Season 1 | 3B | 813,648,022 | 0.88 | 5.05 |
| Guard Cell Length | Adaxial | Irrigated | Season 1 | 7B | 71,275,260 | -1.53 | 5.72 |
| Guard Cell Length | Adaxial | Irrigated | Season 1 | 7B | 744,071,081 | -1.56 | 8.12 |
| Guard Cell Length | Abaxial | TOS1 | Season 2 | 2B | 125,065,337 | 1.61 | 9.82 |
| Guard Cell Length | Abaxial | TOS1 | Season 2 | 3B | 28,249,993 | 1.81 | 9.26 |
| Guard Cell Length | Abaxial | TOS1 | Season 2 | 3B | 711,988,819 | -2.19 | 3.38 |
| Guard Cell Length | Abaxial | TOS1 | Season 2 | 5B | 681,423,027 | -0.63 | 3.03 |
| Guard Cell Length | Abaxial | TOS1 | Season 2 | 6A | 323,594,208 | -1.01 | 8.87 |
| Guard Cell Length | Abaxial | TOS2 | Season 2 | 1A | 569,847,820 | 1.02 | 7.04 |
| Guard Cell Length | Abaxial | TOS2 | Season 2 | 2B | 257,283,535 | -0.96 | 5.61 |
| Guard Cell Length | Abaxial | TOS2 | Season 2 | 4B | 526,124,380 | 1.78 | 5.10 |
| Guard Cell Length | Abaxial | TOS2 | Season 2 | 6A | 444,511,746 | -1.21 | 10.58 |
| Guard Cell Length | Abaxial | TOS2 | Season 2 | 6B | 106,688,178 | 1.97 | 7.30 |
| Guard Cell Length | Adaxial | TOS2 | Season 2 | 2B | 462,771,824 | -1.57 | 5.20 |
| Guard Cell Length | Adaxial | TOS2 | Season 2 | 3B | 24,990,535 | -1.07 | 3.10 |

| | | | | | | | |
|---|---|---|---|---|---|---|---|
| Guard Cell Length | Adaxial | TOS2 | Season 2 | 3B | 117,316,836 | 1.43 | 3.45 |
| Guard Cell Length | Adaxial | TOS2 | Season 2 | 5B | 536,696,466 | 2.61 | 8.79 |
| Guard Cell Width | Abaxial | Rainfed | Season 1 | 7A | 677,127,635 | -0.80 | 5.25 |
| Guard Cell Width | Adaxial | Rainfed | Season 1 | 1A | 560,537,130 | -0.69 | 4.64 |
| Guard Cell Width | Adaxial | Rainfed | Season 1 | 1B | 555,625,561 | 0.67 | 2.99 |
| Guard Cell Width | Adaxial | Rainfed | Season 1 | 3B | 589,989,645 | 0.97 | 9.21 |
| Guard Cell Width | Abaxial | Irrigated | Season 1 | 1A | 547,618,085 | -0.76 | 4.44 |
| Guard Cell Width | Abaxial | Irrigated | Season 1 | 1B | 585,013,536 | -0.75 | 5.19 |
| Guard Cell Width | Abaxial | Irrigated | Season 1 | 1B | 632,575,711 | -0.84 | 8.51 |
| Guard Cell Width | Abaxial | Irrigated | Season 1 | 6B | 203,090,199 | -0.68 | 4.57 |
| Guard Cell Width | Abaxial | Irrigated | Season 1 | 6B | 478,949,889 | 0.51 | 2.52 |
| Guard Cell Width | Abaxial | Irrigated | Season 1 | 7B | 91,203,950 | -1.30 | 4.92 |
| Guard Cell Width | Adaxial | Irrigated | Season 1 | 3A | 562,430,885 | 0.95 | 3.76 |
| Guard Cell Width | Adaxial | Irrigated | Season 1 | 7A | 137,223,473 | -0.92 | 2.85 |
| Guard Cell Width | Abaxial | TOS1 | Season 2 | 6B | 133,929,458 | -0.83 | 4.80 |
| Guard Cell Width | Abaxial | TOS1 | Season 2 | 7A | 105,813,606 | -0.81 | 7.57 |
| Guard Cell Width | Adaxial | TOS1 | Season 2 | 3A | 58,294,106 | 0.95 | 8.58 |
| Guard Cell Width | Adaxial | TOS1 | Season 2 | 5A | 593,322,475 | 0.56 | 3.21 |
| Guard Cell Width | Adaxial | TOS1 | Season 2 | 6B | 129,701,061 | 0.65 | 4.52 |

| | | | | | | | |
|---|---|---|---|---|---|---|---|
| Guard Cell Width | Abaxial | TOS2 | Season 2 | 1A | 489,023,694 | 2.26 | 7.20 |
| Guard Cell Width | Abaxial | TOS2 | Season 2 | 1A | 575,737,813 | 0.84 | 5.92 |
| Guard Cell Width | Adaxial | TOS2 | Season 2 | 1A | 550,414,031 | 0.94 | 7.64 |
| Guard Cell Width | Adaxial | TOS2 | Season 2 | 3B | 812,571,724 | -0.86 | 9.83 |
| Guard Cell Width | Adaxial | TOS2 | Season 2 | 6B | 535,019,306 | 0.76 | 6.33 |
| Stomatal Area | Abaxial | Rainfed | Season 1 | 5B | 713,283,368 | 40.28 | 4.52 |
| Stomatal Area | Abaxial | Rainfed | Season 1 | 6B | 16,991,758 | -50.82 | 3.31 |
| Stomatal Area | Abaxial | Rainfed | Season 1 | 7D | 61,474,690 | -79.58 | 10.18 |
| Stomatal Area | Adaxial | Rainfed | Season 1 | 1B | 682,638,576 | -38.42 | 3.95 |
| Stomatal Area | Adaxial | Rainfed | Season 1 | 1D | 49,690,814 | -129.03 | 9.31 |
| Stomatal Area | Adaxial | Rainfed | Season 1 | 2B | 84,346,596 | -51.84 | 6.06 |
| Stomatal Area | Adaxial | Rainfed | Season 1 | 6A | 524,424,694 | -79.71 | 8.04 |
| Stomatal Area | Adaxial | Rainfed | Season 1 | 6A | 555,940,653 | -66.30 | 10.10 |
| Stomatal Area | Adaxial | Rainfed | Season 1 | 7A | 85,140,641 | -48.45 | 8.27 |
| Stomatal Area | Adaxial | Rainfed | Season 1 | 7B | 717,428,482 | 39.87 | 5.08 |

| | | | | | | | |
|---|---|---|---|---|---|---|---|
| Stomatal Area | Abaxial | Irrigated | Season 1 | 1B | 323,674,856 | -42.97 | 3.44 |
| Stomatal Area | Abaxial | Irrigated | Season 1 | 3B | 607,408,705 | -45.50 | 5.02 |
| Stomatal Area | Abaxial | Irrigated | Season 1 | 5B | 76,477,566 | -36.35 | 4.16 |
| Stomatal Area | Abaxial | Irrigated | Season 1 | 7D | 56,638,565 | 56.37 | 6.30 |
| Stomatal Area | Adaxial | Irrigated | Season 1 | 1B | 426,913,815 | -80.01 | 4.28 |
| Stomatal Area | Adaxial | Irrigated | Season 1 | 2B | 149,197,334 | -100.24 | 2.86 |
| Stomatal Area | Adaxial | Irrigated | Season 1 | 5B | 352,556,732 | 62.51 | 3.34 |
| Stomatal Area | Adaxial | Irrigated | Season 1 | 6B | 607,438,025 | -42.97 | 2.63 |
| Stomatal Area | Adaxial | Irrigated | Season 1 | 7B | 108,638,412 | -58.29 | 6.00 |
| Stomatal Area | Abaxial | TOS1 | Season 2 | 3B | 28,249,993 | 65.81 | 4.55 |
| Stomatal Area | Abaxial | TOS1 | Season 2 | 5B | 597,807,739 | 83.26 | 7.59 |
| Stomatal Area | Abaxial | TOS1 | Season 2 | 7A | 105,815,982 | -32.93 | 3.03 |
| Stomatal Area | Adaxial | TOS1 | Season 2 | 1A | 15,675,703 | -73.99 | 9.27 |
| Stomatal Area | Adaxial | TOS1 | Season 2 | 5A | 591,176,975 | -59.37 | 8.92 |
| Stomatal Area | Adaxial | TOS1 | Season 2 | 6A | 393,060,345 | 58.91 | 8.25 |

| | | | | | | | |
|---|---|---|---|---|---|---|---|
| Stomatal Area | Adaxial | TOS1 | Season 2 | 6B | 163,654,888 | 54.24 | 3.36 |
| Stomatal Area | Abaxial | TOS2 | Season 2 | 1A | 489,023,694 | 173.78 | 12.27 |
| Stomatal Area | Abaxial | TOS2 | Season 2 | 2B | 57,852,165 | 62.88 | 8.97 |
| Stomatal Area | Abaxial | TOS2 | Season 2 | 3A | 649,063,085 | -41.52 | 4.65 |
| Stomatal Area | Abaxial | TOS2 | Season 2 | 5B | 63,154,049 | -45.71 | 5.12 |
| Stomatal Area | Abaxial | TOS2 | Season 2 | 7A | 697,601,843 | 43.90 | 7.43 |
| Stomatal Area | Abaxial | TOS2 | Season 2 | 7B | 210,120,507 | -43.77 | 5.68 |
| Stomatal Area | Adaxial | TOS2 | Season 2 | 1A | 17,925,924 | -51.13 | 3.95 |
| Stomatal Area | Adaxial | TOS2 | Season 2 | 1B | 685,348,909 | -55.01 | 4.15 |
| Stomatal Area | Adaxial | TOS2 | Season 2 | 6A | 616,437,868 | -104.86 | 4.45 |
| Stomatal Area | Adaxial | TOS2 | Season 2 | 6B | 726,374,830 | 91.66 | 10.80 |
| Stomatal Area | Adaxial | TOS2 | Season 2 | 7B | 29,648,398 | -52.44 | 3.19 |
| Stomatal Density | Abaxial | Rainfed | Season 1 | 1A | 139,877,951 | -0.13 | 9.09 |
| Stomatal Density | Abaxial | Rainfed | Season 1 | 2B | 646,101,250 | -0.09 | 3.98 |
| Stomatal Density | Abaxial | Rainfed | Season 1 | 2B | 679,575,828 | 0.13 | 7.59 |
| Stomatal Density | Abaxial | Rainfed | Season 1 | 3B | 28,249,993 | -0.17 | 8.52 |

| | | | | | | | |
|---|---|---|---|---|---|---|---|
| Stomatal Density | Abaxial | Rainfed | Season 1 | 3B | 75,468,220 | -0.26 | 6.56 |
| Stomatal Density | Abaxial | Rainfed | Season 1 | 3B | 537,494,407 | -0.13 | 11.19 |
| Stomatal Density | Abaxial | Rainfed | Season 1 | 4A | 59,477,918 | -0.09 | 5.30 |
| Stomatal Density | Abaxial | Rainfed | Season 1 | 5A | 593,822,668 | -0.12 | 11.30 |
| Stomatal Density | Abaxial | Rainfed | Season 1 | 7B | 722,137,906 | -0.12 | 8.55 |
| Stomatal Density | Adaxial | Rainfed | Season 1 | 1A | 139,877,951 | -0.13 | 5.48 |
| Stomatal Density | Adaxial | Rainfed | Season 1 | 4A | 660,553,574 | -0.10 | 3.78 |
| Stomatal Density | Adaxial | Rainfed | Season 1 | 5A | 597,245,434 | 0.21 | 11.20 |
| Stomatal Density | Adaxial | Rainfed | Season 1 | 6A | 103,685,883 | 0.13 | 6.64 |
| Stomatal Density | Abaxial | Irrigated | Season 1 | 2A | 462,910,177 | -0.12 | 4.34 |
| Stomatal Density | Abaxial | Irrigated | Season 1 | 4A | 683,161,804 | 0.28 | 14.55 |
| Stomatal Density | Abaxial | Irrigated | Season 1 | 6B | 24,575,379 | 0.17 | 7.48 |
| Stomatal Density | Abaxial | Irrigated | Season 1 | 7A | 258,084,246 | -0.12 | 3.64 |
| Stomatal Density | Abaxial | Irrigated | Season 1 | 7A | 500,966,888 | 0.12 | 5.52 |
| Stomatal Density | Abaxial | Irrigated | Season 1 | 7B | 691,332,873 | -0.08 | 3.20 |
| Stomatal Density | Adaxial | Irrigated | Season 1 | 1A | 575,406,410 | 0.15 | 3.70 |
| Stomatal Density | Adaxial | Irrigated | Season 1 | 1B | 650,483,640 | -0.10 | 3.06 |
| Stomatal Density | Adaxial | Irrigated | Season 1 | 3B | 851,566,773 | -0.11 | 3.53 |
| Stomatal Density | Adaxial | Irrigated | Season 1 | 5B | 680,160,246 | 0.12 | 4.19 |

| | | | | | | | |
|---|---|---|---|---|---|---|---|
| Stomatal Density | Adaxial | Irrigated | Season 1 | 7B | 108,872,880 | 0.09 | 3.02 |
| Stomatal Density | Abaxial | TOS1 | Season 2 | 1D | 495,213,341 | 0.19 | 5.24 |
| Stomatal Density | Abaxial | TOS1 | Season 2 | 6A | 456,737,110 | -0.15 | 4.83 |
| Stomatal Density | Abaxial | TOS2 | Season 2 | 2D | 83,018,229 | 0.32 | 9.43 |
| Stomatal Density | Abaxial | TOS2 | Season 2 | 4A | 662,285,005 | -0.15 | 6.87 |
| Stomatal Density | Abaxial | TOS2 | Season 2 | 6B | 601,676,496 | 0.15 | 4.03 |
| Stomatal Density | Adaxial | TOS2 | Season 2 | 3B | 28,249,993 | -0.25 | 5.56 |
| Stomatal Density | Adaxial | TOS2 | Season 2 | 5B | 681,423,027 | 0.15 | 5.29 |
| Stomata Conductance | Abaxial | Rainfed | Season 1 | 6A | 524,425,111 | 0.04 | 6.32 |
| Stomata Conductance | Abaxial | Rainfed | Season 1 | 6B | 141,648,696 | -0.03 | 7.24 |
| Stomata Conductance | Abaxial | Irrigated | Season 1 | 5B | 463,506,962 | 0.04 | 4.39 |
| Stomata Conductance | Abaxial | Irrigated | Season 1 | 6B | 631,016,005 | 0.03 | 5.00 |
| Stomata Conductance | Adaxial | TOS2 | Season 2 | 2B | 48,698,216 | -0.03 | 3.47 |
| Gsmax | Abaxial | Rainfed | Season 1 | 1A | 139,877,951 | -0.04 | 5.52 |
| Gsmax | Abaxial | Rainfed | Season 1 | 3B | 28,279,965 | -0.04 | 4.61 |
| Gsmax | Abaxial | Rainfed | Season 1 | 3B | 75,468,220 | -0.07 | 4.21 |
| Gsmax | Abaxial | Rainfed | Season 1 | 5A | 593,822,668 | -0.03 | 4.44 |

| | | | | | | | |
|---|---|---|---|---|---|---|---|
| Gsmax | Abaxial | Rainfed | Season 1 | 7B | 722,137,906 | -0.03 | 5.83 |
| Gsmax | Adaxial | Rainfed | Season 1 | 1A | 139,877,951 | -0.04 | 3.87 |
| Gsmax | Adaxial | Rainfed | Season 1 | 2B | 778,237,943 | -0.03 | 3.59 |
| Gsmax | Adaxial | Rainfed | Season 1 | 3B | 734,499,113 | 0.04 | 5.88 |
| Gsmax | Adaxial | Rainfed | Season 1 | 5A | 597,245,434 | 0.05 | 6.88 |
| Gsmax | Adaxial | Rainfed | Season 1 | 6A | 533,522,737 | 0.05 | 9.63 |
| Gsmax | Abaxial | Irrigated | Season 1 | 1A | 569,817,822 | 0.04 | 4.97 |
| Gsmax | Abaxial | Irrigated | Season 1 | 1B | 402,627,715 | -0.05 | 5.80 |
| Gsmax | Abaxial | Irrigated | Season 1 | 2B | 537,255,668 | -0.07 | 3.68 |
| Gsmax | Abaxial | Irrigated | Season 1 | 3B | 620,058,540 | 0.09 | 3.50 |
| Gsmax | Abaxial | Irrigated | Season 1 | 3B | 817,739,373 | -0.04 | 6.69 |
| Gsmax | Abaxial | Irrigated | Season 1 | 4A | 686,071,082 | 0.05 | 9.21 |
| Gsmax | Abaxial | Irrigated | Season 1 | 5B | 513,420,834 | -0.03 | 3.75 |
| Gsmax | Abaxial | Irrigated | Season 1 | 7B | 531,020,143 | 0.03 | 3.76 |
| Gsmax | Abaxial | Irrigated | Season 1 | 7B | 744,070,052 | 0.03 | 3.40 |
| Gsmax | Adaxial | Irrigated | Season 1 | 1B | 390,349,007 | -0.05 | 3.98 |
| Gsmax | Adaxial | Irrigated | Season 1 | 3B | 253,591,002 | -0.03 | 3.39 |
| Gsmax | Adaxial | Irrigated | Season 1 | 5A | 597,249,349 | 0.05 | 8.10 |
| Gsmax | Adaxial | Irrigated | Season 1 | 5B | 548,489,764 | -0.05 | 4.08 |

| | | | | | | | |
|---|---|---|---|---|---|---|---|
| Gsmax | Adaxial | Irrigated | Season 1 | 5D | 551,440,832 | -0.07 | 5.80 |
| Gsmax | Adaxial | Irrigated | Season 1 | 6A | 21,499,235 | 0.06 | 10.76 |
| Gsmax | Abaxial | TOS2 | Season 2 | 1D | 487,065,340 | 0.05 | 4.10 |
| Gsmax | Adaxial | TOS2 | Season 2 | 1A | 569,817,822 | 0.06 | 8.48 |
| Gsmax | Adaxial | TOS2 | Season 2 | 5B | 71,390,455 | 0.05 | 4.71 |
| Gs Efficiency | Abaxial | Rainfed | Season 1 | 6A | 524,424,694 | 0.04 | 6.57 |
| Gs Efficiency | Abaxial | Rainfed | Season 1 | 6B | 141,648,696 | -0.03 | 6.90 |
| Gs Efficiency | Abaxial | Irrigated | Season 1 | 1B | 643,977,170 | 0.04 | 2.62 |
| Gs Efficiency | Abaxial | Irrigated | Season 1 | 5B | 73,286,679 | -0.02 | 3.97 |
| Gs Efficiency | Abaxial | Irrigated | Season 1 | 5B | 463,506,962 | 0.04 | 4.87 |
| Gs Efficiency | Abaxial | Irrigated | Season 1 | 6B | 631,157,837 | 0.03 | 4.37 |
| Gs Efficiency | Abaxial | Irrigated | Season 1 | 7A | 582,463,366 | 0.02 | 2.66 |
| Gs Efficiency | Adaxial | TOS2 | Season 2 | 2B | 15,742,085 | 0.03 | 4.05 |
| Gs Efficiency | Adaxial | TOS2 | Season 2 | 6B | 585,226,952 | 0.03 | 3.28 |

**Table S8: The number of distinct putative QTL candidates across all trials by trait and chromosome.** The "." represents 0 and if the number is marked with a * then one of the QTL candidates was also detected in one other trial (*) or two other trials (**).

| **Chromosome** | $g_s$ | **GCW** | **GCL** | **SA** | **SD** | $g_{smax}$ | $g_{se}$ |
|---|---|---|---|---|---|---|---|
| **1A** | . | 5 | 2 | 3 | 3* | 4** | . |
| **1B** | . | 3 | 2 | 4 | 1 | 2 | 1 |
| **1D** | . | . | 1 | 1 | 1 | 1 | . |
| **2A** | . | . | 1 | . | 1 | . | . |
| **2B** | 1 | . | 7 | 3 | 2 | 2 | 1 |
| **2D** | . | . | . | . | 1 | . | . |
| **3A** | . | 2 | 2 | 1 | . | . | . |
| **3B** | . | 2 | 7* | 2 | 5* | 6 | . |
| **4A** | . | . | . | . | 4 | 1 | . |
| **4B** | . | . | 1 | . | . | . | . |
| **5A** | . | 1 | 1 | 1 | 2 | 3 | . |
| **5B** | 1 | . | 3 | 5 | 2 | 3 | 2 |
| **5D** | . | . | . | . | . | 1 | . |
| **6A** | 1 | . | 2 | 4 | 2 | 2 | 1 |
| **6B** | 2 | 5 | 5 | 4 | 2 | . | 3 |
| **7A** | . | 3 | . | 3 | 2 | . | 1 |
| **7B** | . | 1 | 3 | 4 | 3 | 3 | . |
| **7D** | . | . | . | 2 | . | . | . |
| **Total** | 5 | 22 | 37 | 37 | 31 | 28 | 9 |

**Table S9: The list of pleiotropic QTL candidate markers that were identified in more than one trial or trait.**

| Chromosome | Position | Season | Treatment | Surface | Trait |
|---|---:|---|---|---|---|
| 1A | 139,877,951 | Season 1 | Rainfed | Abaxial | SD |
| 1A | 139,877,951 | Season 1 | Rainfed | Adaxial | SD |
| 1A | 139,877,951 | Season 1 | Rainfed | Abaxial | gsmax |
| 1A | 139,877,951 | Season 1 | Rainfed | Adaxial | gsmax |
| 1A | 489,023,694 | Season 2 | TOS2 | Abaxial | GCW |
| 1A | 489,023,694 | Season 2 | TOS2 | Abaxial | SA |
| 1A | 569,817,822 | Season 1 | Irrigated | Abaxial | gsmax |
| 1A | 569,817,822 | Season 2 | TOS2 | Adaxial | gsmax |
| 3B | 28,249,993 | Season 1 | Rainfed | Abaxial | GCL |
| 3B | 28,249,993 | Season 2 | TOS1 | Abaxial | GCL |
| 3B | 28,249,993 | Season 2 | TOS1 | Abaxial | SA |
| 3B | 28,249,993 | Season 1 | Rainfed | Abaxial | SD |
| 3B | 28,249,993 | Season 2 | TOS2 | Adaxial | SD |
| 3B | 75,468,220 | Season 1 | Rainfed | Abaxial | SD |
| 3B | 75,468,220 | Season 1 | Rainfed | Abaxial | gsmax |
| 5A | 593,822,668 | Season 1 | Rainfed | Abaxial | SD |
| 5A | 593,822,668 | Season 1 | Rainfed | Abaxial | gsmax |
| 5A | 597,245,434 | Season 1 | Rainfed | Adaxial | SD |
| 5A | 597,245,434 | Season 1 | Rainfed | Adaxial | gsmax |
| 5B | 463,506,962 | Season 1 | Irrigated | Abaxial | gs |
| 5B | 463,506,962 | Season 1 | Irrigated | Abaxial | gse |
| 5B | 681,423,027 | Season 2 | TOS1 | Abaxial | GCL |
| 5B | 681,423,027 | Season 2 | TOS2 | Adaxial | SD |
| 6A | 524,424,694 | Season 1 | Rainfed | Adaxial | SA |
| 6A | 524,424,694 | Season 1 | Rainfed | Abaxial | gse |
| 6B | 141,648,696 | Season 1 | Rainfed | Abaxial | gs |
| 6B | 141,648,696 | Season 1 | Rainfed | Abaxial | gse |

| | | | | | |
|---|---|---|---|---|---|
| 6B | 631,016,005 | Season 1 | Irrigated | Abaxial | GCL |
| 6B | 631,016,005 | Season 1 | Irrigated | Abaxial | gs |
| 7B | 722,137,906 | Season 1 | Rainfed | Abaxial | SD |
| 7B | 722,137,906 | Season 1 | Rainfed | Abaxial | gsmax |

**Table S10: The number of putative QTLs within a 10 Mbp region by chromosome and trait.** A region was clustered using complete-linkage in hierarchical clustering such that no marker in a region would have distance of more than 10Mbp. The 18 bolded rows indicate a region that may contain pleiotropic QTLs for stomatal traits.

| Chromosome | Range of region (Mbp) | GCL | GCW | SA | SD | $g_s$ | $g_{smax}$ | $g_{se}$ |
|---|---|---|---|---|---|---|---|---|
| 1A | [8,10] | 1 | 0 | 0 | 0 | 0 | 0 | 0 |
| 1A | [15,17] | 0 | 0 | 1 | 0 | 0 | 0 | 0 |
| 1A | [17,19] | 0 | 0 | 1 | 0 | 0 | 0 | 0 |
| **1A** | **[139,140]** | **0** | **0** | **0** | **2** | **0** | **2** | **0** |
| **1A** | **[488,490]** | **0** | **1** | **1** | **0** | **0** | **0** | **0** |
| 1A | [547,549] | 0 | 1 | 0 | 0 | 0 | 0 | 0 |
| 1A | [550,551] | 0 | 1 | 0 | 0 | 0 | 0 | 0 |
| 1A | [560,562] | 0 | 1 | 0 | 0 | 0 | 0 | 0 |
| **1A** | **[569,571]** | **1** | **0** | **0** | **0** | **0** | **2** | **0** |
| **1A** | **[574,576]** | **0** | **1** | **0** | **1** | **0** | **0** | **0** |
| 1B | [323,324] | 0 | 0 | 1 | 0 | 0 | 0 | 0 |
| 1B | [371,372] | 1 | 0 | 0 | 0 | 0 | 0 | 0 |
| 1B | [389,391] | 0 | 0 | 0 | 0 | 0 | 1 | 0 |
| 1B | [402,403] | 0 | 0 | 0 | 0 | 0 | 1 | 0 |
| 1B | [426,428] | 0 | 0 | 1 | 0 | 0 | 0 | 0 |
| **1B** | **[555,556]** | **1** | **1** | **0** | **0** | **0** | **0** | **0** |
| 1B | [584,586] | 0 | 1 | 0 | 0 | 0 | 0 | 0 |
| 1B | [631,633] | 0 | 1 | 0 | 0 | 0 | 0 | 0 |
| 1B | [643,645] | 0 | 0 | 0 | 0 | 0 | 0 | 1 |
| 1B | [650,651] | 0 | 0 | 0 | 1 | 0 | 0 | 0 |
| 1B | [682,684] | 0 | 0 | 1 | 0 | 0 | 0 | 0 |
| 1B | [684,686] | 0 | 0 | 1 | 0 | 0 | 0 | 0 |
| 1D | [35,37] | 1 | 0 | 0 | 0 | 0 | 0 | 0 |
| 1D | [49,50] | 0 | 0 | 1 | 0 | 0 | 0 | 0 |

| | | | | | | | | |
|---|---|---|---|---|---|---|---|---|
| 1D | [487,488] | 0 | 0 | 0 | 0 | 0 | 1 | 0 |
| 1D | [495,497] | 0 | 0 | 0 | 1 | 0 | 0 | 0 |
| 2A | [462,464] | 0 | 0 | 0 | 1 | 0 | 0 | 0 |
| 2A | [766,768] | 1 | 0 | 0 | 0 | 0 | 0 | 0 |
| 2B | [15,16] | 0 | 0 | 0 | 0 | 0 | 0 | 1 |
| 2B | [33,35] | 1 | 0 | 0 | 0 | 0 | 0 | 0 |
| 2B | [48,50] | 0 | 0 | 0 | 0 | 1 | 0 | 0 |
| 2B | [57,59] | 0 | 0 | 1 | 0 | 0 | 0 | 0 |
| 2B | [84,85] | 0 | 0 | 1 | 0 | 0 | 0 | 0 |
| 2B | [125,127] | 1 | 0 | 0 | 0 | 0 | 0 | 0 |
| 2B | [148,150] | 0 | 0 | 1 | 0 | 0 | 0 | 0 |
| 2B | [161,162] | 1 | 0 | 0 | 0 | 0 | 0 | 0 |
| 2B | [203,205] | 1 | 0 | 0 | 0 | 0 | 0 | 0 |
| 2B | [257,258] | 1 | 0 | 0 | 0 | 0 | 0 | 0 |
| 2B | [462,463] | 1 | 0 | 0 | 0 | 0 | 0 | 0 |
| 2B | [537,538] | 0 | 0 | 0 | 0 | 0 | 1 | 0 |
| 2B | [646,647] | 0 | 0 | 0 | 1 | 0 | 0 | 0 |
| 2B | [679,681] | 0 | 0 | 0 | 1 | 0 | 0 | 0 |
| 2B | [687,689] | 1 | 0 | 0 | 0 | 0 | 0 | 0 |
| 2B | [778,779] | 0 | 0 | 0 | 0 | 0 | 1 | 0 |
| 2D | [83,84] | 0 | 0 | 0 | 1 | 0 | 0 | 0 |
| 3A | [58,59] | 0 | 1 | 0 | 0 | 0 | 0 | 0 |
| 3A | [558,560] | 1 | 0 | 0 | 0 | 0 | 0 | 0 |
| 3A | [561,563] | 0 | 1 | 0 | 0 | 0 | 0 | 0 |
| 3A | [648,650] | 0 | 0 | 1 | 0 | 0 | 0 | 0 |
| 3A | [740,742] | 1 | 0 | 0 | 0 | 0 | 0 | 0 |
| 3B | [24,26] | 1 | 0 | 0 | 0 | 0 | 0 | 0 |
| **3B** | **[28,29]** | **2** | **0** | **1** | **2** | **0** | **1** | **0** |
| **3B** | **[75,77]** | **0** | **0** | **0** | **1** | **0** | **1** | **0** |

| | | | | | | | | |
|---|---|---|---|---|---|---|---|---|
| 3B | [117,118] | 1 | 0 | 0 | 0 | 0 | 0 | 0 |
| 3B | [252,254] | 0 | 0 | 0 | 0 | 0 | 1 | 0 |
| 3B | [536,538] | 0 | 0 | 0 | 1 | 0 | 0 | 0 |
| 3B | [589,591] | 0 | 1 | 0 | 0 | 0 | 0 | 0 |
| 3B | [607,608] | 0 | 0 | 1 | 0 | 0 | 0 | 0 |
| 3B | [619,621] | 0 | 0 | 0 | 0 | 0 | 1 | 0 |
| 3B | [711,713] | 1 | 0 | 0 | 0 | 0 | 0 | 0 |
| 3B | [734,735] | 0 | 0 | 0 | 0 | 0 | 1 | 0 |
| 3B | [812,814] | 0 | 1 | 0 | 0 | 0 | 0 | 0 |
| 3B | [813,814] | 2 | 0 | 0 | 0 | 0 | 0 | 0 |
| 3B | [817,818] | 0 | 0 | 0 | 0 | 0 | 1 | 0 |
| 3B | [851,852] | 0 | 0 | 0 | 1 | 0 | 0 | 0 |
| 4A | [59,60] | 0 | 0 | 0 | 1 | 0 | 0 | 0 |
| 4A | [660,662] | 0 | 0 | 0 | 1 | 0 | 0 | 0 |
| 4A | [661,663] | 0 | 0 | 0 | 1 | 0 | 0 | 0 |
| 4A | [683,684] | 0 | 0 | 0 | 1 | 0 | 0 | 0 |
| 4A | [685,687] | 0 | 0 | 0 | 0 | 0 | 1 | 0 |
| 4B | [525,527] | 1 | 0 | 0 | 0 | 0 | 0 | 0 |
| 5A | [438,440] | 1 | 0 | 0 | 0 | 0 | 0 | 0 |
| 5A | [591,592] | 0 | 0 | 1 | 0 | 0 | 0 | 0 |
| 5A | [592,594] | 0 | 1 | 0 | 0 | 0 | 0 | 0 |
| **5A** | **[593,595]** | **0** | **0** | **0** | **1** | **0** | **1** | **0** |
| **5A** | **[596,598]** | **0** | **0** | **0** | **1** | **0** | **2** | **0** |
| 5B | [63,64] | 0 | 0 | 1 | 0 | 0 | 0 | 0 |
| 5B | [71,73] | 0 | 0 | 0 | 0 | 0 | 1 | 0 |
| 5B | [73,74] | 0 | 0 | 0 | 0 | 0 | 0 | 1 |
| 5B | [76,78] | 0 | 0 | 1 | 0 | 0 | 0 | 0 |
| 5B | [352,354] | 0 | 0 | 1 | 0 | 0 | 0 | 0 |
| **5B** | **[463,464]** | **0** | **0** | **0** | **0** | **1** | **0** | **1** |

| | | | | | | | | |
|---|---|---|---|---|---|---|---|---|
| 5B | [513,514] | 0 | 0 | 0 | 0 | 0 | 1 | 0 |
| 5B | [517,519] | 1 | 0 | 0 | 0 | 0 | 0 | 0 |
| 5B | [536,537] | 1 | 0 | 0 | 0 | 0 | 0 | 0 |
| 5B | [548,549] | 0 | 0 | 0 | 0 | 0 | 1 | 0 |
| 5B | [597,598] | 0 | 0 | 1 | 0 | 0 | 0 | 0 |
| 5B | [679,681] | 0 | 0 | 0 | 1 | 0 | 0 | 0 |
| **5B** | **[680,682]** | **1** | **0** | **0** | **1** | **0** | **0** | **0** |
| 5B | [713,714] | 0 | 0 | 1 | 0 | 0 | 0 | 0 |
| 5D | [550,552] | 0 | 0 | 0 | 0 | 0 | 1 | 0 |
| 6A | [21,22] | 0 | 0 | 0 | 0 | 0 | 1 | 0 |
| 6A | [103,104] | 0 | 0 | 0 | 1 | 0 | 0 | 0 |
| 6A | [323,324] | 1 | 0 | 0 | 0 | 0 | 0 | 0 |
| 6A | [392,394] | 0 | 0 | 1 | 0 | 0 | 0 | 0 |
| 6A | [443,445] | 1 | 0 | 0 | 0 | 0 | 0 | 0 |
| 6A | [456,458] | 0 | 0 | 0 | 1 | 0 | 0 | 0 |
| **6A** | **[524,525]** | **0** | **0** | **1** | **0** | **1** | **0** | **1** |
| 6A | [533,535] | 0 | 0 | 0 | 0 | 0 | 1 | 0 |
| 6A | [555,557] | 0 | 0 | 1 | 0 | 0 | 0 | 0 |
| 6A | [616,617] | 0 | 0 | 1 | 0 | 0 | 0 | 0 |
| 6B | [16,18] | 0 | 0 | 1 | 0 | 0 | 0 | 0 |
| 6B | [24,26] | 0 | 0 | 0 | 1 | 0 | 0 | 0 |
| 6B | [106,108] | 1 | 0 | 0 | 0 | 0 | 0 | 0 |
| 6B | [129,131] | 0 | 1 | 0 | 0 | 0 | 0 | 0 |
| 6B | [133,134] | 0 | 1 | 0 | 0 | 0 | 0 | 0 |
| **6B** | **[140,142]** | **0** | **0** | **0** | **0** | **1** | **0** | **1** |
| 6B | [163,164] | 0 | 0 | 1 | 0 | 0 | 0 | 0 |
| 6B | [199,200] | 1 | 0 | 0 | 0 | 0 | 0 | 0 |
| 6B | [202,204] | 0 | 1 | 0 | 0 | 0 | 0 | 0 |
| 6B | [304,306] | 1 | 0 | 0 | 0 | 0 | 0 | 0 |

| | | | | | | | | |
|---|---|---|---|---|---|---|---|---|
| 6B | [478,480] | 0 | 1 | 0 | 0 | 0 | 0 | 0 |
| 6B | [534,536] | 0 | 1 | 0 | 0 | 0 | 0 | 0 |
| 6B | [584,586] | 0 | 0 | 0 | 0 | 0 | 0 | 1 |
| 6B | [601,602] | 0 | 0 | 0 | 1 | 0 | 0 | 0 |
| 6B | [607,609] | 0 | 0 | 1 | 0 | 0 | 0 | 0 |
| **6B** | **[631,632]** | **1** | **0** | **0** | **0** | **1** | **0** | **1** |
| 6B | [632,634] | 1 | 0 | 0 | 0 | 0 | 0 | 0 |
| 6B | [726,728] | 0 | 0 | 1 | 0 | 0 | 0 | 0 |
| 7A | [84,86] | 0 | 0 | 1 | 0 | 0 | 0 | 0 |
| **7A** | **[105,107]** | **0** | **1** | **1** | **0** | **0** | **0** | **0** |
| 7A | [137,138] | 0 | 1 | 0 | 0 | 0 | 0 | 0 |
| 7A | [257,259] | 0 | 0 | 0 | 1 | 0 | 0 | 0 |
| 7A | [500,502] | 0 | 0 | 0 | 1 | 0 | 0 | 0 |
| 7A | [582,583] | 0 | 0 | 0 | 0 | 0 | 0 | 1 |
| 7A | [677,678] | 0 | 1 | 0 | 0 | 0 | 0 | 0 |
| 7A | [696,698] | 0 | 0 | 1 | 0 | 0 | 0 | 0 |
| 7B | [29,31] | 0 | 0 | 1 | 0 | 0 | 0 | 0 |
| 7B | [71,72] | 1 | 0 | 0 | 0 | 0 | 0 | 0 |
| 7B | [90,92] | 0 | 1 | 0 | 0 | 0 | 0 | 0 |
| **7B** | **[108,110]** | **0** | **0** | **1** | **1** | **0** | **0** | **0** |
| 7B | [153,155] | 1 | 0 | 0 | 0 | 0 | 0 | 0 |
| 7B | [210,211] | 0 | 0 | 1 | 0 | 0 | 0 | 0 |
| 7B | [530,532] | 0 | 0 | 0 | 0 | 0 | 1 | 0 |
| 7B | [690,692] | 0 | 0 | 0 | 1 | 0 | 0 | 0 |
| 7B | [717,718] | 0 | 0 | 1 | 0 | 0 | 0 | 0 |
| **7B** | **[721,723]** | **0** | **0** | **0** | **1** | **0** | **1** | **0** |
| **7B** | **[743,745]** | **1** | **0** | **0** | **0** | **0** | **1** | **0** |
| 7D | [56,57] | 0 | 0 | 1 | 0 | 0 | 0 | 0 |
| 7D | [61,62] | 0 | 0 | 1 | 0 | 0 | 0 | 0 |

**Table S11: The number of QTLs reported in literature by trait and chromosome.** The last column shows the number of putative QTLs we found for the same trait and chromosome.

| Trait | Chromosome | # Reported QTLs | Source | # Putative QTLs found in our study |
|---|---|---|---|---|
| SA | 1A | 4 | Ahmed et al. 2021, Chaplin et al. 2025 | 3 |
| SA | 1B | 10 | Shahinnia et al. 2016, Liu et al. 2025, Ahmed et al. 2021, Chaplin et al. 2025 | 4 |
| SA | 2A | 3 | Ahmed et al. 2021 | 0 |
| SA | 2B | 5 | Liu et al. 2025, Ahmed et al. 2021, Chaplin et al. 2025 | 3 |
| SA | 2D | 3 | Ahmed et al. 2021, Chaplin et al. 2025 | 0 |
| SA | 3A | 2 | Chaplin et al. 2025 | 1 |
| SA | 3B | 7 | Ahmed et al. 2021, Chaplin et al. 2025 | 2 |
| SA | 3D | 3 | Ahmed et al. 2021, Chaplin et al. 2025 | 0 |
| SA | 4A | 1 | Shahinnia et al. 2016 | 0 |
| SA | 4B | 10 | Shahinnia et al. 2016, Liu et al. 2025, Ahmed et al. 2021, Chaplin et al. 2025 | 0 |
| SA | 5A | 5 | Shahinnia et al. 2016, Liu et al. 2025, Ahmed et al. 2021, Chaplin et al. 2025 | 1 |
| SA | 5B | 22 | Shahinnia et al. 2016, Liu et al. 2025, Ahmed et al. 2021, Chaplin et al. 2025 | 5 |
| SA | 5D | 3 | Shahinnia et al. 2016, Liu et al. 2025 | 0 |
| SA | 6B | 6 | Ahmed et al. 2021, Chaplin et al. 2025 | 4 |

| | | | | |
|---|---|---|---|---|
| SA | 7A | 10 | Ahmed et al. 2021, Chaplin et al. 2025 | 3 |
| SA | 7B | 2 | Chaplin et al. 2025 | 4 |
| SA | 7D | 2 | Chaplin et al. 2025 | 2 |
| SD | 1A | 6 | Liu et al. 2025, Ahmed et al. 2021, Chaplin et al. 2025 | 3 |
| SD | 1B | 3 | Ahmed et al. 2021, Chaplin et al. 2025 | 1 |
| SD | 1D | 2 | Ahmed et al. 2021 | 1 |
| SD | 2A | 7 | Ahmed et al. 2021, Chaplin et al. 2025 | 1 |
| SD | 2B | 2 | Ahmed et al. 2021, Chaplin et al. 2025 | 2 |
| SD | 2D | 1 | Ahmed et al. 2021 | 1 |
| SD | 3A | 7 | Shahinnia et al. 2016, Liu et al. 2025, Ahmed et al. 2021 | 0 |
| SD | 3B | 8 | Ahmed et al. 2021 | 5 |
| SD | 4A | 6 | Shahinnia et al. 2016, Ahmed et al. 2021 | 4 |
| SD | 4B | 8 | Shahinnia et al. 2016, Liu et al. 2025, Ahmed et al. 2021 | 0 |
| SD | 5A | 13 | Shahinnia et al. 2016, Liu et al. 2025, Ahmed et al. 2021, Chaplin et al. 2025 | 2 |
| SD | 5B | 4 | Shahinnia et al. 2016, Liu et al. 2025, Ahmed et al. 2021, Chaplin et al. 2025 | 2 |
| SD | 5D | 1 | Chaplin et al. 2025 | 0 |
| SD | 6A | 1 | Chaplin et al. 2025 | 2 |
| SD | 6B | 7 | Ahmed et al. 2021 | 2 |
| SD | 6D | 2 | Ahmed et al. 2021 | 0 |

| | | | | |
|---|---|---|---|---|
| SD | 7A | 10 | Shahinnia et al. 2016, Liu et al. 2025, Ahmed et al. 2021 | 2 |
| SD | 7B | 8 | Ahmed et al. 2021, Chaplin et al. 2025 | 3 |
| gs | 1A | 1 | Chaplin et al. 2025 | 0 |
| gs | 2A | 1 | Chaplin et al. 2025 | 0 |
| gs | 2B | 2 | Wang et al. 2015, Chaplin et al. 2025 | 1 |
| gs | 2D | 1 | Wang et al. 2015 | 0 |
| gs | 3A | 1 | Chaplin et al. 2025 | 0 |
| gs | 3D | 1 | Chaplin et al. 2025 | 0 |
| gs | 4A | 1 | Wang et al. 2015 | 0 |
| gs | 6D | 1 | Wang et al. 2015 | 0 |
| gs | 7A | 1 | Chaplin et al. 2025 | 0 |
| gs | 7B | 2 | Wang et al. 2015 | 0 |
| gse | 1A | 1 | Chaplin et al. 2025 | 0 |
| gse | 3A | 1 | Chaplin et al. 2025 | 0 |
| gse | 4A | 1 | Chaplin et al. 2025 | 0 |
| gse | 6D | 1 | Chaplin et al. 2025 | 0 |
| gsmax | 1A | 2 | Chaplin et al. 2025 | 4 |
| gsmax | 1B | 1 | Chaplin et al. 2025 | 2 |
| gsmax | 2B | 2 | Chaplin et al. 2025 | 2 |
| gsmax | 3B | 1 | Chaplin et al. 2025 | 6 |
| gsmax | 4B | 1 | Chaplin et al. 2025 | 0 |
| gsmax | 5B | 1 | Chaplin et al. 2025 | 3 |
| gsmax | 6A | 1 | Chaplin et al. 2025 | 2 |
| gsmax | 7A | 1 | Chaplin et al. 2025 | 0 |
| gsmax | 7B | 5 | Chaplin et al. 2025 | 3 |

| | | | | |
|---|---|---|---|---|
| GCL | 1A | 1 | Shahinnia et al. 2016 | 2 |
| GCL | 1B | 6 | Shahinnia et al. 2016, Liu et al. 2025, Chaplin et al. 2025 | 2 |
| GCL | 2B | 2 | Chaplin et al. 2025 | 7 |
| GCL | 3A | 2 | Chaplin et al. 2025 | 2 |
| GCL | 3B | 2 | Shahinnia et al. 2016, Chaplin et al. 2025 | 7 |
| GCL | 4B | 10 | Shahinnia et al. 2016, Liu et al. 2025, Chaplin et al. 2025 | 1 |
| GCL | 5A | 4 | Chaplin et al. 2025 | 1 |
| GCL | 5B | 8 | Liu et al. 2025, Chaplin et al. 2025 | 3 |
| GCL | 6A | 1 | Chaplin et al. 2025 | 2 |
| GCL | 6B | 2 | Chaplin et al. 2025 | 5 |
| GCL | 7A | 4 | Shahinnia et al. 2016, Chaplin et al. 2025 | 0 |
| GCL | 7B | 3 | Chaplin et al. 2025 | 3 |
| GCL | 7D | 6 | Shahinnia et al. 2016, Chaplin et al. 2025 | 0 |
| GCW | 1A | 3 | Chaplin et al. 2025 | 5 |
| GCW | 1B | 3 | Chaplin et al. 2025 | 3 |
| GCW | 2A | 1 | Liu et al. 2025 | 0 |
| GCW | 2B | 4 | Liu et al. 2025, Chaplin et al. 2025 | 0 |
| GCW | 2D | 1 | Chaplin et al. 2025 | 0 |
| GCW | 3B | 2 | Chaplin et al. 2025 | 2 |
| GCW | 4A | 1 | Chaplin et al. 2025 | 0 |
| GCW | 5B | 10 | Liu et al. 2025, Chaplin et al. 2025 | 0 |

| | | | | |
|---|---|---|---|---|
| GCW | 5D | 3 | Liu et al. 2025, Chaplin et al. 2025 | 0 |
| GCW | 6B | 5 | Chaplin et al. 2025 | 5 |
| GCW | 7A | 7 | Liu et al. 2025, Chaplin et al. 2025 | 3 |
| GCW | 7B | 4 | Chaplin et al. 2025 | 1 |
| GCW | 7D | 1 | Chaplin et al. 2025 | 0 |

**Table S12: A list of potential pleiotropic or stable QTL candidates that were found within a 10 Mbp region (as determined by hierarchical clustering based on complete linkage of marker position) in more than one trial or trait.** The source column indicates whether the QTL candidate was found in the Chaplin et al. (2025) (G) or this study (H). The size column shows the effect size. The row color alternate between white and grey to easily see markers in the same range. The rows are bolded if there are more than one QTL candidate for the same trait and surface within the same range.

| Chromosome | Range of region (Mbp) | Position (bp) | Size | Source | Treatment | Surface | Trait | Season |
|---|---|---|---|---|---|---|---|---|
| 1A | [15,17] | 15,656,623 | 0.0749 | G | TOS1 | Abaxial | gsmax | S1 |
| 1A | [15,17] | 15,675,703 | -73.9856 | H | TOS1 | Adaxial | SA | S2 |
| 1A | [139,140] | 139,877,951 | -0.1260 | H | Rainfed | Abaxial | SD | S1 |
| 1A | [139,140] | 139,877,951 | -0.1285 | H | Rainfed | Adaxial | SD | S1 |
| 1A | [139,140] | 139,877,951 | -0.0352 | H | Rainfed | Abaxial | gsmax | S1 |
| 1A | [139,140] | 139,877,951 | -0.0359 | H | Rainfed | Adaxial | gsmax | S1 |
| 1A | [376,378] | 376,842,796 | 2.3820 | G | TOS1 | Adaxial | GCW | S1 |
| 1A | [376,378] | 376,842,796 | 113.4830 | G | TOS1 | Adaxial | SA | S1 |
| 1A | [376,378] | 377,335,032 | 0.2674 | G | TOS1 | Adaxial | SD | S1 |
| 1A | [488,490] | 489,023,694 | 2.2648 | H | TOS2 | Abaxial | GCW | S2 |
| 1A | [488,490] | 489,023,694 | 173.7842 | H | TOS2 | Abaxial | SA | S2 |
| 1A | [569,571] | 569,817,822 | 0.0353 | H | Irrigated | Abaxial | gsmax | S1 |
| 1A | [569,571] | 569,817,822 | 0.0618 | H | TOS2 | Adaxial | gsmax | S2 |

| | | | | | | | | |
|---|---|---|---|---|---|---|---|---|
| 1A | [569,571] | 569,847,820 | 1.0181 | H | TOS2 | Abaxial | GCL | S2 |
| **1A** | **[574,576]** | **574,778,628** | **0.1278** | **G** | **TOS1** | **Adaxial** | **SD** | **S1** |
| **1A** | **[574,576]** | **575,406,410** | **0.1526** | **H** | **Irrigated** | **Adaxial** | **SD** | **S1** |
| **1A** | **[574,576]** | **575,536,483** | **0.9697** | **G** | **TOS2** | **Abaxial** | **GCW** | **S1** |
| 1A | [574,576] | 575,536,483 | 0.0578 | G | TOS1 | Adaxial | **gsmax** | S1 |
| **1A** | **[574,576]** | **575,737,813** | **0.8410** | **H** | **TOS2** | **Abaxial** | **GCW** | **S2** |
| **1B** | **[555,556]** | **555,411,043** | **-56.8919** | **G** | **TOS2** | **Abaxial** | **SA** | **S1** |
| **1B** | **[555,556]** | **555,412,597** | **66.3915** | **G** | **TOS1** | **Abaxial** | **SA** | **S1** |
| 1B | [555,556] | 555,412,597 | 72.2615 | G | TOS1 | Adaxial | SA | S1 |
| 1B | [555,556] | 555,625,561 | 0.6743 | H | Rainfed | Adaxial | GCW | S1 |
| 1B | [555,556] | 555,625,935 | 0.9916 | H | Rainfed | Adaxial | GCL | S1 |
| 2B | [462,463] | 462,199,334 | -2.1604 | G | TOS1 | Abaxial | GCW | S2 |
| 2B | [462,463] | 462,771,824 | -1.5748 | H | TOS2 | Adaxial | GCL | S2 |
| **2B** | **[687,689]** | **688,831,778** | **1.4355** | **G** | **TOS1** | **Abaxial** | **GCL** | **S1** |
| **2B** | **[687,689]** | **688,832,053** | **1.0334** | **H** | **Irrigated** | **Abaxial** | **GCL** | **S1** |
| **3A** | **[648,650]** | **648,742,426** | **-46.3013** | **G** | **TOS1** | **Abaxial** | **SA** | **S1** |
| **3A** | **[648,650]** | **648,742,426** | **-80.4846** | **G** | **TOS2** | **Abaxial** | **SA** | **S1** |
| **3A** | **[648,650]** | **649,063,085** | **-41.5156** | **H** | **TOS2** | **Abaxial** | **SA** | **S2** |
| **3A** | **[740,742]** | **741,357,936** | **-1.2963** | **H** | **Irrigated** | **Abaxial** | **GCL** | **S1** |

| | | | | | | | | |
|---|---|---|---|---|---|---|---|---|
| **3A** | **[740,742]** | **741,360,511** | **-1.5800** | **G** | **TOS2** | **Abaxial** | **GCL** | **S1** |
| 3A | [743,744] | 743,381,074 | 0.0187 | G | TOS1 | Abaxial | gs | S1 |
| 3A | [743,744] | 743,381,074 | 0.0152 | G | TOS1 | Abaxial | gse | S1 |
| **3B** | **[28,29]** | **28,249,993** | **0.9057** | **H** | **Rainfed** | **Abaxial** | **GCL** | **S1** |
| **3B** | **[28,29]** | **28,249,993** | **1.8077** | **H** | **TOS1** | **Abaxial** | **GCL** | **S2** |
| 3B | [28,29] | 28,249,993 | 65.8142 | H | TOS1 | Abaxial | SA | S2 |
| 3B | [28,29] | 28,249,993 | -0.1708 | H | Rainfed | Abaxial | SD | S1 |
| 3B | [28,29] | 28,249,993 | -0.2514 | H | TOS2 | Adaxial | SD | S2 |
| 3B | [28,29] | 28,279,965 | -0.0392 | H | Rainfed | Abaxial | gsmax | S1 |
| 3B | [75,77] | 75,468,220 | -0.2596 | H | Rainfed | Abaxial | SD | S1 |
| 3B | [75,77] | 75,468,220 | -0.0722 | H | Rainfed | Abaxial | gsmax | S1 |
| 3B | [813,814] | 813,286,435 | 0.6309 | H | Rainfed | Abaxial | GCL | S1 |
| 3B | [813,814] | 813,648,022 | 0.8795 | H | Irrigated | Adaxial | GCL | S1 |
| 5A | [591,593] | 591,952,394 | -0.9558 | G | TOS1 | Abaxial | GCL | S1 |
| 5A | [591,593] | 592,188,836 | 0.1021 | G | TOS1 | Adaxial | SD | S1 |
| 5A | [592,594] | 592,814,280 | -0.1099 | G | TOS1 | Abaxial | SD | S1 |
| 5A | [592,594] | 593,322,475 | 0.5555 | H | TOS1 | Adaxial | GCW | S2 |
| 5A | [593,595] | 593,822,668 | -0.1209 | H | Rainfed | Abaxial | SD | S1 |
| 5A | [593,595] | 593,822,668 | -0.0273 | H | Rainfed | Abaxial | gsmax | S1 |

| | | | | | | | | |
|---|---|---|---|---|---|---|---|---|
| 5A | [596,598] | 597,245,434 | 0.2052 | H | Rainfed | Adaxial | SD | S1 |
| **5A** | **[596,598]** | **597,245,434** | **0.0528** | **H** | **Rainfed** | **Adaxial** | **gsmax** | **S1** |
| **5A** | **[596,598]** | **597,249,349** | **0.0510** | **H** | **Irrigated** | **Adaxial** | **gsmax** | **S1** |
| 5A | [665,667] | 665,150,454 | 1.3960 | G | TOS1 | Adaxial | GCL | S1 |
| 5A | [665,667] | 665,150,454 | 76.7119 | G | TOS1 | Adaxial | SA | S1 |
| 5B | [406,408] | 406,738,159 | 1.8485 | G | TOS2 | Abaxial | GCL | S2 |
| 5B | [406,408] | 406,738,159 | 53.8168 | G | TOS2 | Abaxial | SA | S2 |
| 5B | [406,408] | 407,109,507 | -1.8399 | G | TOS2 | Adaxial | GCL | S2 |
| 5B | [463,464] | 463,506,962 | 0.0412 | H | Irrigated | Abaxial | gs | S1 |
| 5B | [463,464] | 463,506,962 | 0.0374 | H | Irrigated | Abaxial | gse | S1 |
| 5B | [548,549] | 548,489,764 | -0.0489 | H | Irrigated | Adaxial | gsmax | S1 |
| 5B | [548,549] | 548,494,031 | -0.7950 | G | TOS1 | Abaxial | GCL | S1 |
| 5B | [680,682] | 681,423,027 | -0.6255 | H | TOS1 | Abaxial | GCL | S2 |
| **5B** | **[680,682]** | **681,423,027** | **0.1537** | **H** | **TOS2** | **Adaxial** | **SD** | **S2** |
| **5B** | **[680,682]** | **681,423,212** | **0.1638** | **G** | **TOS2** | **Adaxial** | **SD** | **S1** |
| 5B | [680,682] | 681,423,212 | 0.0541 | G | TOS2 | Adaxial | gsmax | S1 |
| 5B | [713,714] | 713,283,368 | 40.2831 | H | Rainfed | Abaxial | SA | S1 |
| 5B | [713,714] | 713,283,368 | 39.3817 | G | TOS2 | Adaxial | SA | S1 |
| 5B | [713,714] | 713,297,219 | 0.4624 | G | TOS2 | Adaxial | GCW | S1 |

| | | | | | | | | |
|---|---|---|---|---|---|---|---|---|
| 5D | [552,553] | 552,286,738 | 0.5054 | G | TOS2 | Adaxial | GCW | S1 |
| 5D | [552,553] | 552,685,132 | -0.1722 | G | TOS2 | Adaxial | SD | S1 |
| 6A | [524,525] | 524,424,694 | -79.7076 | H | Rainfed | Adaxial | SA | S1 |
| 6A | [524,525] | 524,424,694 | 0.0390 | H | Rainfed | Abaxial | gse | S1 |
| 6A | [524,525] | 524,425,111 | 0.0391 | H | Rainfed | Abaxial | gs | S1 |
| 6A | [581,582] | 581,018,698 | 0.1905 | G | TOS1 | Abaxial | SD | S1 |
| 6A | [581,582] | 581,018,698 | 0.0674 | G | TOS1 | Abaxial | gsmax | S1 |
| **6B** | **[23,25]** | **23,484,434** | **-0.5318** | **G** | **TOS2** | **Abaxial** | **GCW** | **S1** |
| **6B** | **[23,25]** | **23,602,849** | **-0.9566** | **G** | **TOS1** | **Abaxial** | **GCW** | **S1** |
| 6B | [140,142] | 141,648,696 | -0.0299 | H | Rainfed | Abaxial | gs | S1 |
| 6B | [140,142] | 141,648,696 | -0.0284 | H | Rainfed | Abaxial | gse | S1 |
| 6B | [482,483] | 482,341,150 | -0.9228 | G | TOS1 | Adaxial | GCL | S1 |
| 6B | [482,483] | 482,975,955 | -0.9113 | G | TOS1 | Adaxial | GCW | S1 |
| 6B | [482,483] | 482,975,955 | -59.3518 | G | TOS1 | Adaxial | SA | S1 |
| 6B | [631,632] | 631,016,005 | -1.3287 | H | Irrigated | Abaxial | GCL | S1 |
| 6B | [631,632] | 631,016,005 | 0.0323 | H | Irrigated | Abaxial | gs | S1 |
| 6B | [631,632] | 631,157,837 | 0.0255 | H | Irrigated | Abaxial | gse | S1 |
| 7A | [105,107] | 105,813,606 | -0.8099 | H | TOS1 | Abaxial | GCW | S2 |
| 7A | [105,107] | 105,815,982 | -32.9286 | H | TOS1 | Abaxial | SA | S2 |

| | | | | | | | | |
|---|---|---|---|---|---|---|---|---|
| 7A | [613,615] | 614,132,821 | 1.1340 | G | TOS1 | Adaxial | GCW | S1 |
| 7A | [613,615] | 614,132,821 | 125.5968 | G | TOS1 | Adaxial | SA | S1 |
| **7A** | **[677,678]** | **677,126,838** | **-0.7621** | **G** | **TOS1** | **Abaxial** | **GCW** | **S1** |
| **7A** | **[677,678]** | **677,127,635** | **-0.8015** | **H** | **Rainfed** | **Abaxial** | **GCW** | **S1** |
| **7B** | **[62,63]** | **62,016,354** | **-0.0551** | **G** | **TOS1** | **Adaxial** | **gsmax** | **S1** |
| **7B** | **[62,63]** | **62,057,627** | **-0.0632** | **G** | **TOS2** | **Adaxial** | **gsmax** | **S1** |
| 7B | [108,110] | 108,638,412 | -58.2851 | H | Irrigated | Adaxial | SA | S1 |
| 7B | [108,110] | 108,872,880 | 0.0938 | H | Irrigated | Adaxial | SD | S1 |
| 7B | [457,458] | 457,347,337 | -1.2474 | G | TOS2 | Adaxial | GCL | S1 |
| 7B | [457,458] | 457,582,679 | -44.2490 | G | TOS1 | Adaxial | SA | S1 |
| 7B | [526,527] | 526,910,362 | 0.1994 | G | TOS1 | Adaxial | SD | S1 |
| 7B | [526,527] | 526,910,362 | 0.0470 | G | TOS1 | Adaxial | gsmax | S1 |
| 7B | [526,527] | 526,913,571 | 0.1268 | G | TOS2 | Abaxial | SD | S1 |
| 7B | [547,548] | 547,215,666 | 0.1693 | G | TOS1 | Abaxial | SD | S1 |
| 7B | [547,548] | 547,215,666 | 0.0591 | G | TOS1 | Abaxial | gsmax | S1 |
| 7B | [721,723] | 722,137,906 | -0.1158 | H | Rainfed | Abaxial | SD | S1 |
| 7B | [721,723] | 722,137,906 | -0.0326 | H | Rainfed | Abaxial | gsmax | S1 |
| 7B | [743,745] | 744,070,052 | 0.0328 | H | Irrigated | Abaxial | gsmax | S1 |
| 7B | [743,745] | 744,071,081 | -1.5554 | H | Irrigated | Adaxial | GCL | S1 |

| | | | | | | | | |
|---|---|---|---|---|---|---|---|---|
| 7D | [6,8] | 6,846,653 | 2.2379 | G | TOS2 | Adaxial | GCL | S2 |
| 7D | [6,8] | 6,846,653 | 86.2714 | G | TOS2 | Abaxial | SA | S2 |
| **7D** | **[56,57]** | **56,579,964** | **0.6862** | **G** | **TOS1** | **Abaxial** | **GCL** | **S1** |
| 7D | [56,57] | 56,579,964 | 0.7833 | G | TOS2 | Abaxial | GCW | S1 |
| **7D** | **[56,57]** | **56,638,565** | **56.3652** | **H** | **Irrigated** | **Abaxial** | **SA** | **S1** |
| **7D** | **[56,57]** | **56,638,565** | **1.0882** | **G** | **TOS2** | **Abaxial** | **GCL** | **S1** |
| 7D | [56,57] | 56,638,565 | 1.1383 | G | TOS2 | Adaxial | GCL | S1 |
| **7D** | **[56,57]** | **56,960,435** | **64.3205** | **G** | **TOS2** | **Abaxial** | **SA** | **S1** |
